# Neuromodulation by Monoamines is a Bilaterian Innovation

**DOI:** 10.1101/2022.08.01.501419

**Authors:** Matthew Goulty, Gaelle Botton-Amiot, Ezio Rosato, Simon Sprecher, Roberto Feuda

## Abstract

Monoamines like serotonin, dopamine, and adrenaline/noradrenaline (epinephrine/ norepinephrine) act as neuromodulators that tune the response of the nervous system to the environment with predictable advantages for fitness. For instance, monoamines influence action selection depending on the internal state of the organism, contribute to ‘higher’ cognitive functions like learning and memory formation and modulate fundamental homeostatic needs such as sleep or feeding. Despite their significance and the extensive research in model organisms, the evolutionary origin of the monoaminergic system is uncertain. Here using a phylogenomic approach we study the evolution of the majority of genes involved in the production, modulation, and detection of monoamines. Our analyses suggest that most of the genes of the monoaminergic system originated in the common ancestor of bilaterians. These findings suggest that the monoaminergic synaptic pathway is a bilaterian innovation. We hypothesise that monoaminergic neuromodulation contributed to the diversification and complexification of behaviour and forms found in Bilateria.

## Introduction

By modulating neural activity, monoamines provide neural circuits with the functional plasticity required to adapt to the continuous remodelling of internal states and to the variability of environmental stimuli. Monoamines contribute to cognitive phenomena including learning, memory formation and emotions as well as to homeostatic processes such as feeding and sleep ^1,2^. However, despite the remarkable importance of the monoaminergic system, it is unclear whether it is a novelty that arose within Bilateria or if it evolved prior to them. Clarifying the *tempo and mode* of the origin of the monoaminergic system is fundamental to understanding the emergence of complex behaviours and their potential role in the diversification and complexification of animals.

All monoamines are synthesized from aromatic amino acids – primarily tyrosine and tryptophan – using multiple enzymes and co-factors^3^ (see Figure 1A). Typically, the first reaction adds a hydroxyl group to the amino acid. Members of the aromatic amino acid hydroxylase (AAAH) family, phenylalanine hydroxylases (PAHs), tyrosine hydroxylases (THs), and tryptophan hydroxylases (TPHs), mediate this. Then, follows the removal of a carboxyl group by members of the amino acid decarboxylase family (AADC) of enzymes, which includes dopa decarboxylases (DDCs), histidine decarboxylases (HDCs), and tyrosine decarboxylases (TDCs). Some monoamines, such as octopamine, epinephrine and norepinephrine, are modified further by the addition of a second hydroxyl group in a reaction catalysed by Cu(II) monooxygenases, such as dopamine beta-hydroxylases (DBHs), tyramine beta-hydroxylases (TBHs) and monooxygenase DBH-like (MOXDs). Other monoamines are methylated by phenylethanolamine-N-methyltransferases (PNMTs). Different combinations of these enzymes, but arranged in a similar order, produce all the monoamines described in animals. (Figure 1A). In addition to synthetic pathways, to function the monoaminergic system requires additional elements. Vesicular monoamine transporters (VMATs) concentrate the monoamines in vesicles before secretion into the synaptic cleft (Figure 1B). Several types of G-protein coupled receptors (GPCRs, e.g., dopaminergic receptors, serotonergic receptors, etc.) detect monoamines on either side of the synapse triggering signalling cascades, which induce modulation of electrophysiological responses. Finally, different proteins control the level of monoamines in the synaptic cleft by reuptake (such as transporters of the SLC6 family, like SerTs and DATs) or by degradation (by catabolic enzymes, like monoamine oxidases, MAOs).

**Figure 1:**
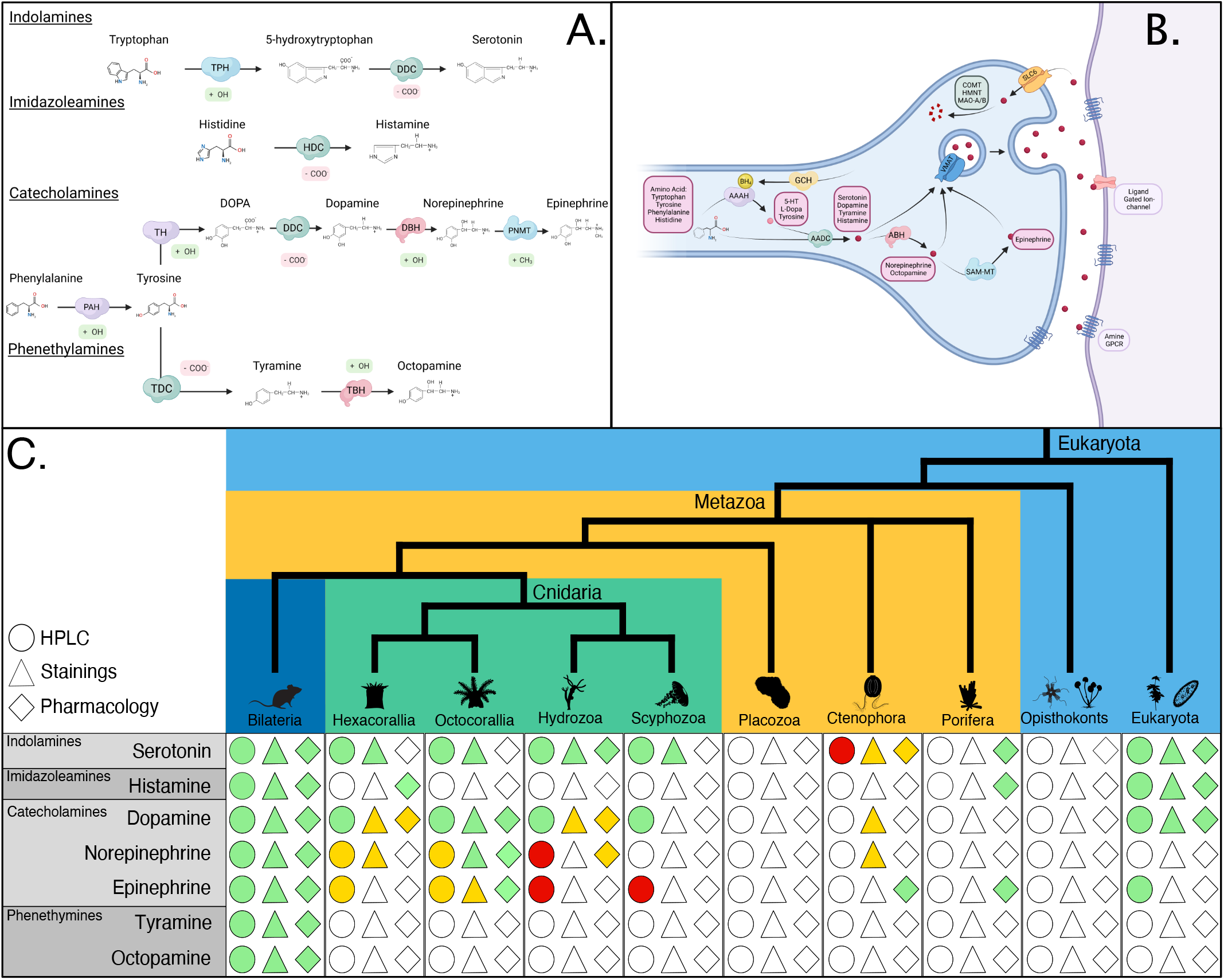
An overview of the monoamine system. **(A)** Synthesis pathways for key monoamines including the substrate, chemical modification, enzymes, and products. Arrows indicate reactions with the facilitating enzyme overlain. For each enzyme, the label shows the name while the colour and shape indicated the gene family it belongs to. Chemical modifications are shown next to the enzymes responsible. **(B)** Cartoon of a synapse with the different enzymes required for the production and detection of monoamines. **(C)** Current molecular evidence from the literature supporting the presence of monoamines outside Bilaterians in the literature. Green indicates positive results, orange displays uncertain or partial evidence (e.g., precursors, related compounds) and red shows negative results. Staining refers to any chemical or immuno-staining experiments; pharmacology refers to evidence-based drug perturbations, adding inhibitors or other chemical interference experiments; HPLC = High-Pressure Liquid Chromatography (see Table S1 for references and details). Figure A and B were made with Biorender and modified in Inkscape.

The distribution of monoamines and associated pathway components outside bilaterian animals remains unclear^4–6^. On the one hand, molecular and chemical evidence suggests a scattered distribution of monoamines outside bilaterians (summarized in Figure 1C and Table S1). On the other hand, previous studies ^4,7–14^ investigating the phylogenetic distribution of the monoaminergic pathway genes have not reached a definitive conclusion (summarized in Table 1). A main cause of uncertainty is the limited number of non-bilaterian genomes, such as those of sponges (Porifera), ctenophores, placozoans, and cnidarians, that have been investigated so far ^4,7–14^. Additionally, these studies have focussed on subsets of genes rather than on the monoaminergic system as a whole ^4,7–14^. Furthermore, understanding the evolution of single-gene families in ‘deep time’ is particularly challenging because of the weakness of the phylogenetic signal in single-gene alignments, especially if many sequences from distantly related organisms are included ^15,16^.

**Table 1.** Summary of previous work and main conclusions.

| Paper | Genes | Non-Bilaterians | Methods | Conclusions |
| --- | --- | --- | --- | --- |
| **Iyer et al, 2004** | AAAH, AADC, BH, PNMT, COMT, HNMT, MAO | 0 | Similarity; NJ Tree; ML Tree; Bayesian Tree | DDC and HDC diverge in Metazoa TH sister to TPH and PAH |
| **Caveney et al, 2006** | SLC6 | 0 | PCR; Similarity; NJ Tree | 3 transporter Clades Predate Bilateria |
| **Anctil, 2009** | AAAH, AADC, BH, PNMT, MAO, GPCRs, SLC6, VMAT | *Nematostella vectensis* (*Renilla sp.\**) | Similarity; Motif; NJ Tree; ML Tree | Homologs of most genes found in *Nematostella vectensis* |
| **Cao et al, 2010** | AAAH | 2 (*Nematostella vectensis*, *Trichoplax adhaerens*) | Similarity; NJ Tree; ML Tree | TPH and PAH sisters TH and TPH Bilaterian |
| **Siltberg-Liberles et al, 2008** | AAAH | 0 | Bayesian Tree | TPH and TH are sisters Duplications in Metazoa |
| **Kutchko and Siltberg-Liberles, 2013** | AADC, BH, MAO | 0 | Similarity; ML Tree | DDC Homolog to TDC and HDC, Metazoan unique TBH homolog of DBH and MOXD PNMT Vertebrate Specific |
| **Krishnan and Schiöth, 2015** | GPCRs, AAAH, AADC, BH, PNMT | 4 (*Nematostella vectensis*, *Trichoplax adhaerens*, *Mnemiopsis leidyi*, *Amphimedon queenslandica*) | Similarity | Whole Pathway Present in Cnidaria DDC, DBH and GPCRs in other non-bilaterians |
| **Francis et al, 2017** | AADC, BH, MAO | 2 Placozoans; 36 Cnidarians; 20 Sponges; 13 Ctenophores | Similarity; ML Tree | DBH Homologs in non-bilaterians DDC Ortholog in Hydrozoa DDC Homologs in non-bilaterians |
| **Moroz et al, 2021** | AAAH, AADC, SLC6, BH, PNMT, VMAT, 5HT3R | *Trichoplax adhaerens*; 6 Cnidarians; *Mnemiopsis leidyi*; *Pleurobrachia bachei*; *Amphimedon queenslandica* | Uncertain | Only DDC and DBH have Orthologs in non-Bilateria |

Here, we present a systematic investigation of the canonical genes involved in the monoaminergic system. We have studied the pattern of duplication of 18 genes that encode proteins involved in the synthesis, turnover, or reception of monoamines. We took advantage of modern phylogenomic data using a wide sample of animals, especially non-bilaterians, and their close relatives. Our approach deployed methods to minimize the effect of unstable sequences and have reconstructed the pattern of gene duplication using recently developed maximum likelihood reconciliation techniques. Our results strongly indicate that many orthologs of the genes involved in the monoaminergic pathways originated in the Bilateria stem-group. This supports the conclusion that the monoaminergic system is a bilaterian innovation.

## Results

### The key enzymatic machinery for monoamine synthesis is a bilaterian novelty

Common problems with previous studies were the limited taxonomical sampling and/or the use of limited phylogenetic methods (see Table 1 and ^4,7–14^). To overcome these limitations, we sampled 47 animal species, 21 non-bilaterians and 18 opisthokonts (Table S2, and Methods for further details). To minimize bias associated with poor quality genomes we selected species with a high BUSCO completeness score^17^ (see Table S2) while maintaining large taxonomical diversity. After homology and orthogroup identification (see Material and Methods and Table S3 and S4), we computed gene trees using maximum likelihood inference (ML). In addition to ultrafast bootstrap^18^ (UFB), nodal support was estimated using the transferable bootstrap expectation score^19^ (TBE), a method that has been designed to account for and identify short and problematic sequences that have limited or conflicting phylogenetic signals^19^. Importantly, the presence of rogue/unstable sequences can affect the phylogenetic relationships and the pattern of duplication. To mitigate this effect, we used the Leaf Stability index^20^ as implemented in RogueNaRok^21^ and the t-index ^19^ from the TBE analysis) to identify unstable taxa. Finally, we used GeneRax, a recently developed ML method ^22^ to estimate the pattern of duplication and losses.

The first orthogroup we investigated included all the sequences encoding for the aromatic amino acid hydroxylase family (AAAH), enzymes that add a hydroxyl group to the aromatic ring of amino acid. This includes three key enzyme types: phenylalanine hydroxylases (PAHs) that synthesise tyrosine from phenylalanine; tyrosine hydroxylases (THs) which mediate the initial step in the synthesis of dopamine, epinephrine, and norepinephrine; and tryptophan hydroxylases (TPHs), which start the synthesis of serotonin (see Figure 1A). The phylogenetic tree (Figures 2A, S1A and B) recovered the monophyly of TPHs (TBE=0.99, UFB=100), THs (TBE=0.99 and UFB=98) and TPHs plus THs (TBE=0.97 and UFB=75). Unlike PAHs, TPHs and THs were unique to Bilateria, apart from a single sequence from the sponge *Amphimedon queenslandica*. However, both the results of the LSI and the t-index (see methods for more details and Table S5) suggested that the phylogenetic position of this sequence was unstable and was likely a phylogenetic artefact. The absence of TPHs or THs in other sponges we analysed further confirmed this conclusion. We removed the *Amphimedon queenslandica* sequence from the dataset and performed the gene tree to species tree reconciliation. This result corroborated the phylogenetic trees indicating that THs and TPHs originated in the bilaterian stem group (Figures 1B, S1C and D) and that they represent a bilaterian innovation.

**Figure 2:**
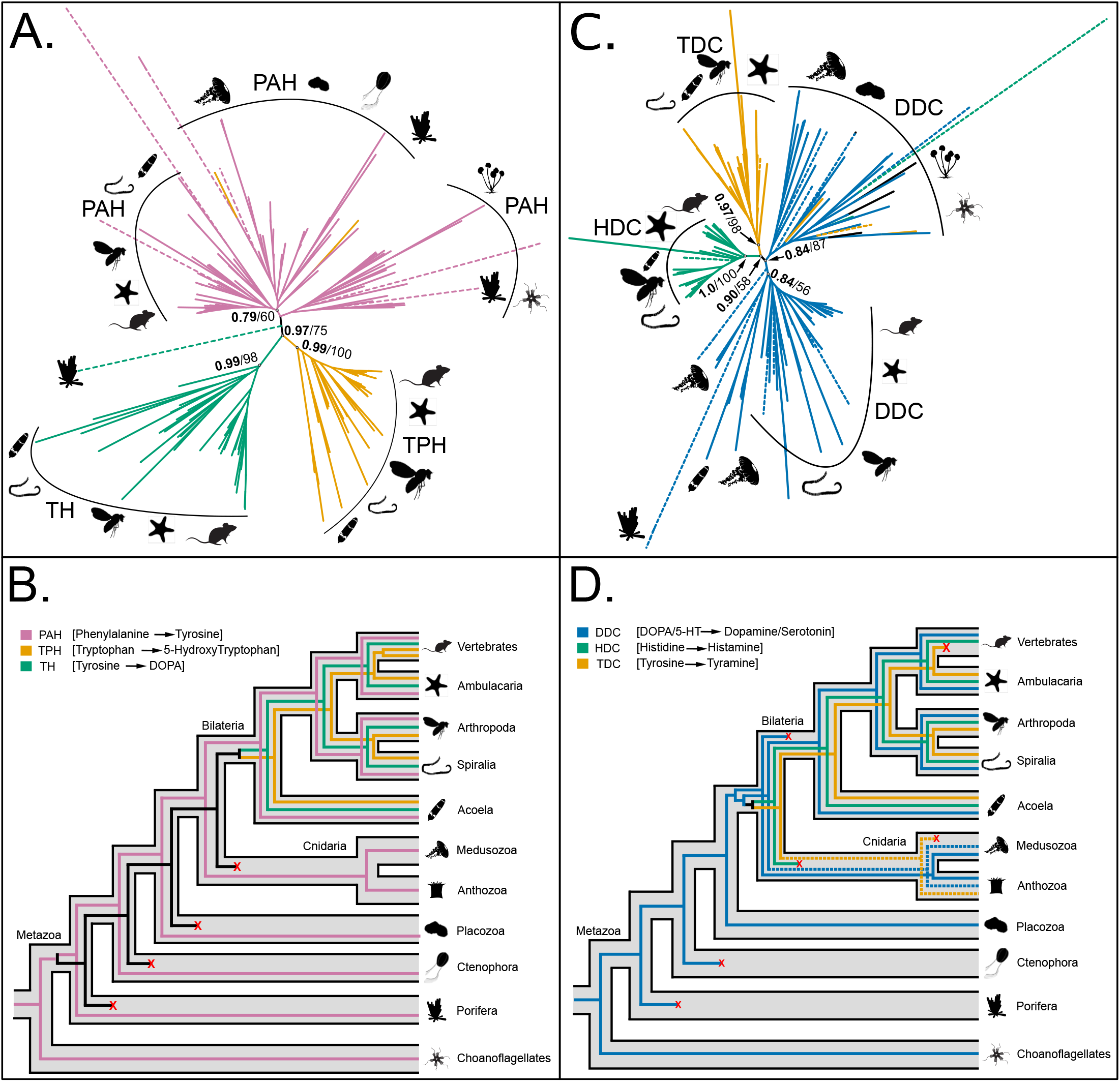
Phylogeny and reconciliation for aromatic amino acid hydroxylases (AAAHs) and amino acid decarboxylases (AADCs). **(A)** Transfer bootstrap expectation tree and **(B)** simplified illustration of reconciliation calculated using Generax for AAAH sequences. **(C)** Transfer bootstrap expectation tree and **(D)** simplified illustration of reconciliation calculated using Generax for AADC sequences. The nodal supports shown are TBE scores (in bold), and ultrafast bootstrap proportion supports (in italic) for key nodes. Dashed lines indicate sequences identified as unstable in the t-index and LSI analysis (see table S5 for details).

Amino acid hydroxylases require the cofactor tetrahydrobiopterin to function ^3^, which is synthesised by GTP cyclohydrolases (GCHs) among other enzymes^3^. Our phylogenetic tree (Figure S2A, S3A and B) for the GCHs suggested that differently from the aromatic amino acid hydroxylases, the GCHs are found across all animal species and opisthokonts (Figure S2A, SA and B). Phylogenies and reconciliations show that the gene tree closely resembled the species tree suggesting that there are no major duplication or losses events in this gene family (Figure S2B, S3C and D). GCHs are not specific to monoaminergic pathways^3^, which may explain their wider distribution.

Next, we investigated the evolution of aromatic amino acid decarboxylases (AADC) enzymes, which remove a carboxyl group as part of monoamine synthesis^3^ (Figure 1A). Enzymes of the AADC family include dopa decarboxylases (DDCs), histidine decarboxylases (HDCs) and tyrosine decarboxylases (TDCs). The phylogenetic tree (Figure 2C, S4A and B) recovered the monophyly of HDCs and TDCs (TBE=0.9 and UFB=58). Both groups included only bilaterian sequences, except putative TDCs from the cnidarian *Paramuricea biscaya*. In contrast, we identified two clades of DDC encoding genes. The first included mainly bilaterian sequences (TBE=0.86, UFB=69) while the second included the majority of non-bilaterian and non-metazoan sequences (TBE=0.84, UFB=87). In the first DDC clade, the sequences from the sponge *Sycon ciliatum* and the cnidarian *Hydra magnipapillata and Clytia hemisphaerica* (TBE=0.57) were positioned as a sister group to the bilaterian DDCs (TBE=0.85). Analyses by LSI and the t-index (see Table S5) identified the sequences from *S. ciliatum* as unstable (supported by the alternate positioning in the UFB tree Figure S4B) but not the sequences from *H. magnipapillata and C. hemisphaerica* (see Table S5). Likely the latter represent putative DDC orthologs, which matches a previous observation^7^. The reconciliation analysis (Figure 2D, S4C and D) performed after excluding the unstable sequences (see above) suggested that HDCs and TDCs originated in the stem group of Cnidaria and Bilateria. However, this scenario relies only on sequences from a single species (the coral *P. biscaya*). For DDCs, the reconciliation suggested a similar pattern, origin in the stem group of Cnidaria and Bilateria. This requires the loss of DDC orthologs in all cnidarians except hydrozoans (Figure 2D, S4C and D). In summary, our results suggest that DDCs, HDCs and TDCs are almost exclusive to Bilateria, with the reconciliation placing their origin in the stem lineage of Cnidaria and Bilateria. However, this ‘earlier’ scenario (compared to origin in the stem lineage of Bilateria) depends on a few and sparsely distributed cnidarian sequences.

Dopamine beta hydroxylases (DBHs), tyramine beta hydroxylases (TBHs), and monooxygenase dopamine beta hydroxylase-like (MOXDs) are involved in the synthesis of norepinephrine and octopamine^3^ (Figure 1A). In vertebrates, DBHs are used to synthesize norepinephrine^3^ while in arthropods TBHs are involved in the production of octopamine^3^. MOXDs are not functionally characterized. The phylogenetic tree (Figure 3A, S5A and B) supported the monophyly of MOXDs (TBE=0.94, UFB=87) and of DBHs and TBHs (TBE=1.0, UFB=100), which were almost exclusively limited to bilaterians except for a single sequence from *P. biscaya* in both clades. While the TBE tree placed the non-bilaterian sequences as the sister group of the DBHs/TBHs clade (TBE=0.64), the UFB tree assigned them as the sister group of the bilaterian MOXDs (UFB=48) (Figure S5A and B, respectively). To consider these differences, we performed the reconciliation analysis on both trees (Figure S5C, D, E and F). Both analyses supported the split between MOXDs and DBHs/TBHs occurring in the ancestor of Cnidaria and Bilateria (Figure 3B and Figures S8C, D, E and F). However, they differed in the classification of cnidarians sequences as orthologs of DBHs/TBHs (as in the TBE tree, Figure S5A) or of MOXD (as in the UFB tree, Figure S5B). Furthermore, contra to previous observations ^23^, both trees (Figure 3A and B) indicated that the fruit fly TBH is orthologous (1:1) to human DBH, suggesting that this name distinction is purely semantic (see Figure 3A and B).

**Figure 3:**
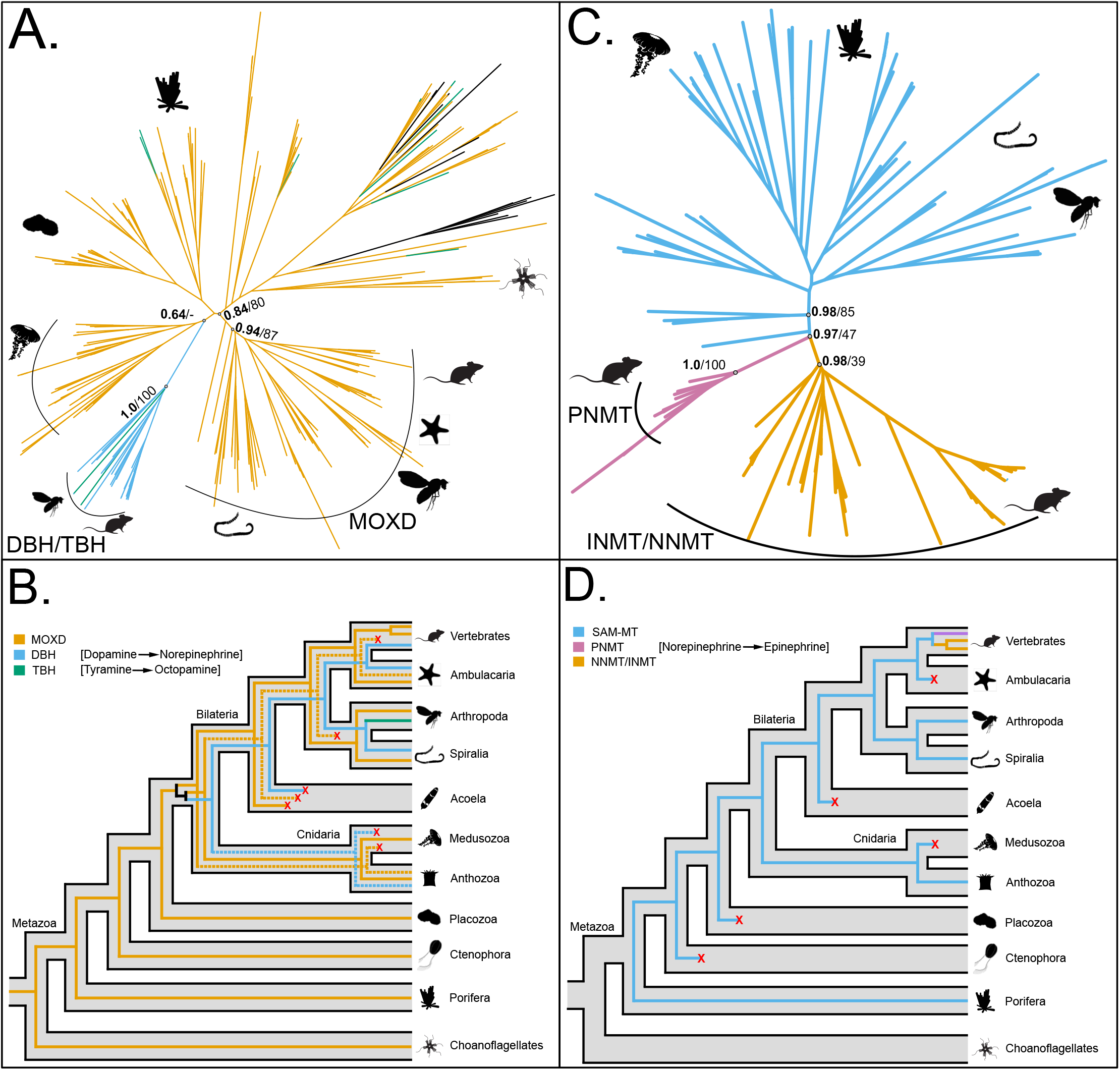
Phylogeny and reconciliation for beta-hydroxylases (BHs) and phenylethanolamine-N-methyltransferases (PNMTs). **(A)** Transfer bootstrap expectation tree and **(B)** simplified illustration of reconciliation calculated using Generax for BH sequences. **(C)** Transfer bootstrap expectation tree and **(D)** simplified illustration of reconciliation calculated using Generax for PNMT sequences. The nodal supports shown are TBE scores (in bold), and ultrafast bootstrap proportion supports (in italic) for key nodes. Dashed lines indicate sequences identified as unstable in the t-index and LSI analysis (see table S5 for details).

Phenylethanolamine-N-methyltransferases (PNMTs) catalyse the last step in the production of epinephrine from norepinephrine and belong to the SAM-binding methyltransferase superfamily (SAM-MT) ^24^. Our analysis indicated that the enzymes of the SAM-MT family are widely distributed across Metazoa, but, PNMTs are uniquely present in vertebrates (TBE=1, UFB=100 Figure 3C, S6A and B). Additionally, we showed that these sequences form a monophyletic group with nicotinamide-N-methyltransferases (NNMTs) and indolethylamine-N-methyltransferases (INMTs), both also unique to vertebrates (Figure 3C). The reconciliation (Figure 3D, S6A and B) suggested that PNMTs diverged from the INMTs/NNMTs at the time of origin of the jawed vertebrates (gnathostomes).

### Transporter gene families predate the origin of neurons but monoamine-specific orthologs are unique to Bilateria

Transporters concentrate monoamines inside vesicles for secretion into the synaptic cleft^3^. In Bilateria vesicular monoamine transporters (VMATs) perform this role. These are part of the solute ligand carrier family 18 (SLC18), along with vesicular acetylcholine transporters (VACHTs)^25^. The phylogeny of the SLC18 orthogroup (Figure 4A and B) supported the monophyly of bilaterian VMATs (TBE=0.86, UFB=44) and VACHTs (TBE=1.0, UFB=100) (Figure 4A, S7A and B, and Table S4). Three cnidarian sequences were placed within the VMATs clade; however, the LSI and the t-index identified them as unstable (see Table S4). Furthermore, our phylogenetic results supported the monophyly of VMATs and VACHTs (TBE=0.99, UFB=100, see Figures S7A and B). Interestingly, the analysis placed some sequences from sponges, choanoflagellates and other holozoans as outgroups of the bilaterian VMAT/VACHT clade (Figure 4A and S7A and B). The reconciliation performed after removing the unstable cnidarian sequences confirmed these findings, namely that VMATs and VACHTs descend from a duplication in the bilaterian stem group (Figure 4B, S7C and D) and that all non-bilaterians except sponges lack VMATs/VACHTs orthologs.

**Figure 4:**
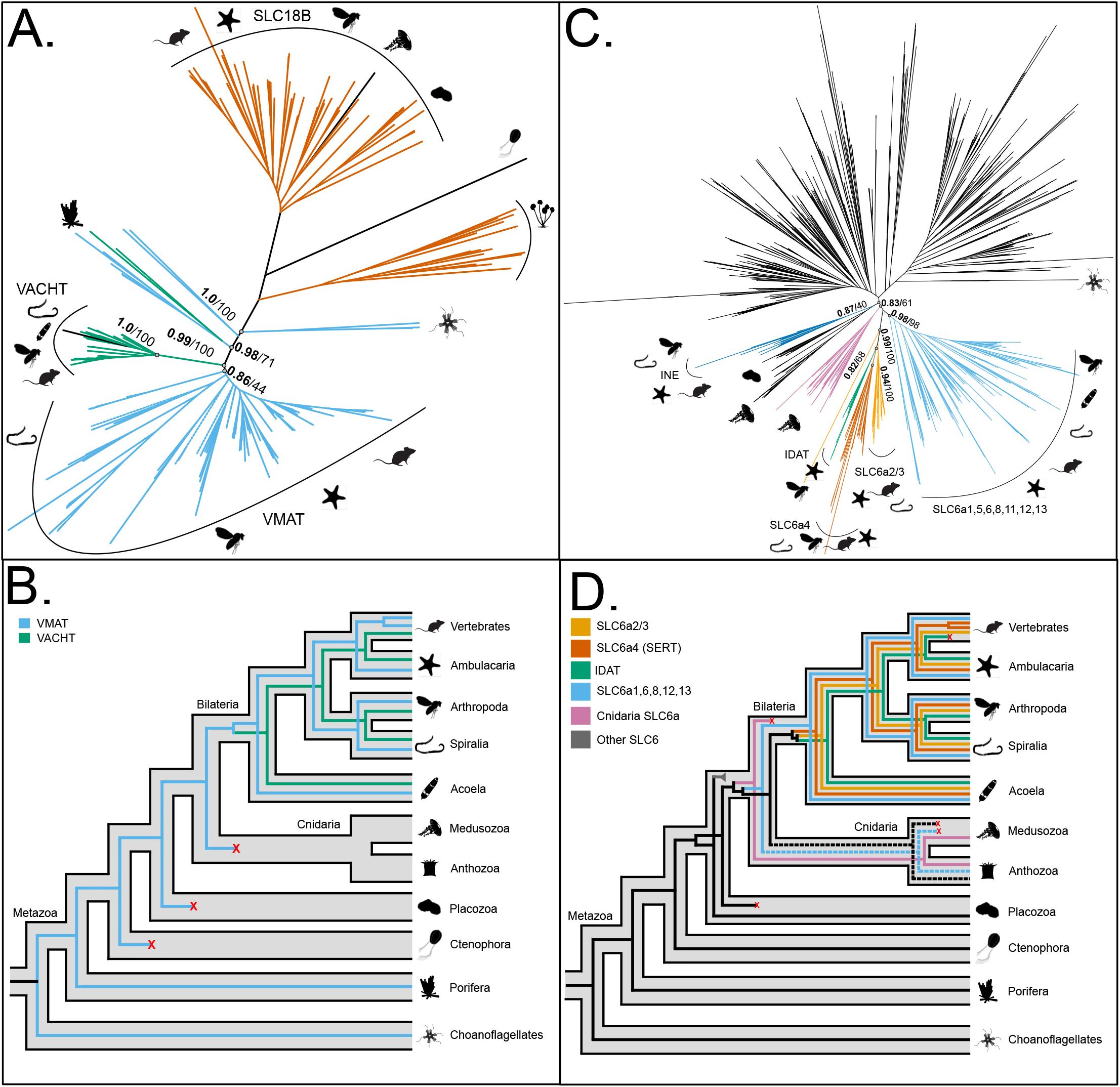
Phylogeny and reconciliation for vesicular monoamine transporters (VMATs) and Solute ligand carrier 6 (SLC6) family members. **(A)** Transfer bootstrap expectation tree and **(B)** simplified illustration of reconciliation calculated using Generax for VMAT sequences. **(C)** Transfer bootstrap expectation tree and **(D)** simplified illustration of reconciliation calculated using Generax for SLC6 sequences. The nodal supports shown are TBE scores (in bold), and ultrafast bootstrap proportion supports (in italic) for key nodes. Dashed lines indicate sequences identified as unstable in the t-index and LSI analysis (see table S5 for details).

SLC6 is a large family of transporters including those for serotonin (SerTs/SLC6a4), dopamine in invertebrates (iDATs) and dopamine/epinephrine/norepinephrine in vertebrates (SLC6a2/3s). These proteins contribute to the regulation of the concentration of monoamines in the synaptic cleft^3,11^. All monoamine transporters formed a clade almost exclusive to bilaterians (TBE=0.99, UFB=100), apart from a single sequence from the cnidarian *P. biscaya* (Figure 4C, S8A and B). Our analyses suppored the monophyly of SLC6a4 (TBE=0.94, UFB=100), iDATs (TBE=0.98, UFB=100), and SLC6a2/3s (TBE=0.96, UFB=47). Furthermore, we identified a sister group relationship between the transporters for monoamines and a cluster including transporters for GABA, taurine and creatine (TBE=0.79, UFB=50), which were mainly identified in bilaterians (i.e., SLC6a1s, SLC6a6s and SLC6a8s in Figure 4C). Both these transporter clades were recovered as the sister group to a large cnidarian-specific clade (TBE=0.83, UFB=61). The reconciliation analyses suggested that the monoamine transporters originated in the stem of Bilateria (Figure 4D, S8C and D). Based on a single sequence from *P. biscaya* we identified an ortholog to all monoamine transporters in corals. However, evidence from more species is needed to corroborate this finding.

### Evolutionary history of monoamine catabolic enzymes

The correct functioning of neural circuits requires tight control of the level of monoamines. Several catabolic enzymes are used in animals for breaking down/inactivating monoamines, which provide an key mechanism of regulation^26^. In bilaterians, this is mediated by Catechol-o-methyltransferases (COMTs), Histamine-N-methyltransferases (HNMTs) and monoamine oxidases (MAOs) (see Figure 1B).

COMTs add a methyl group and consequently inactivate dopamine, epinephrine, and norepinephrine^27^. The phylogenetic trees and reconciliation analyses (Figure S9 and S10) identified COMTs as primarily present in deuterostomes. The phylogenies placed echinoderm COMTs as a sister group to the vertebrate transmembrane-O-methyltransferase clade (TBE=0.65 and UFB=60), which also may be able to inactivate dopamine, epinephrine, and norepinephrine^28^. Additionally, this orthogroup included sequences from two corals (*P. biscaya* and *Heliopora coerulea*), two sponges *(S. ciliatum* and *Oscarella pearsei*), a rotifer (*Adineta vaga*) and several non-metazoans (Table S3). The reconciliation inferred many secondary losses across Metazoa (Figure S9 and S10) Thus, COMTs seem to contrast with other monoaminergic components that usually appear conserved across Bilateria (see Table S3).

Likewise, the enzymes of the Histamine-N-methyltransferase (HNMTs) family inactivate histamine by adding a methyl group^29^. We identified HNMTs in Deuterostomia, Acoela, and Anthozoa (Cnidaria), with sequences present in almost every species from these groups (Table S3). Phylogenetic trees and reconciliation analyses (Figure S11 and S12) suggested that HNMTs originated in the ancestral group of Cnidaria/Bilateria followed by many secondary losses (Figure S11B and S12C and D).

Monoamine oxidases (MAOs) inactivate all monoamines as well as other amino acid derivatives such as tryptamine, benzylamine and kynuramine^30^. Both the TBE and the UFB trees (Figure S13A, S14A and B) supported the monophyly of bilaterian MAOs including both vertebrate paralogs; TBE=0.92, UFB=43). We identified a monophyletic clade of cnidarian sequences (TBE=0.98, UFB=86) as sister to the bilaterian MAOs. Several choanoflagellate and other eukaryote sequences also made up part of the larger MAO clade the metazoan sequences were in (TBE=0.99, UFB=86) (Figure S13A, S14A and B). Furthermore, we recognise a MAO-like group that including animal and non-animal sequences, as the sister group to the large MAO-A/B clade (TBE=0.93, UFB=66). Interestingly, Medusozoa, Porifera and *D. melanogaster* retain MAO-like sequences only (Figure S13A, S14A and B). We did not identify any MAO in ctenophores or placozoans, while at least one homolog from each clade (MAO-A/B and MAO-like) is present in all other animals groups (Figure S13A, S14A and B). The reconciliation analysis suggested that the two major MAO clades have an ancient origin, which significantly predates the origin of animals and neurons (Figure S13B, S14C and D).

### Most monoaminergic receptors are specific to bilaterians

Once released in the synaptic cleft, monoamines are detected by specific transmembrane proteins expressed on both sides of the synapse. These receptors are the effectors that trigger signalling cascades inducing a physiological response. Most monoamine receptors are G-protein-coupled receptors (GPCRs) of class A^5,14^. However, serotonin can be detected also by ligand-gated ion channel receptors from the cys-loop repeated gene family^3^ (5HT3Rs).

Our analyses revealed that all putative 5HT3R sequences form an orthogroup with the ligand-gated zinc-activated ion channels. This group comprises vertebrate sequences and some urochordate and acoel sequences (see Table S3). The phylogenetic tree indicated that vertebrate 5HT3Rs are monophyletic (TBE= 0.96, UFB = 86; Figure S15A and S16A and B). The reconciliation analysis suggested that several losses occurred in bilaterians. On the contrary 5HT3Rs underwent an expansion in vertebrate lineages (Figure S15B and S16C and D).

Then we turned our attention to the GPCRs. Our analysis split the monoaminergic GPCRs into two orthogroups (see Table S3). The first, hereafter OG30, contained histamine receptors 1, 3 and 4 (HRH1, HRH3, HRH4) and acetylcholine muscarinic receptors (ACMs). It included bilaterian and non-bilaterian sequences. The second, hereafter OG1, comprised the remaining known monoaminergic GPCRs, including the receptors for serotonin, dopamine, epinephrine/norepinephrine, octopamine/tyramine, trace amines and histamine receptor 2 (HRH2). OG1 also included sequences from bilaterians and non-bilaterians (Table S3). However, given the taxonomic composition and the lack of outgroups, it was not possible to understand the pattern of duplication and the following orthology relationships.

To overcome this limitation, we expanded our analysis to include all GPCR orthogroups. To reduce the computational burden, we selected only GPCRs with seven transmembrane domains (see methods) and non-redundant sequences (see methods). Using this approach, we focussed on 2,837 GPCRs and using CLANs^31^ (a similarity-based method) to identified clusters of related sequences. Applying a p-value=1e^-60^ we observed that the majority of bilaterian sequences from OG1 and OG30 clustered together (Figure S19A). To identify potential outgroups, we relaxed the parameters and used a p-value=1e^-40^ (Figure S19C). At this threshold we observed that OG1, OG30, adenosine receptors, and melatonin receptors fromed a cluster with many interconnected sequences and that this group has sporadic connections with opsins and other GPCRs. To re-focus monoaminergic GPCRs, we increased the p-value to 1e^-42^ (Figure S19B) and identified 1,277 sequences connected to OG1 and OG30.

While CLANs is a powerful tool to investigate the relationships between a large numbers of sequences, this approach cannot clarify the specific evolutionary history and patterns of gene duplication. Thus, we performed phylogenetic analyses using opsin sequences as the outgroup. Both the UFB and the TBE trees (Figure 5A, S20 A and B) provided support for the monophyly of OG30 (TBE=0.97 and UFB=74) and of OG1+OG30 (TBE=0.96 and UFB=25). These clades are composed mainly of bilaterian sequences except for a few from cnidarians that are the sister group to ACM receptors within OG30. Additionally, both trees supported the existence of a large cnidarian-specific monophyletic group (TBE=0.65, UFB=66). We also identified a clade of adenosine receptors (TBE=0.73, UFB=63), which included sequences from bilaterians, cnidarians and placozoans. The adenosine receptors clade was placed as the sister the group comprising OG11, OG30 and the cnidarian-specific clade. The main disagreement between the UFB and the TBE trees was the position of OG1. The UFB tree recovered OG1 as monophyletic, albeit with low support (UFB=37). The TBE tree split OG1 in two clades with OG30 nested within it (TBE=0.70) (Figure 5A, S20 A and B). Another difference was the phylogenetic position of two placozoan sequences that were placed within the OG1 by the UFB tree but were considered a sister group to OG1 and OG30 by the TBE tree. However, note that these sequences were identified as unstable by the LSI analysis (Table S4).

**Figure 4:**
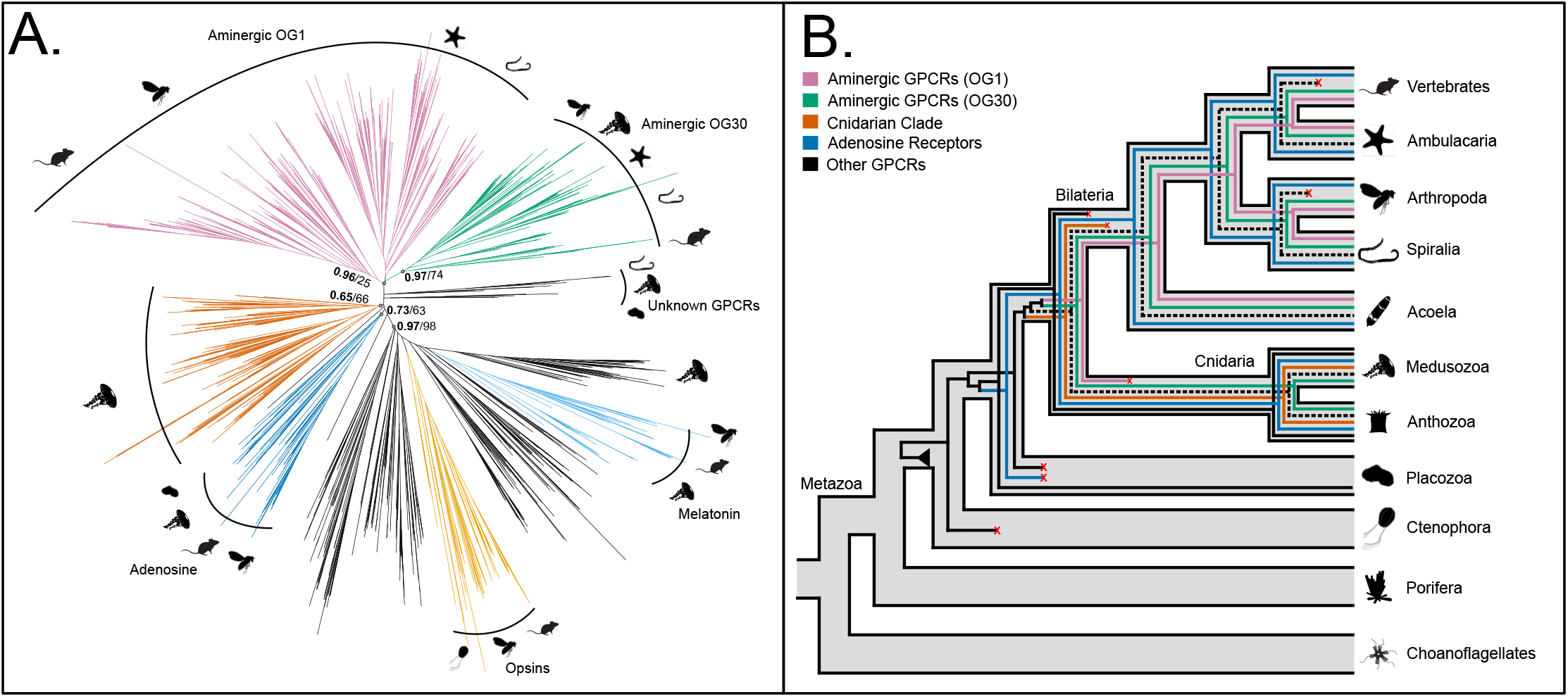
Phylogeny and reconciliation of the monoaminergic GPCRs. **(A)** Transfer bootstrap expectation tree and **(B)** simplified illustration of reconciliation calculated using Generax for monoaminergic GPCRs. The nodal supports shown are TBE scores (in bold), and ultrafast bootstrap proportion supports (in italic) for key nodes. Dashed lines indicate sequences identified as unstable in the t-index and LSI analysis (see table S5 for details).

The reconciliation performed using the TBE and UFB trees supported that OG1, OG30 and the cnidarian specific clade originated from a duplication that gave rise to the adenosine receptors and to a “proto-monoamine” receptor in the ancestor of placozoans, cnidarians and bilaterian. The “proto-monoamine” receptor subsequently gave rise, in the cnidarian/bilaterian stem group, to the canonical monoamine receptors (OG30 and OG1) and to the cnidarian-specific clade (Figure 5B, S20C, D, E and F). Both trees supported OG1 and OG30 originating from a single gene in the cnidarian/bilaterian stem group. Subsequently, OG1 underwent a significant expansion in the bilaterian stem group giving rise to the modern bilaterian monoaminergic receptors. However, the two trees disagree on the precise number of copies of OG1 in the common ancestor of bilaterians with UFB indicating a single copy (Figure S20C and D) and TBE suggesting multiple copies (Figure S20E and F).

In summary, our results suggest that the monoaminergic GPCRs originated from a gene duplication event that also gave rise to the adenosine receptors in the eumetazoan stem-group. Modern monoaminergic receptors can be divided into two subclades. The first is bilaterian specific and contains receptors for serotonin, dopamine, epinephrine/norepinephrine, octopamine/tyramine, trace amines and HRH2. The second includes HRH1, HRH3, HRH4 and ACM receptors comprising both bilaterian and cnidarian sequences. Our reconciliation analyses suggest that the two subclades diverged in the Bilateria/Cnidaria stem group.

## Discussion

In this study, we have provided strong phylogenetic evidence that the monoaminergic system, with the necessary complexity required to modulate neuronal circuits, is a bilaterian innovation. Our results (summarized in Figure 6A) allow for a substantial clarification of the *tempo and mode* of its evolution.

**Figure 6:**
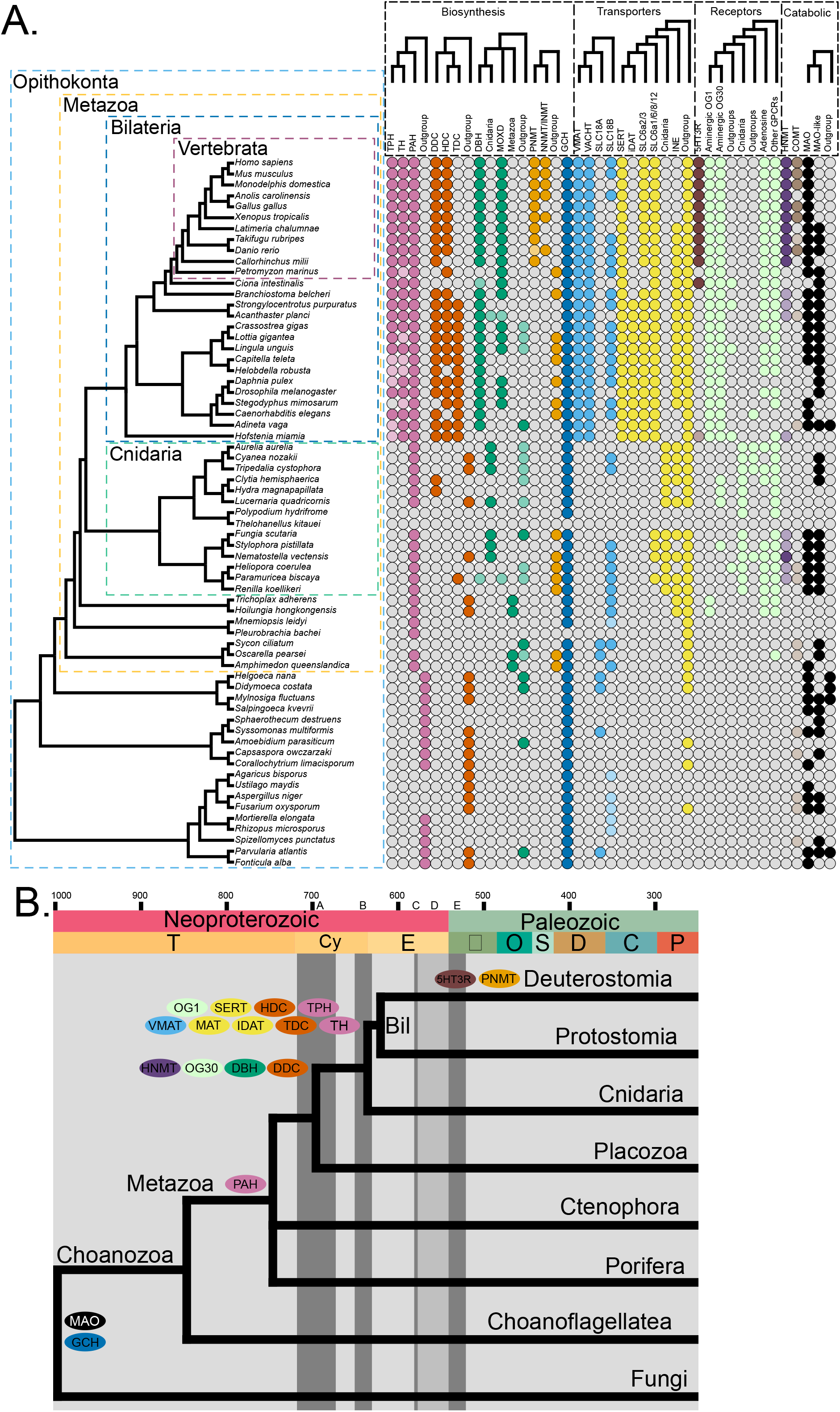
Synopsis of the evolution of the monoaminergic system. **(A)** Presence/absence of monoamine pathway genes inferred using the reconciliation analysis. Colours correspond to orthogroups, with darker shades indicating matches to specific InterProScan profile (Table S4). **(B)**. A simplified time-calibrated species tree illustrating the origin of the key monoaminergic genes and the presence of neurons. A = Sturtian glaciation; B = Marinoan glaciation; C = Gaskiers glaciation; D = Occurrence of Ediacaran Biota/early animal fossils; and E = the Cambrian explosion. Species tree dated following Dos reis *et al*, 2015^62^, Geological column is shown in accordance with the ICS International Chronostratigraphic Chart (updated 2017) ^63^.

Compared to previous work^4,7–14^ we have included a larger taxonomic sample, which has allowed us to clarify further the evolution of the monoaminergic enzymatic machinery and that of the receptors. We have used novel statistical methods to pinpoint unstable taxa and to reconcile the gene trees with the species tree, which has improved the robustness of our phylogenetic inferences. We have confirmed previous observations, for instance that THs and TPHs are present only in Bilateria^4,8,12^. However, our main contribution is that we were able to show that a large majority of monoaminergic genes evolved in the Bilateria stem group and that others, such as 5HT3Rs and PNMTs, originated in vertebrates. The distribution of orthologs summarized in Figure 6A, suggests a scenario where the monoaminergic system was assembled by co-opting old enzymes with a broad function (such as GCH and MAO, which predate Metazoa and neurons) alongside newly evolved bilaterian-specific genes (such as TPH and TH) into a new functional unit.

In spite of our evidence the conclusion that the monoaminergic system is a bilaterian innovation may seem in conflict with existing experimental observations reporting monoamines in non-bilaterian metazoans^6^ (Figure 1C). How then could non-bilaterians produce monoamines? One possible solution is the functional flexibility of the synthetic enzymes. For example, previous studies have indicated that PAHs could produce serotonin in *Drosophila melanogaster* and mice although less efficiently than TPHs ^32–34^. Additionally, mutational analyses of mammalian AAAHs have shown that a few simple mutations can change the substrate specificity of PAHs from phenylalanine to tryptophan^35,36^. Therefore, some non-bilaterian animals might use PAH to produce serotonin. Additionally, some non-bilaterians have a single homolog of DDC, HDC and TDC (see Figure 6A). In some species the encoded enzyme may be able to carry out some or all the enzymatic functions of their bilaterian counterparts. Another possibility is the existence of alternative pathways for monoamine synthesis. For instance, it has been proposed that the hydroxylation of tyrosine into L-DOPA could be mediated by a tyrosinase instead of TH in *Hydra* and in the sea anemone *Metridium senile*, based on DOPA by-products detected by HPLC ^37,38^. TH-independent dopamine synthesis has also been invoked in *Caenorhabditis elegans*, to explain that TH-deficient mutants still have ∼40% of normal dopamine levels ^39^. This alternative synthesis has been proposed to participate in the synthesis of peripheral dopamine in young mice ^40,41^.

Our phylogenetic analysis suggests that the receptors for serotonin, dopamine, epinephrine/norepinephrine, octopamine/tyramine, trace amines and the HRH2s are uniquely present in Bilateria. Thus, they originated from a single gene duplication event in the cnidarian/bilaterian stem group. While the orthologs of histamine receptors 1, 3 and 4 are limited to Bilateria, they appear to be secondarily absent in Cnidaria, with just the ACM receptors conserved across both Cnidaria and Bilateria. We also identified an independent expansion of GPCRs that are unique to Cnidaria but closely related to the canonical monoaminergic receptors. The functional role of these cnidarian receptors is yet to be determined but, we speculate that they could drive the response to monoamines observed in cnidarians^6^.

Overall, our data indicate that the key genes for the monoaminergic system emerged around the Cryogenian/Ediacaran boundary about 650-600 Mya (Figure 6B), pre-dating the Cambrian explosion (540 Mya). In the pre-Cambrian, the earth system underwent radical changes in the geochemical condition of the oceans, providing new opportunities for animals to explore new ecological niches^42^. Paleontological evidence from this time period suggests that animals, previously limited to filter-feeding and low-motility lifestyles (as exhibited by most non-bilaterians), started evolving more energetically intensive and complex modes of life. This required the evolution of complex behaviours involving environmental sensing, target-directed locomotion and predation^42^. We speculate that monoamines played a crucial role in the development of complex behaviour in Bilaterian. They became molecular tools that were able to modulate neuronal circuits in a localised fashion, increasing complexity and functional plasticity of the nervous system. Importantly monoamine’s conserved use energy regulation and activity states may be vital, allowing early bilaterians to modulate their behaviour and adapt to the more complex ecosystems they found themselves in. Thus, the origin of the monoaminergic system in the pre-Cambrian provided and unprecedented opportunity to the nervous system for further complexity to evolve at a time when the earth system was experiencing exceptional changed (e.g., increase of O2 levels in the oceans^42^). We suggest that a new functional organisation of the nervous system coupled to ‘ecological need’ sparked the arm-race that led to the complexification and diversification of body plans.

In summary, our work provides strong phylogenomic evidence supporting a stem-bilaterian origin of the monoaminergic system (Figure 6A and B). Further studies will inform on the molecular details, such as the function of related enzymes and GPCRs in non-bilaterian animals. In future work, the further availability of high quality genomes (e.g., chromosomal level assembly) for many more organisms will afford a more precise characterization of orthology relationships.

## Methods

### Species Selection

To assess the completeness of the proteomes we used BUSCO^17^ genes with the eukaryota_odb10 database composed of 255 single-copy genes. Seven cnidarians (see table S1) proteomes were assembled starting from transcriptome using Trinity ^43^ and Transdecoder ^44^. Initially, we selected 101 species (including 47 animals and 54 eukaryotes) considering both taxonomical diversity and BUSCO completeness score (see table S1). However, after preliminary phylogenetic analysis, we focused on 65 opisthokonts species (see below for details).

### Homologs and orthogroup identification

The KEGG pathways ^45^ for serotonergic and dopaminergic synapses as well as published literature were used to identify monoamine pathway genes (See Table S4). To identify homologous sequences for each gene of interest we used BLASTP ^46^ with known sequences from SwissProt ^47^, EggNOG ^48^ or KEGG ^45^ as seeds (detailed in Table S2). All BLAST hits with an e-value < 1e^-25^ were extracted and annotated using BLASTP against the SwissProt database.

Starting from the putative homologs to identify the orthogroups we used Broccoli using default parameters and the maximum likelihood tree reconstruction method ^49^. Each orthogroups were then annotated based on sequences from *Homo sapiens, Mus musculus* and *Drosophila melanogaster* (Table S3). Furthermore, orthogroups of interest were analysed using InterProScan ^50^ and manually inspected for matches to profiles associated with known monoamine genes (details in Table S4).

Initial inspection of the orthogroup distributions and preliminary maximum likelihood trees (see below for details) showed that most sequences of interest are present in metazoans and their close relatives (Table S3). Given this taxonomical distribution for further analysis, we focused on opisthokonts species.

### GPCR analysis

We assembled a dataset of monoamine receptors by combing the different Broccoli orthogroups with GPCR annotations (see main text for justification). Phobius ^51^ was used to estimate the number of transmembrane domains (TMDs) and sequences with 7TMDs were kept and CD-hit was used to eliminate homologs with >80% similarity. We used CLANs ^31^ to cluster sequences based on their similarity across different stringency threshold p-values (from 1e^-100^ to 1e^-30^). Similarly, to the orthogroups the clusters were annotated using sequences from *Homo sapiens, Mus musculus* and *Drosophila melanogaster*. All sequences connecting to the cluster which contained the known aminergic GPCRs at p-value=1e^-42^ were extracted, and opsin sequences were used as outgroup.

### Phylogenetic Analyses

#### Gene trees

Each gene family of interest was aligned using MAFFT ^52^ using (--auto option and 1,000 max iterations) and we removed sites with >70% gaps using trimAl ^53^.IQ-TREE2 ^54^ was used to reconstruct the gene trees under the best fitting model accordingly with BIC (Table S2). Node support was calculated using 1,000 Ultrafast Bootstrap ^55^ (UFB) repeats. We also estimated nodal support using transferable bootstrap expectation (TBE) ^19^ scores from 100 non-parametric bootstrap repeats (see main text for justification).

#### Reconciliation analyses

Generax ^22^ was used to reconcile the gene tree with the species tree using the undatedDL model using the closest proxies to the best-fit models, selected by IQ-TREE2. Specific rate parameters were estimated for each orthogroup (--family-specific-rates option). Reconciliations were visualised using recphyloXML ^56^.

To account for the uncertainty in non-bilaterian animal relationships we performed the reconciliation using the ctenophores first ^57,58^ and sponges first hypothesis ^59,60^. Similarly, we performed a reconciliation including the non-opisthokont species. However, given the lack (or instability - see above) of monoaminergic genes in ctenophore and sponges and in general outside metazoans (see table S3), the differences between the scenarios were marginal (Supplementary Material).

### Rogue Taxa Analysis

To evaluate the presence of problematic sequences ^15^, we used two orthogonal methods the t-index ^19^ and the Leaf stability index ^20,61^. The t-index evaluates how often the taxa moves position across the bootstrap repeats (i.e., how unstable the sequence is). The Leaf stability index (LSI) uses the quartet frequencies to estimate sequence stability, and it was calculated using RogueNaRok ^21^. The LSI was estimated using the TBE bootstrap trees and the UFB trees independently. To identify unstable taxa, we mainly used the t-index and considered sequences with a score > 2 as unstable. We further validated this instability with LSI where for a species sequences the lower the value the more unstable it is. Where unstable sequences may have an influence on the topology or reconciliation, we removed them from the alignment and re-ran phylogenetic analyses as described above.

## Abbreviations

AAAH: Aromatic Amino Acid Hydroxylase
AADC: Aromatic amine Decarboxylase
TPH: Tryptophan Hydroxylase
TH: Tyrosine Hydroxylase
PAH: Phenylalanine Hydroxylase
DDC: Dopa Decarboxylase
HDC: Histidine Decarboxylase
TDC: Tyrosine Decarboxylase
DBH: Dopamine Beta Hydroxylase
TBH: Tyramine Beta Hydroxylase
MOXD: Monooxygenase DBH-like
PNMT: Phenylethanolamine-N-Methyltransferase
INMT: Indolethylamine-N-Methyltransferase
NNMT: Nicotinamide-N-Methyltransferase
VMAT: Vesicular Monoamine Transporter
VACHT: Vesicular Acetylcholine Transporter
SLC: Solute Ligand Carrier
SERT: Serotonin Transporter
DAT: Dopamine Transporter
IDAT: Invertebrate Dopamine Transporter
NET: Nor-epinephrine Transporter
HRH: Histamine Receptor
ACM: Acetylcholine Muscarinic Receptor
GPCR: G-protein Coupled Receptor
GCH: GTP Cyclo-hydrolase
MAO: Monoamine Oxidase
HNMT: Histamine-N-Methyltransferase
COMT: Catechol-O-Methyltransferase

## Acknowledgements

This study was supported by a Royal Society University Research Fellowship and Grants (UF160226 and RGF\EA\180052) to R.F, by the Swiss National Science Foundation (Grants IZCOZ0_182957 and 310030_188471) to SP, by the European Union’s Horizon 2020 research and innovation programme (Marie Sklodowska-Curie grant agreement No 765937) to ER and by a PhD studentship from the University of Leicester to M.G.

## Authors Contributions

R.F. and M.G. conceived the study. M.G. and G.B. performed the analyses. R.F., M.G., G.B., E.R. and S.S, performed the data interpretation. R.F. and M.G. wrote the main text with the help of G.B., E.R. and S.S.

## Data availability

The data underlying this article and the supplementary figures are available on Figshare https://leicester.figshare.com/account/home?inst=leicester#/projects/144243

## SUPPLEMENTARY FIGURES AND TABLES

**Figure S1**. Analysis of AAAHs. **A)** TBE tree with TBE score supports. **B)** Maximum likelihood tree with ultrafast bootstrap proportion supports. **C)** TBE tree with TBE score supports without rogue sequences. **D)** Maximum likelihood tree with ultrafast bootstrap proportion supports without rogue sequences **E)** reconciliation using sponge-first hypothesis **F)** reconciliation using ctenophore-first hypothesis.

**Figure S2**. Evolution of GCHs. **A)** Transfer bootstrap estimation (TBE) tree. **B)** Simplified illustration of gene tree to species tree reconciliation. Support shown are TBE scores (in bold) and ultrafast bootstrap proportions for key nodes.

**Figure S3**. Analysis of GCHs. **A)** TBE tree with TBE score supports. **B)** Maximum likelihood tree with ultrafast bootstrap proportion supports. **C)** Reconciliation using sponge-first hypothesis **D)** Reconciliation using ctenophore-first hypothesis.

**Figure S4**. Analysis of AADCs. **A)** TBE tree with TBE score supports. **B)** Maximum likelihood tree with ultrafast bootstrap proportion supports. **C)** TBE tree with TBE score supports without rogue sequences. **D)** Maximum likelihood tree with ultrafast bootstrap proportion supports without rogue sequences **E)** Reconciliation using sponge-first hypothesis **F)** Reconciliation using ctenophore-first hypothesis.

**Figure S5**. Analysis of BHs. **A)** TBE tree with TBE score supports. **B)** Maximum likelihood tree with ultrafast bootstrap proportion supports. **C)** UFB tree reconciliation using sponge-first hypothesis **D)** UFB tree reconciliation using ctenophore-first hypothesis. **E)** TBE tree reconciliation using sponge-first hypothesis **F)** TBE tree reconciliation using ctenophore-first hypothesis.

**Figure S6**. Analysis of PNMTs. **A)** TBE tree with TBE score supports. **B)** Maximum likelihood tree with ultrafast bootstrap proportion supports. **C)** Reconciliation using sponge-first hypothesis **D)** reconciliation using ctenophore-first hypothesis.

**Figure S7**. Analysis of VMATs. **A)** TBE tree with TBE score supports. **B)** Maximum likelihood tree with ultrafast bootstrap proportion supports. **C)** TBE tree with TBE score supports without rogue sequences. **D)** Maximum likelihood tree with ultrafast bootstrap proportion supports without rogue sequences **E)** Reconciliation using sponge-first hypothesis **F)** Reconciliation using ctenophore-first hypothesis.

**Figure S8**. Analysis of SLC6 sequences. **A)** TBE tree with TBE score supports. **B)** Maximum likelihood tree with ultrafast bootstrap proportion supports. **C)** Reconciliation using sponge-first hypothesis **D)** Reconciliation using ctenophore-first hypothesis.

**Figure S9**. Evolution of COMTs. **A)** Transfer bootstrap estimation (TBE) tree. **B)** Simplified illustration of the gene tree to species tree reconciliation. Supports shown are TBE scores (in bold) and ultrafast bootstrap proportions for key nodes.

**Figure S10**. Analysis of COMTs. **A)** TBE tree with TBE score supports. **B)** Maximum likelihood tree with ultrafast bootstrap proportion supports. **C)** Reconciliation using sponge-first hypothesis **D)** Reconciliation using ctenophore-first hypothesis.

**Figure S11**. Evolution of HNMTs. **A)** Transfer bootstrap estimation (TBE) tree. **B)** Simplified illustration of the gene tree to species tree reconciliation. Supports shown are TBE scores (in bold) and ultrafast bootstrap proportions for key nodes.

**Figure S12**. Analysis of HNMTs. **A)** TBE tree with TBE score supports. **B)** Maximum likelihood tree with ultrafast bootstrap proportion supports. **C)** Reconciliation using sponge-first hypothesis **D)** Reconciliation using ctenophore-first hypothesis.

**Figure S13**. Evolution of MAOs. **A)** Transfer bootstrap estimation (TBE) tree. **B)** Simplified illustration of the gene tree to species tree reconciliation. Supports shown are TBE scores (in bold) and ultrafast bootstrap proportions for key nodes.

**Figure S14**. Analysis of MAOs. **A)** TBE tree with TBE score supports. **B)** Maximum likelihood tree with ultrafast bootstrap proportion supports. **C)** Reconciliation using sponge-first hypothesis **D)** Reconciliation using ctenophore-first hypothesis.

**Figure S15**. Evolution of 5HT3Rs. **A)** Transfer bootstrap estimation (TBE) tree. **B)** Simplified illustration of the gene tree to species tree reconciliation. Supports shown are TBE scores (in bold) and ultrafast bootstrap proportions for key nodes.

**Figure S16**. Analysis of 5HT3Rs. **A)** TBE tree with TBE score supports. **B)** Maximum likelihood tree with ultrafast bootstrap proportion supports. **C)** Reconciliation using sponge-first hypothesis **D)** Reconciliation using ctenophore-first hypothesis.

**Figure S17**. Analysis of OG1. **A)** TBE tree with TBE score supports. **B)** Maximum likelihood tree with ultrafast bootstrap proportion supports. **C)** Reconciliation using sponge-first hypothesis **D)** Reconciliation using ctenophore-first hypothesis.

**Figure S18**. Analysis of OG30. **A)** TBE tree with TBE score supports. **B)** Maximum likelihood tree with ultrafast bootstrap proportion supports. **C)** Reconciliation using sponge-first hypothesis **D)** Reconciliation using ctenophore-first hypothesis.

**Figure S19**. CLANs analysis of GPCRs. **A)** Clusters obtained using a p-value < 1e-100. **B)** Clusters obtained using a p-value < 1e-42 C) Clusters obtained using a p-value < 1e-42 and the connection obtained with a p-value < 1e-40. Each dot represents a sequence, lines indicate similarity between two sequences at or above the p-value threshold. Key clusters were coloured according to sequences from *Homo sapiens, Mus musculus* and *Drosophila melanogaster*. Purple and green indicate bilaterian sequences from OG1 and OG30 respectively.

**Figure S20**. Analysis of GPCRs. **A)** TBE tree with TBE score supports. **B)** Maximum likelihood tree with ultrafast bootstrap proportion supports. **C)** UFB tree reconciliation using sponge-first hypothesis **D)** UFB tree reconciliation using ctenophore-first hypothesis. **E)** TBE tree reconciliation using sponge-first hypothesis **F)** TBE tree reconciliation using ctenophore-first hypothesis.

**Table S1**. Molecular evidence for the presence of monoamines in non-bilaterian animals.

**Table S2**. Species used in this study and their BUSCO completeness scores.

**Table S3**. Distribution of sequences per orthogroups according to Broccoli.

**Table S4**. Query sequence sources, InterProScan annotation motifs and phylogenetic models used for the phylogenetic inferences.

**Table S5**. T-index and Leaf Instability Scores for each gene family.

