## Supplementary_FigureS1-S20 for "Neuromodulation by Monoamines is a Bilaterian Innovation"

S1A.

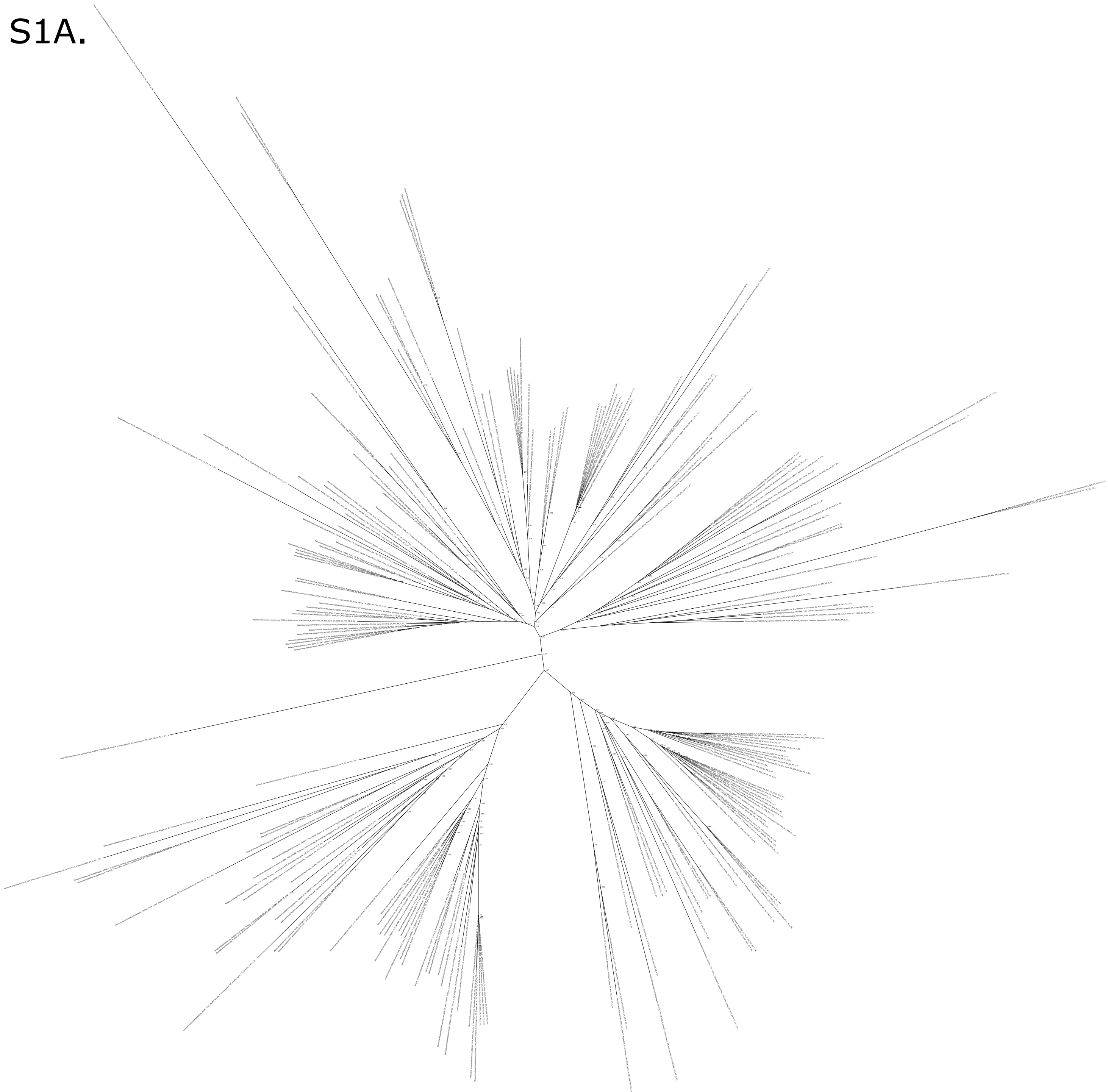

S1B.

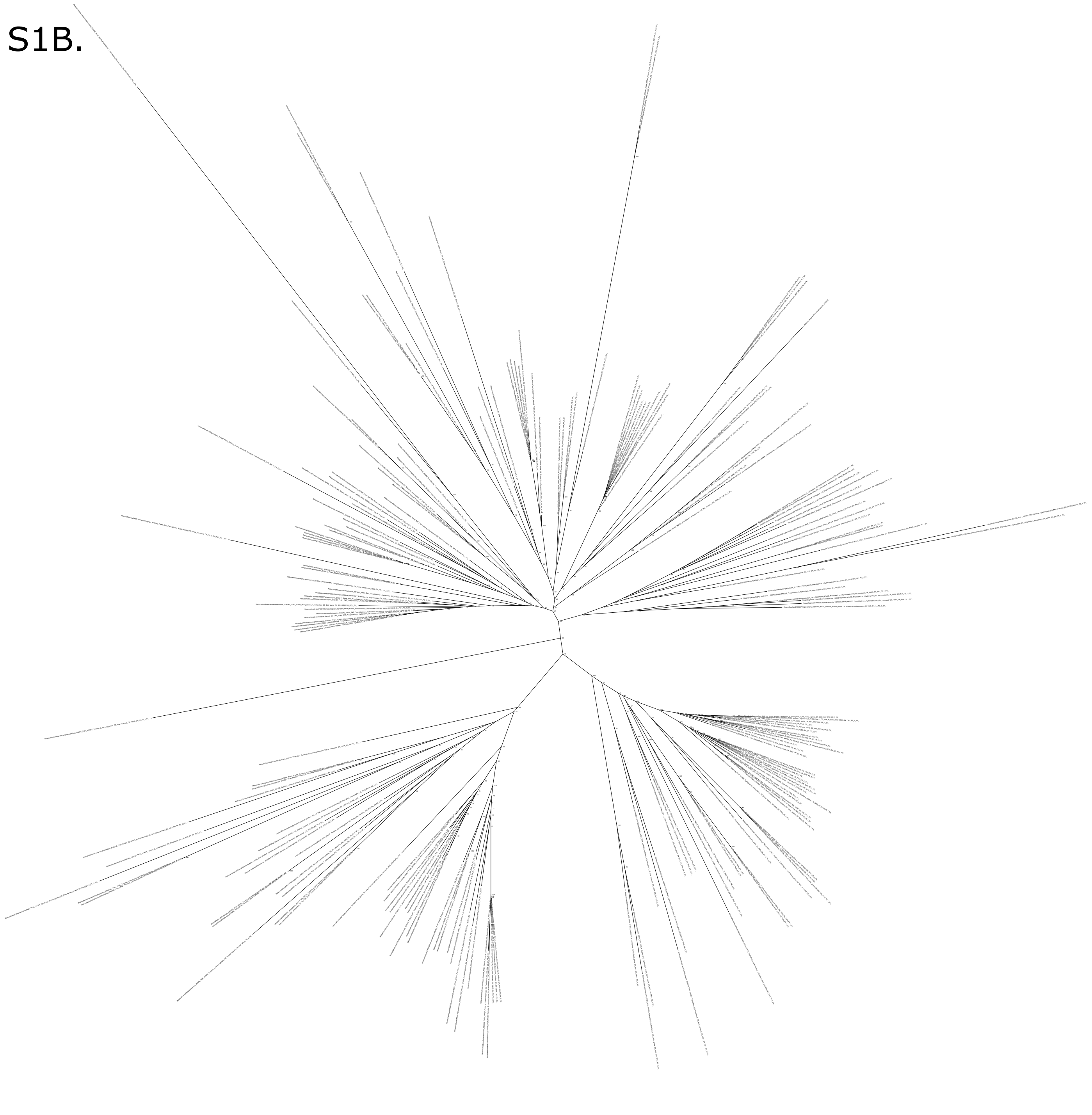

S1C.

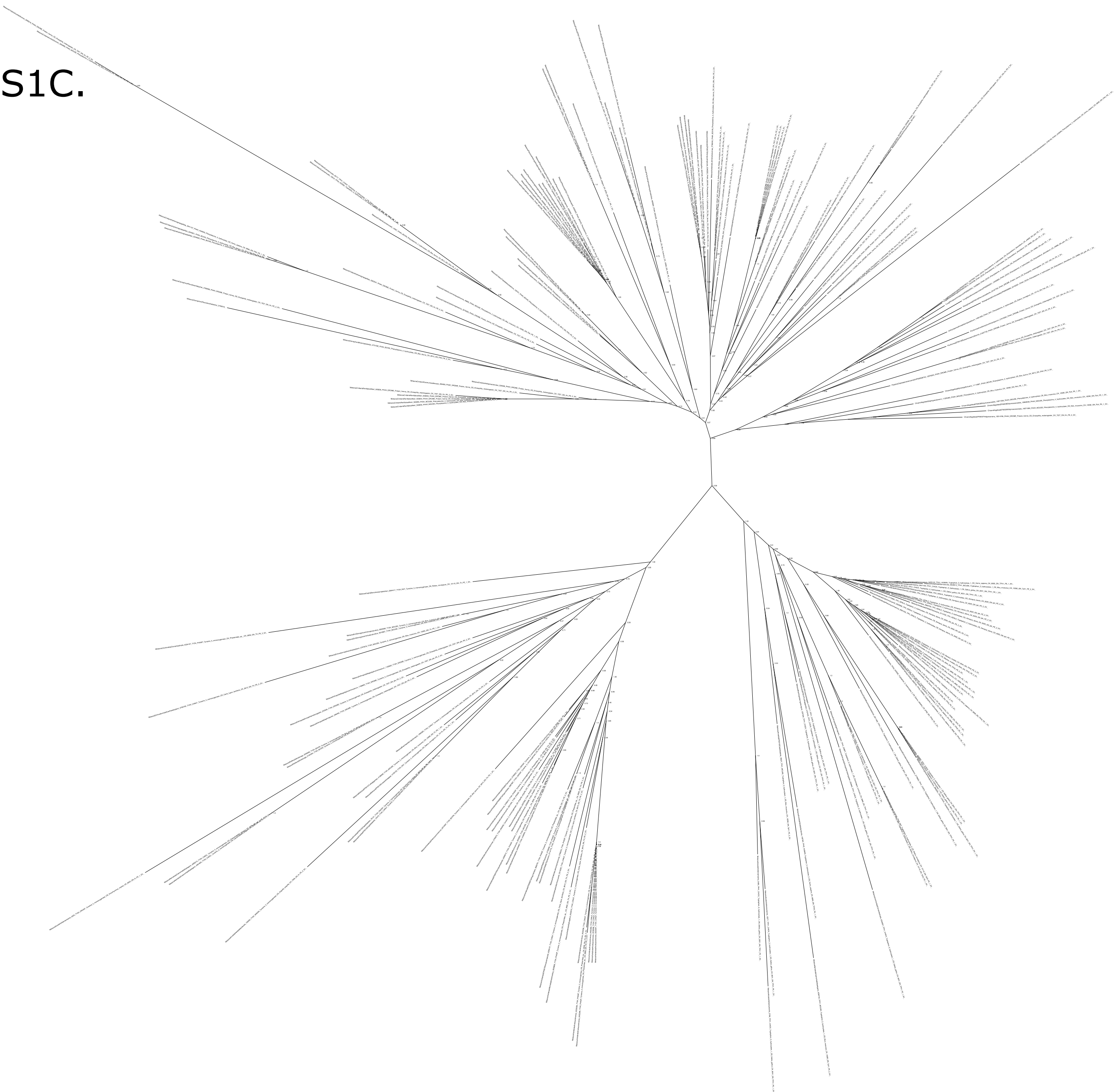

S1D.

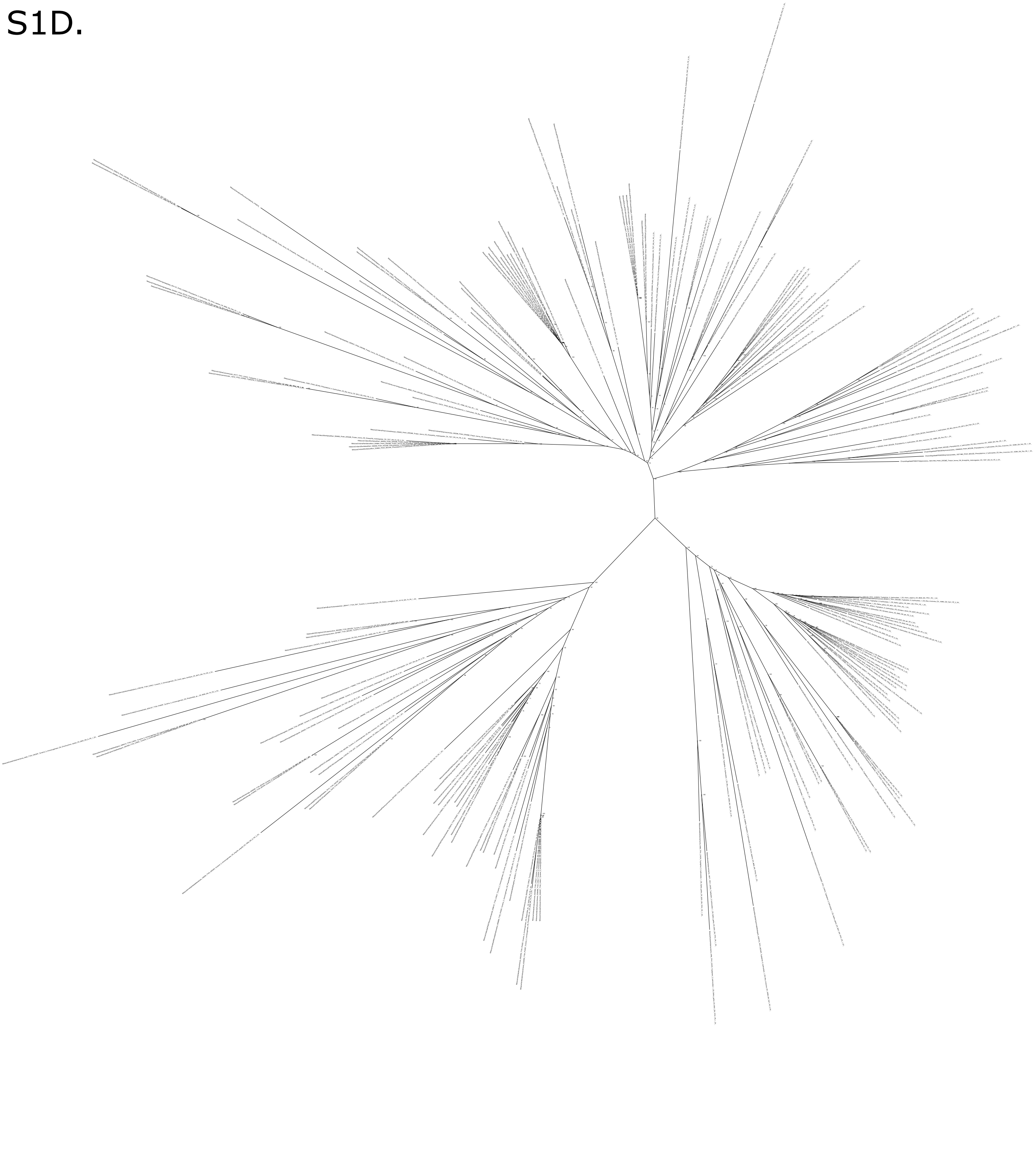

S1E.

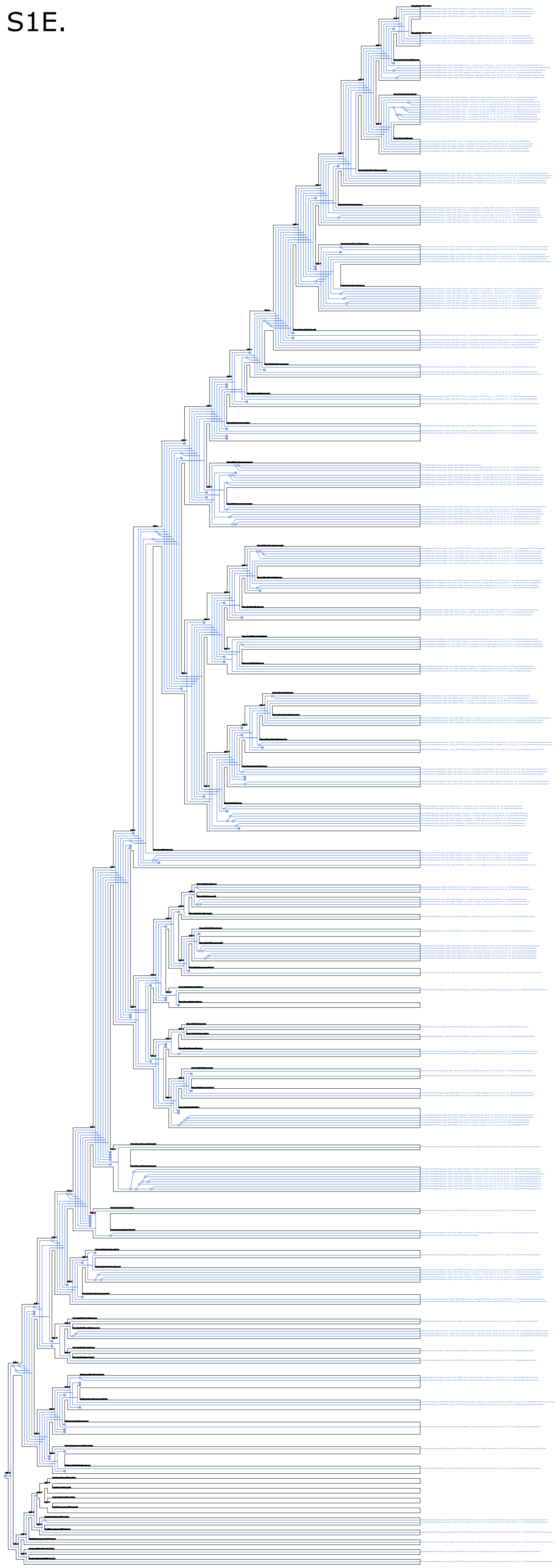

S1F

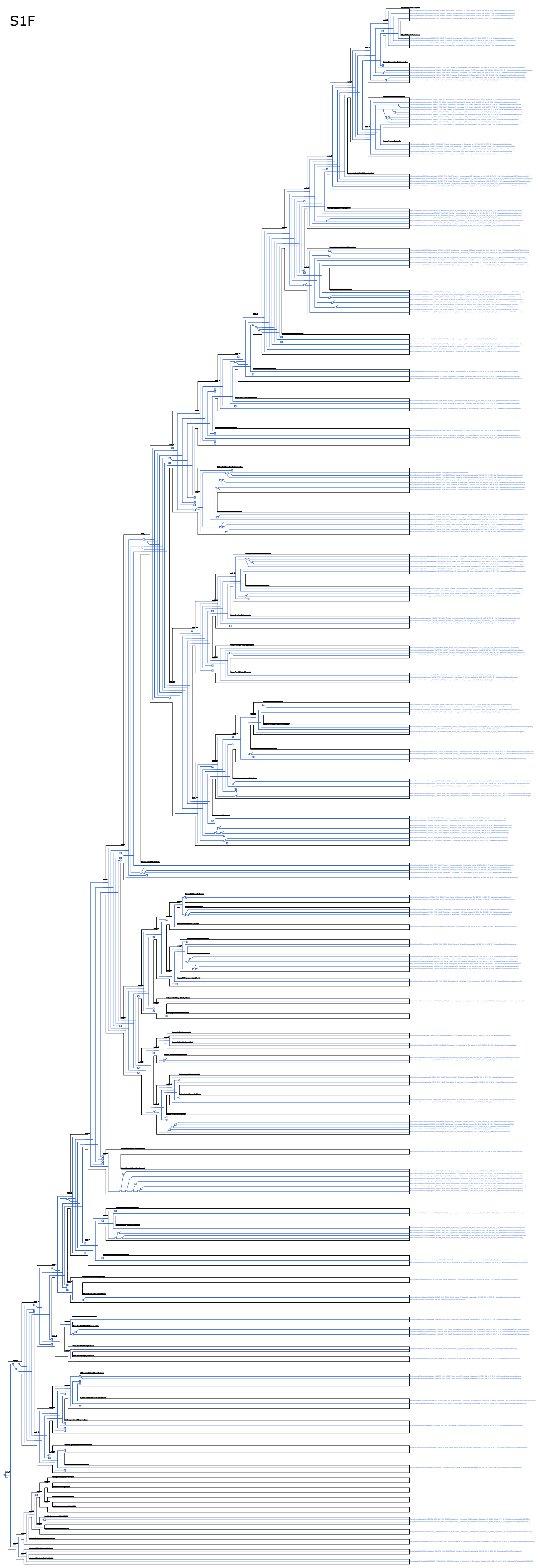

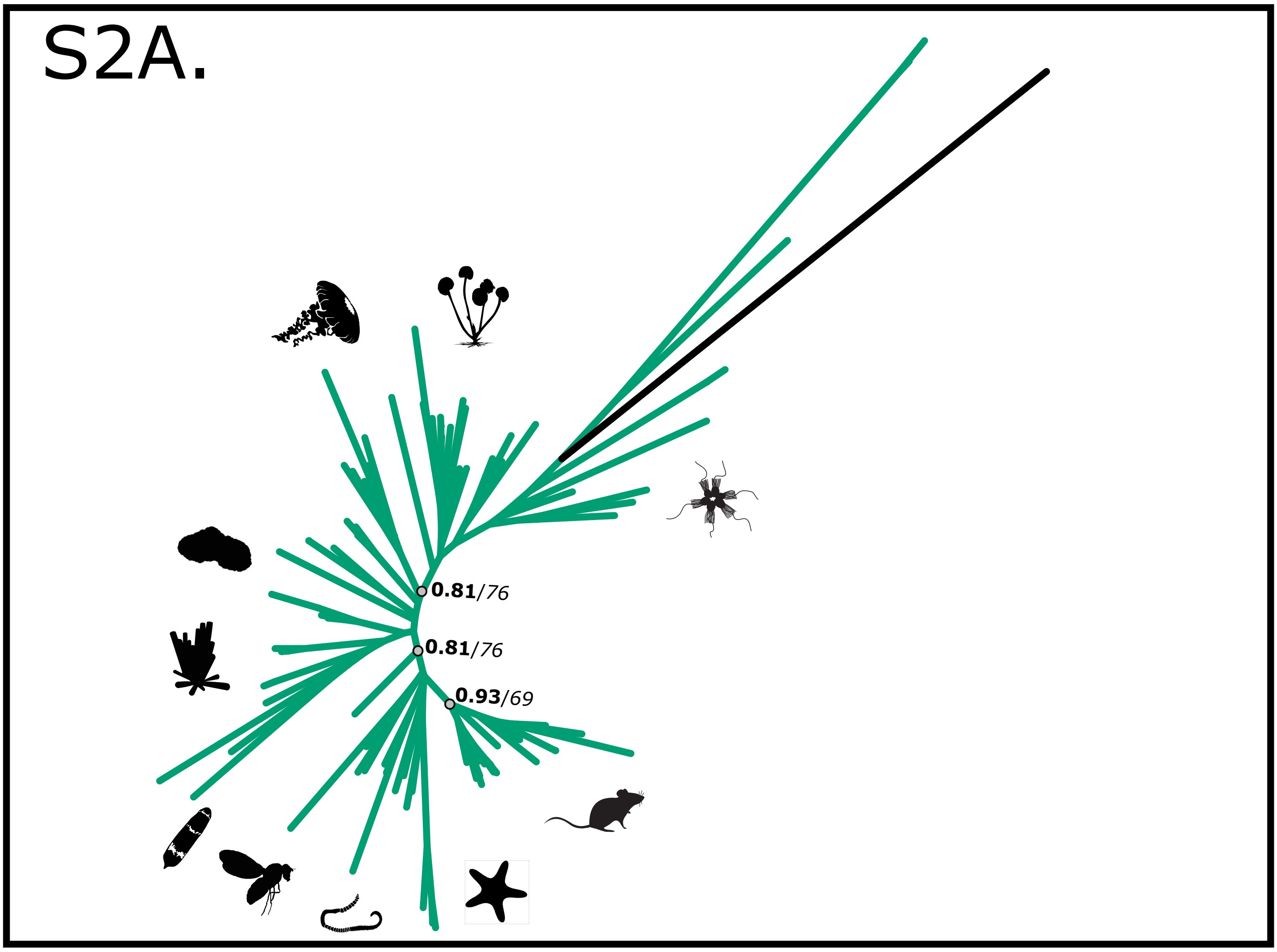
S2A.
0.81/76
0.81/76
0.93/69

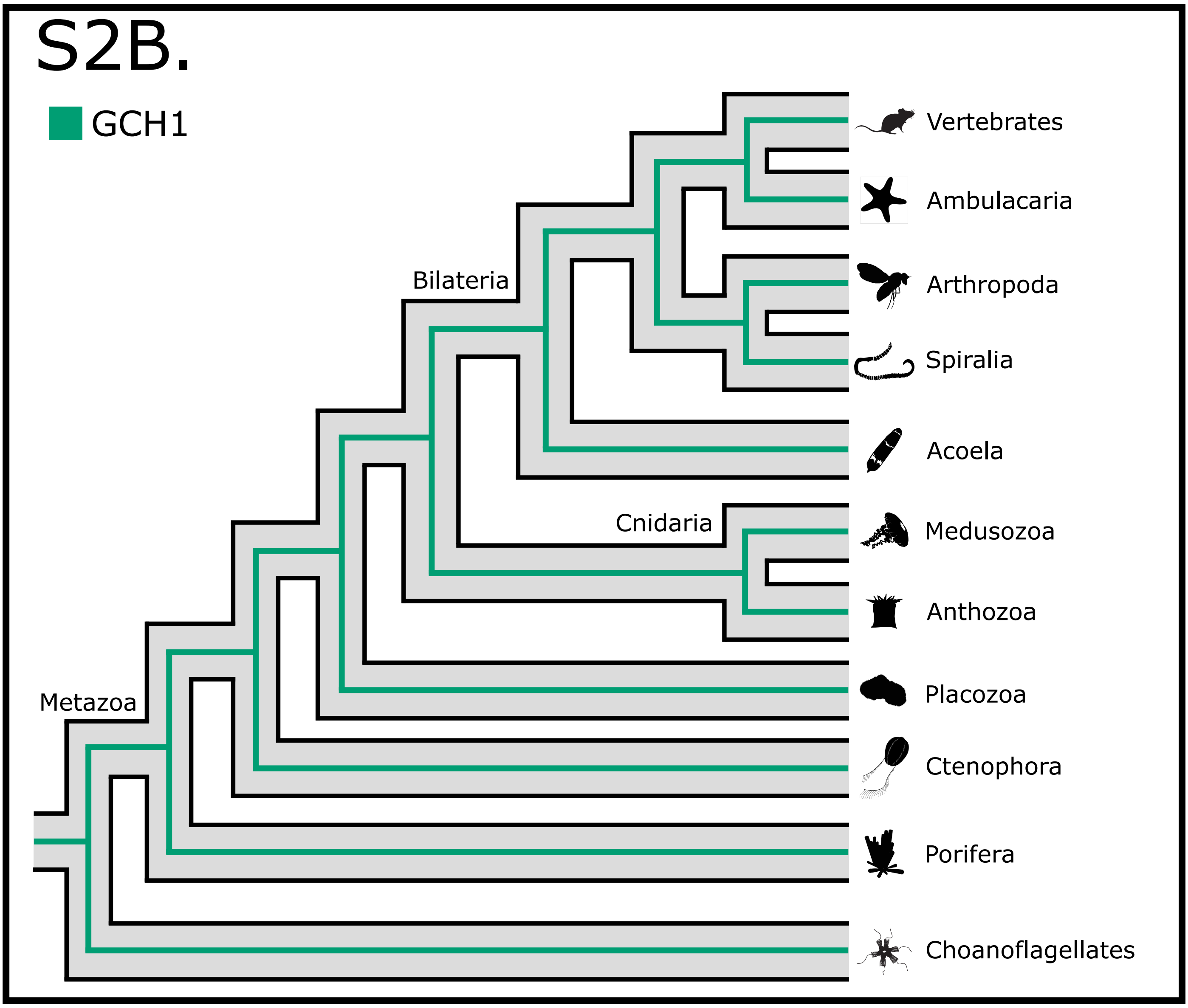
S2B.
GCH1
Bilateria
Cnidaria
Metazoa
Vertebrates
Ambulacaria
Arthropoda
Spiralia
Acoela
Medusozoa
Anthozoa
Placozoa
Ctenophora
Porifera
Choanoflagellates

S3A.

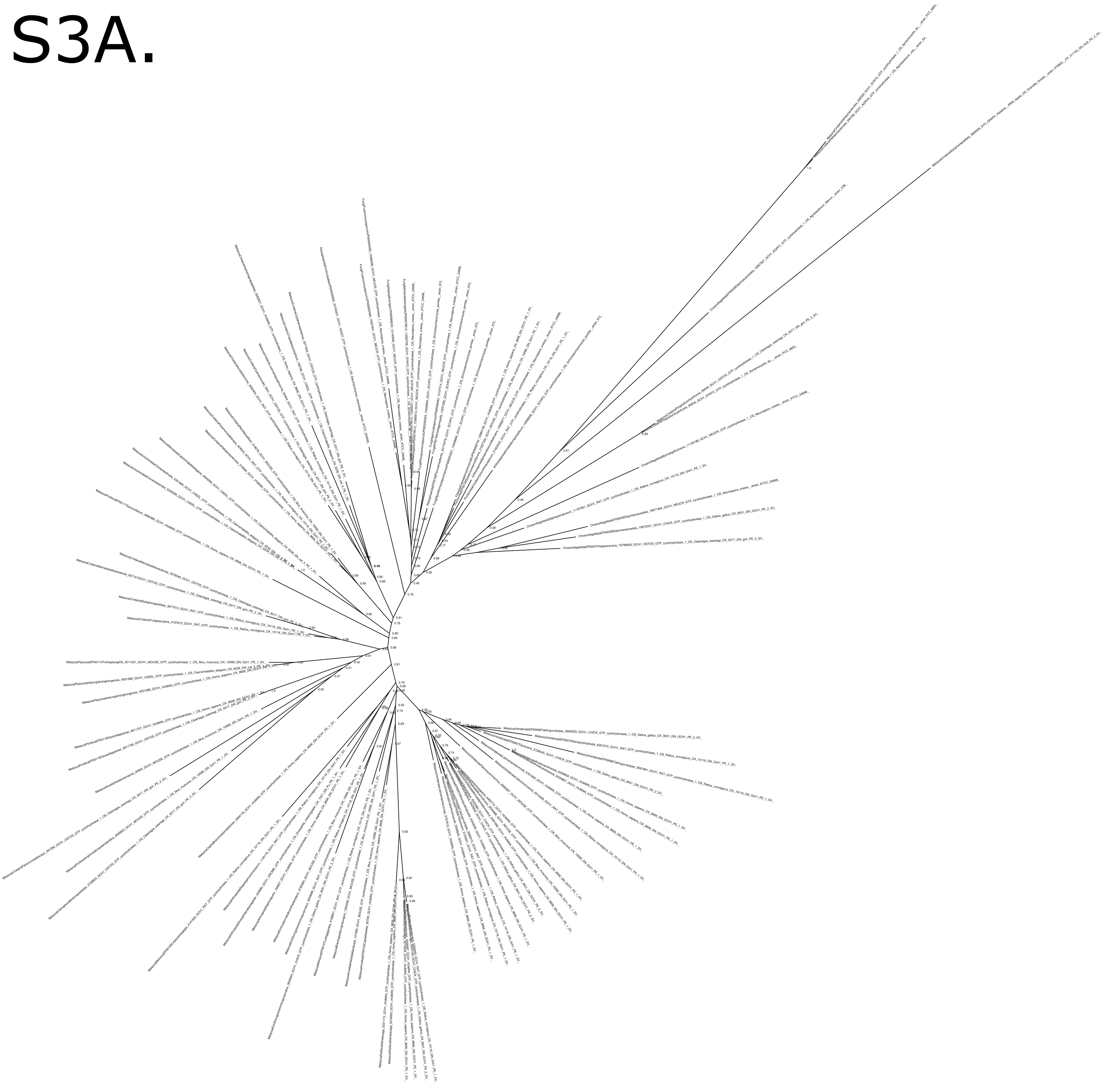

S3B.

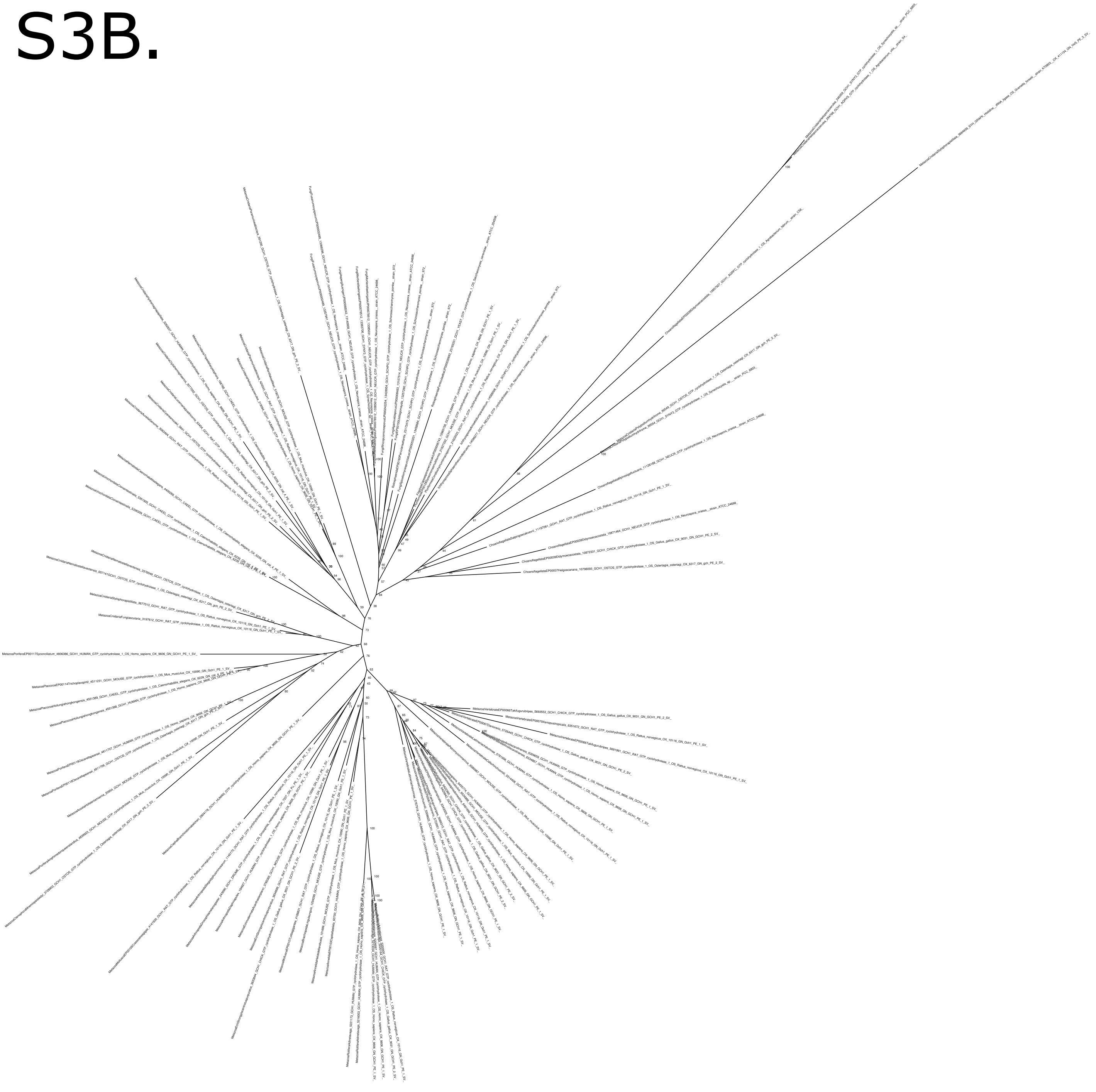

S3C.

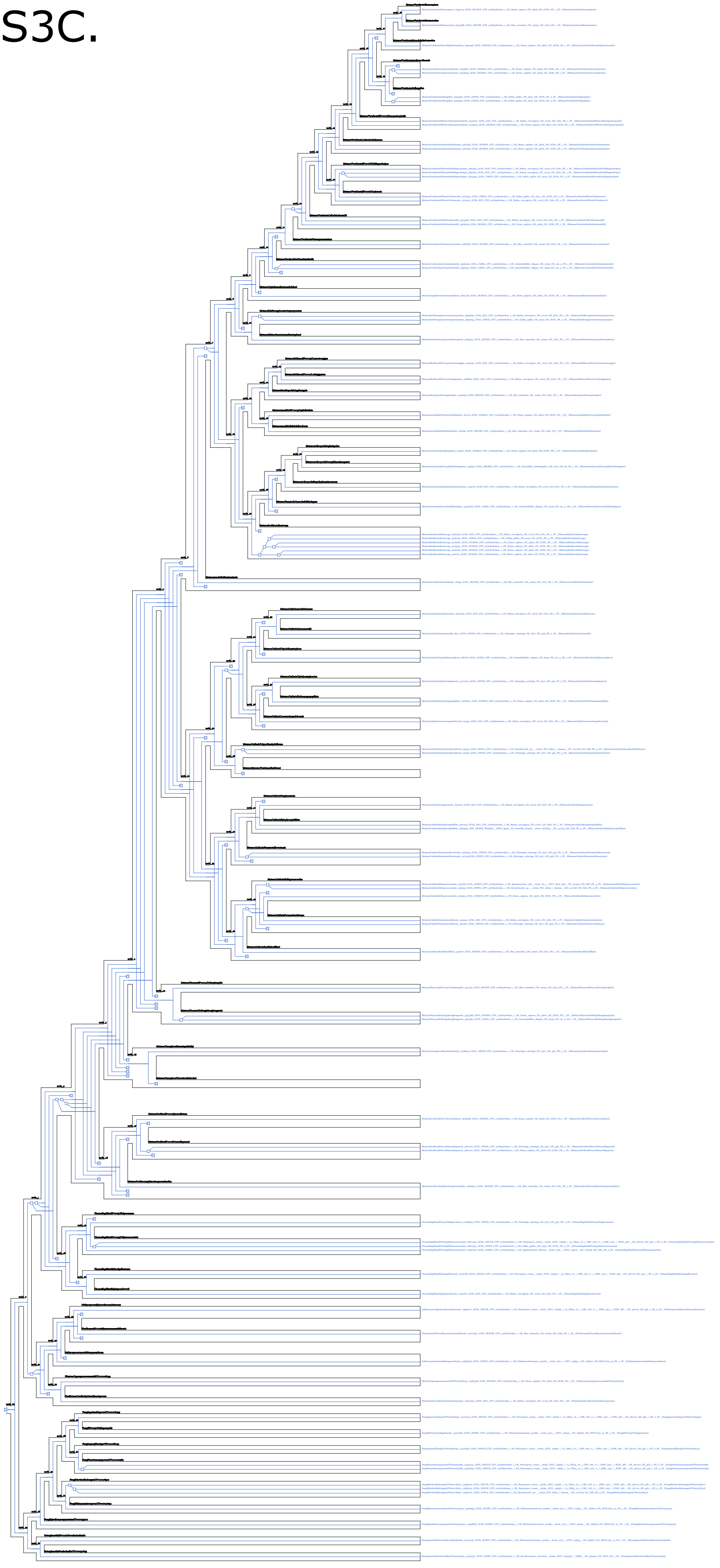

S3D

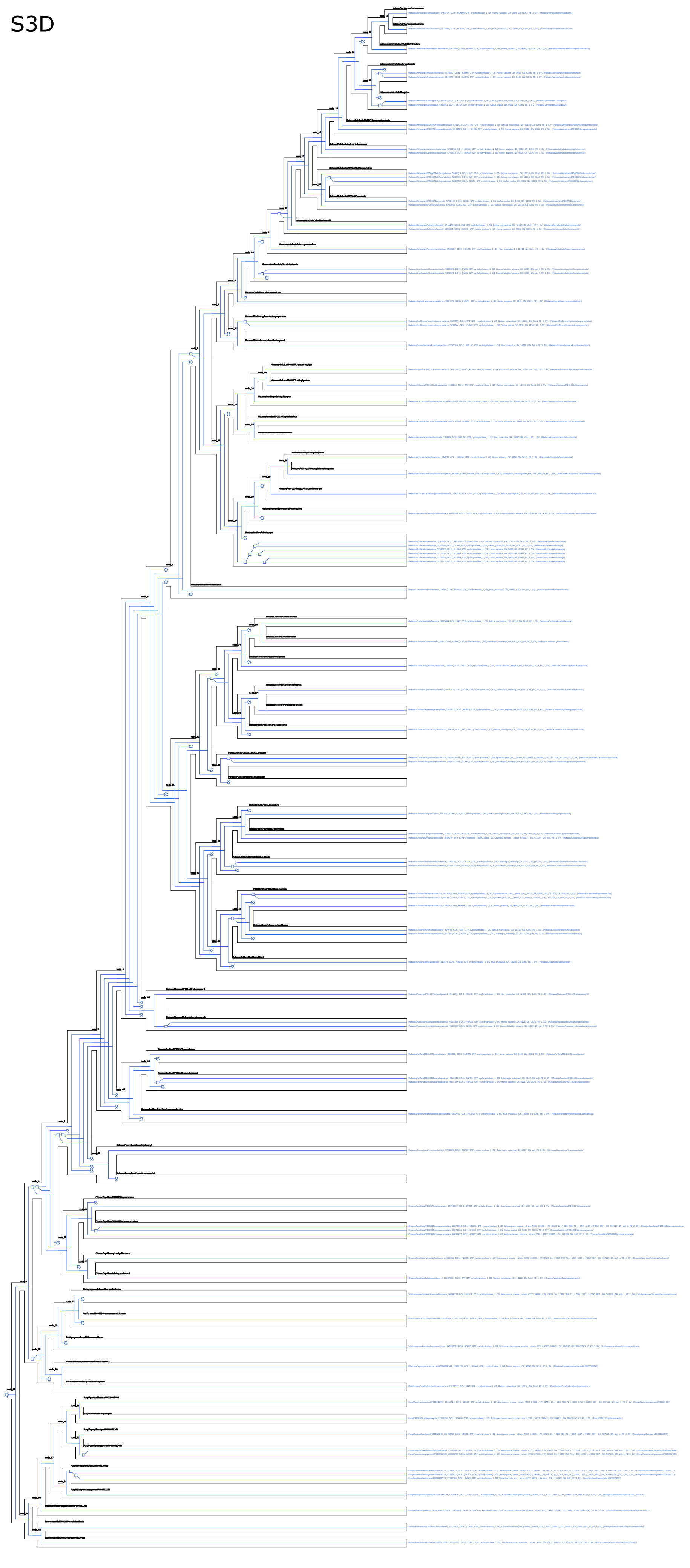

S4A.

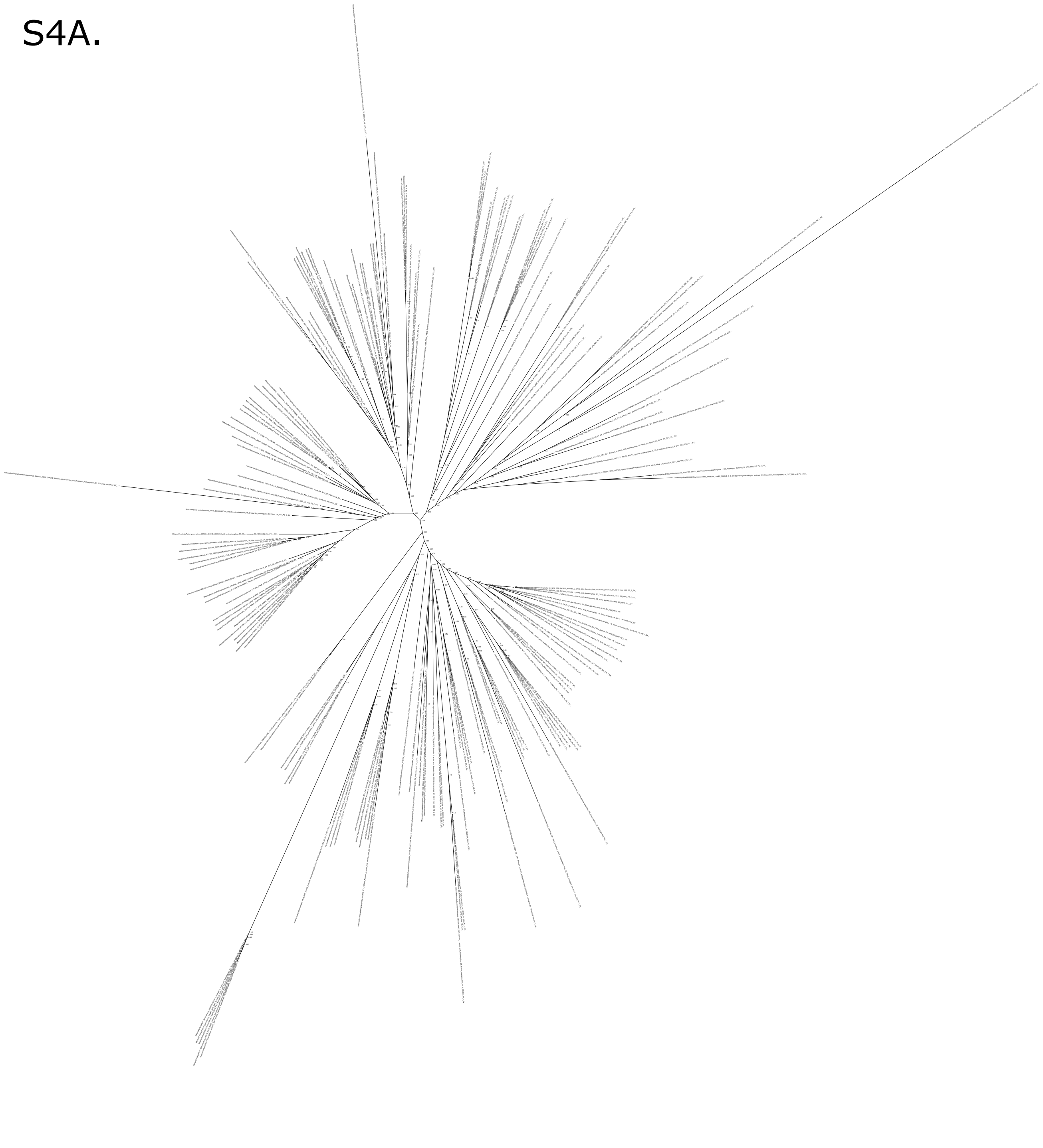

S4B.

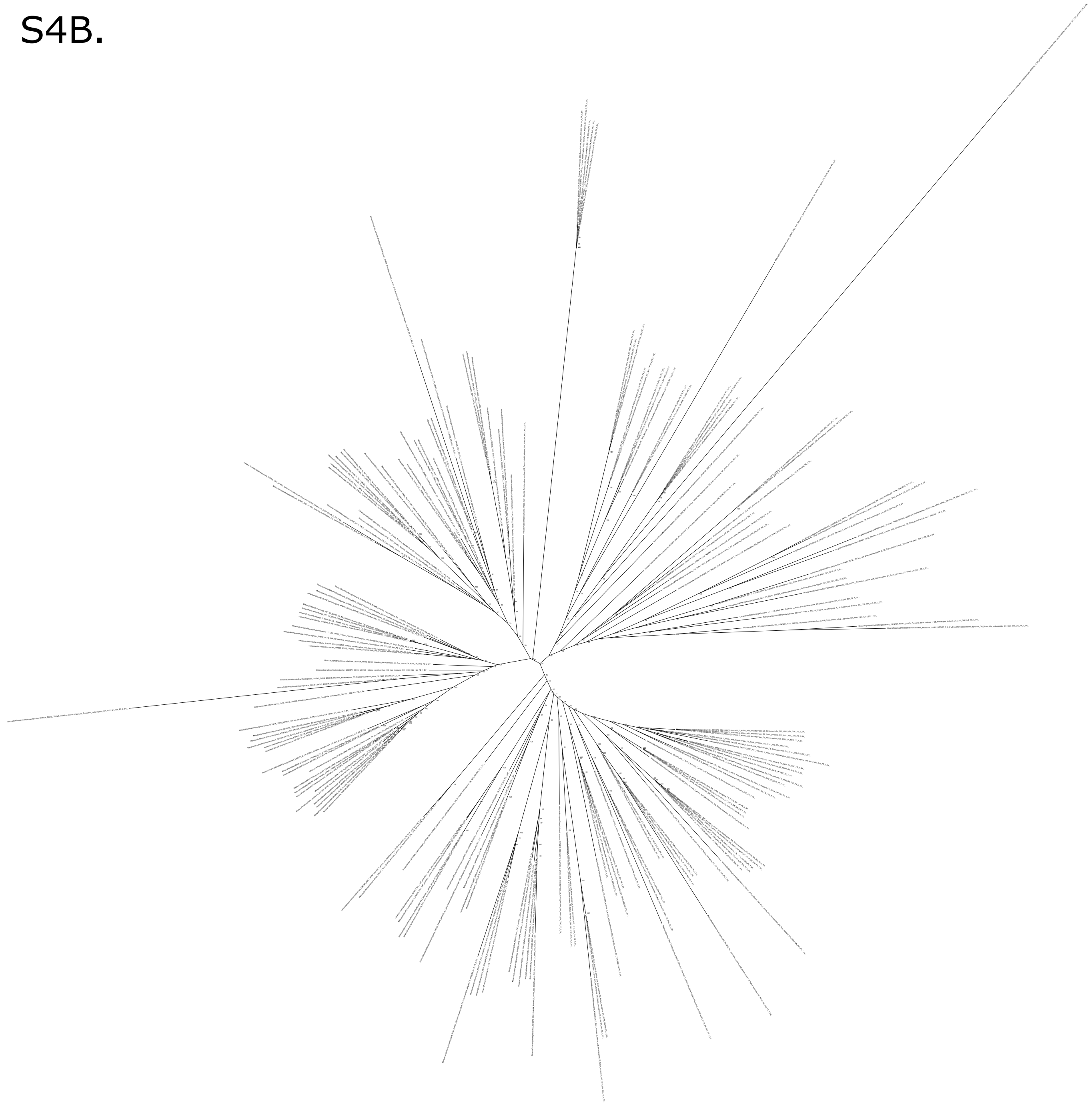

S4C.

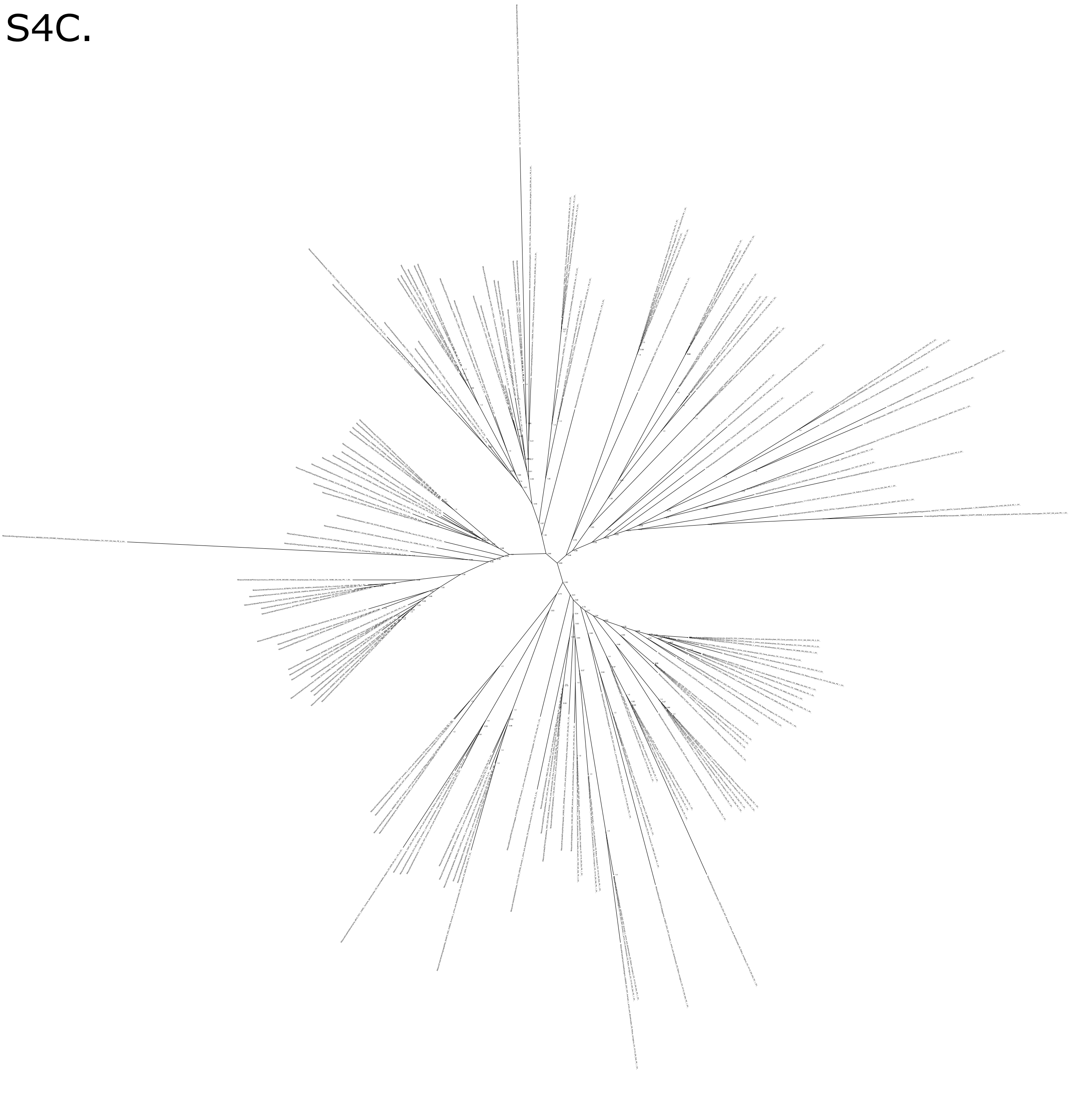

S4D.

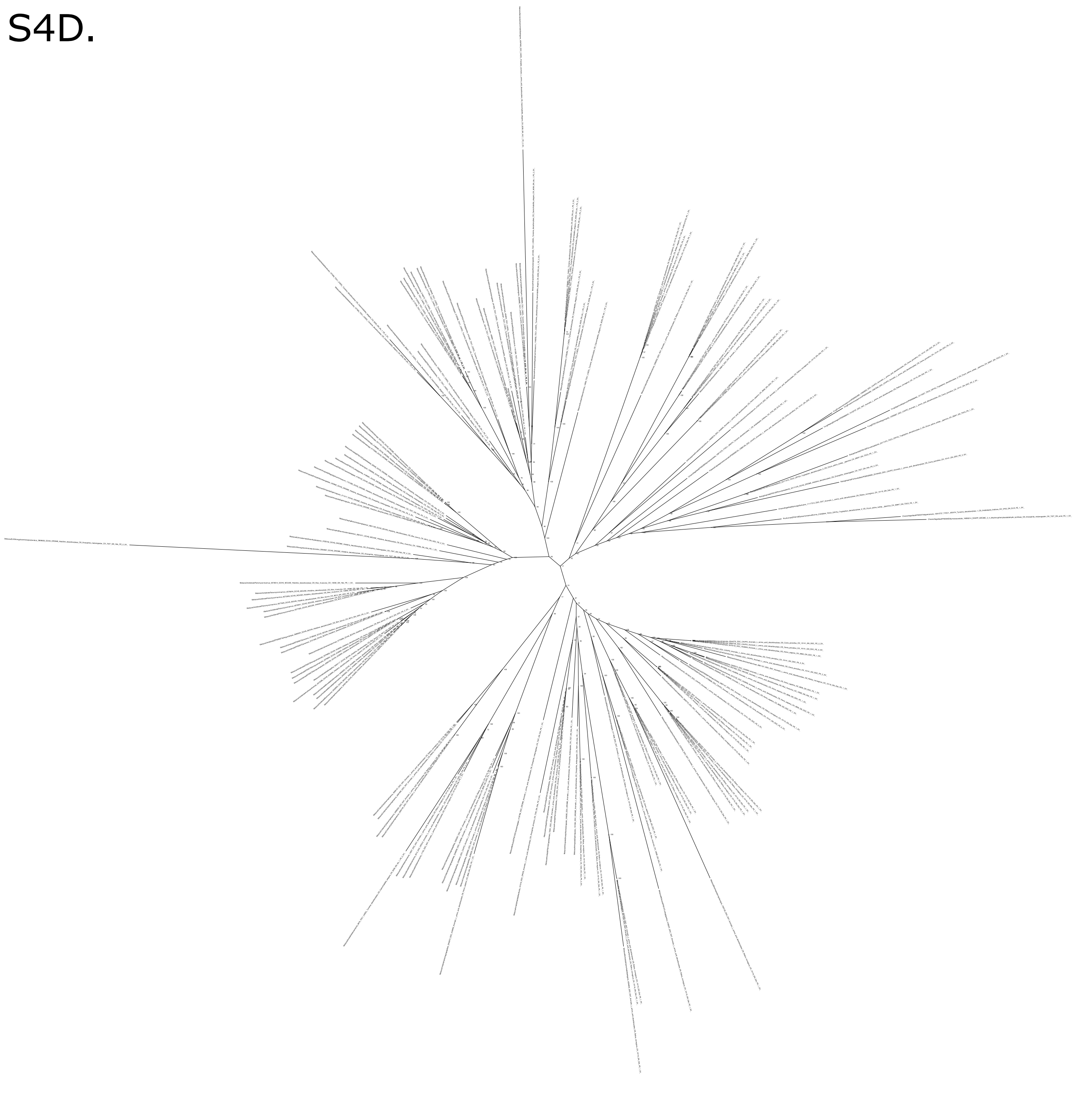

S4E.

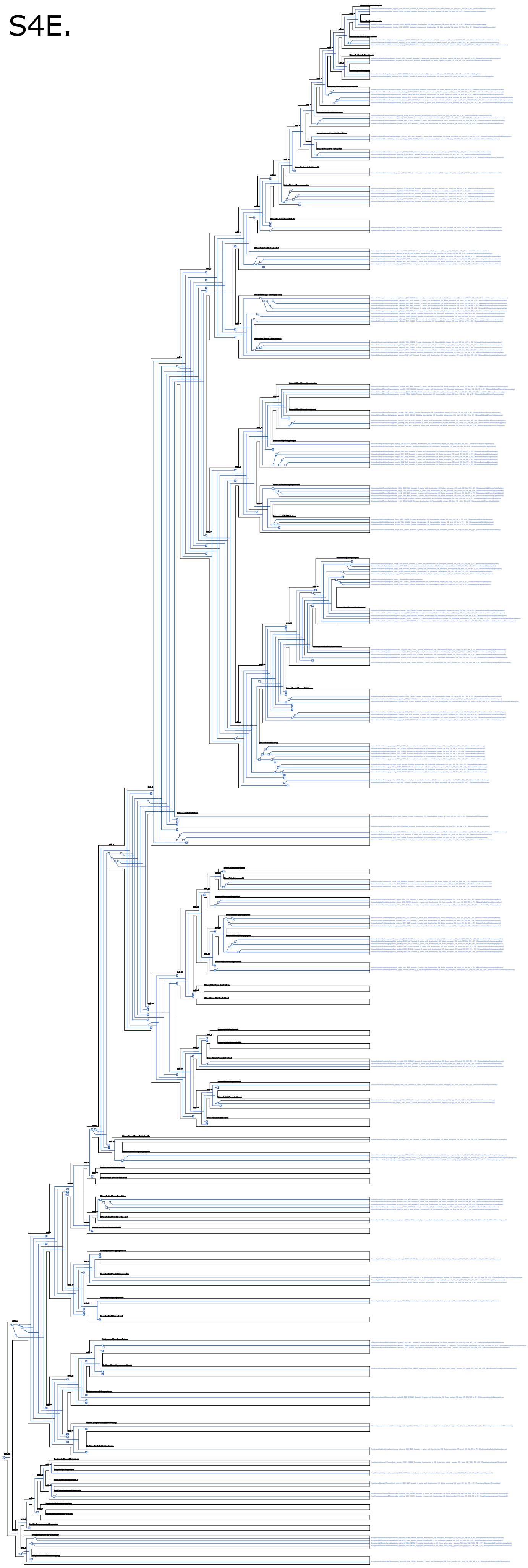

S4F

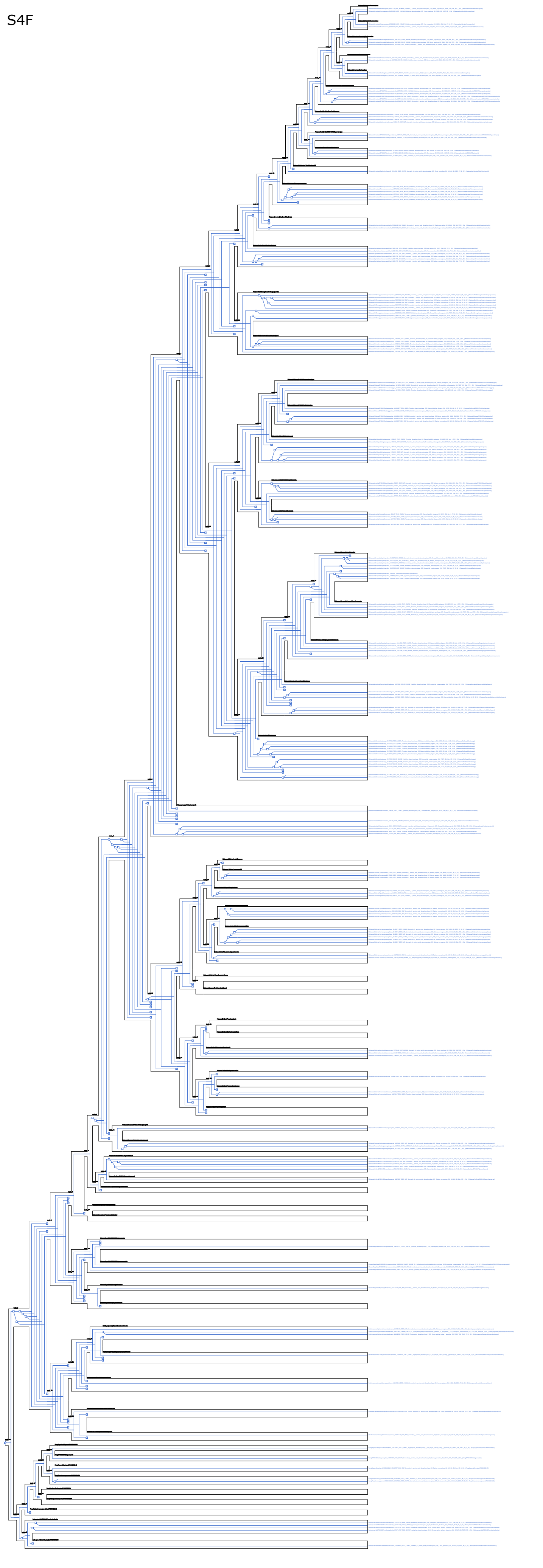

S5A.

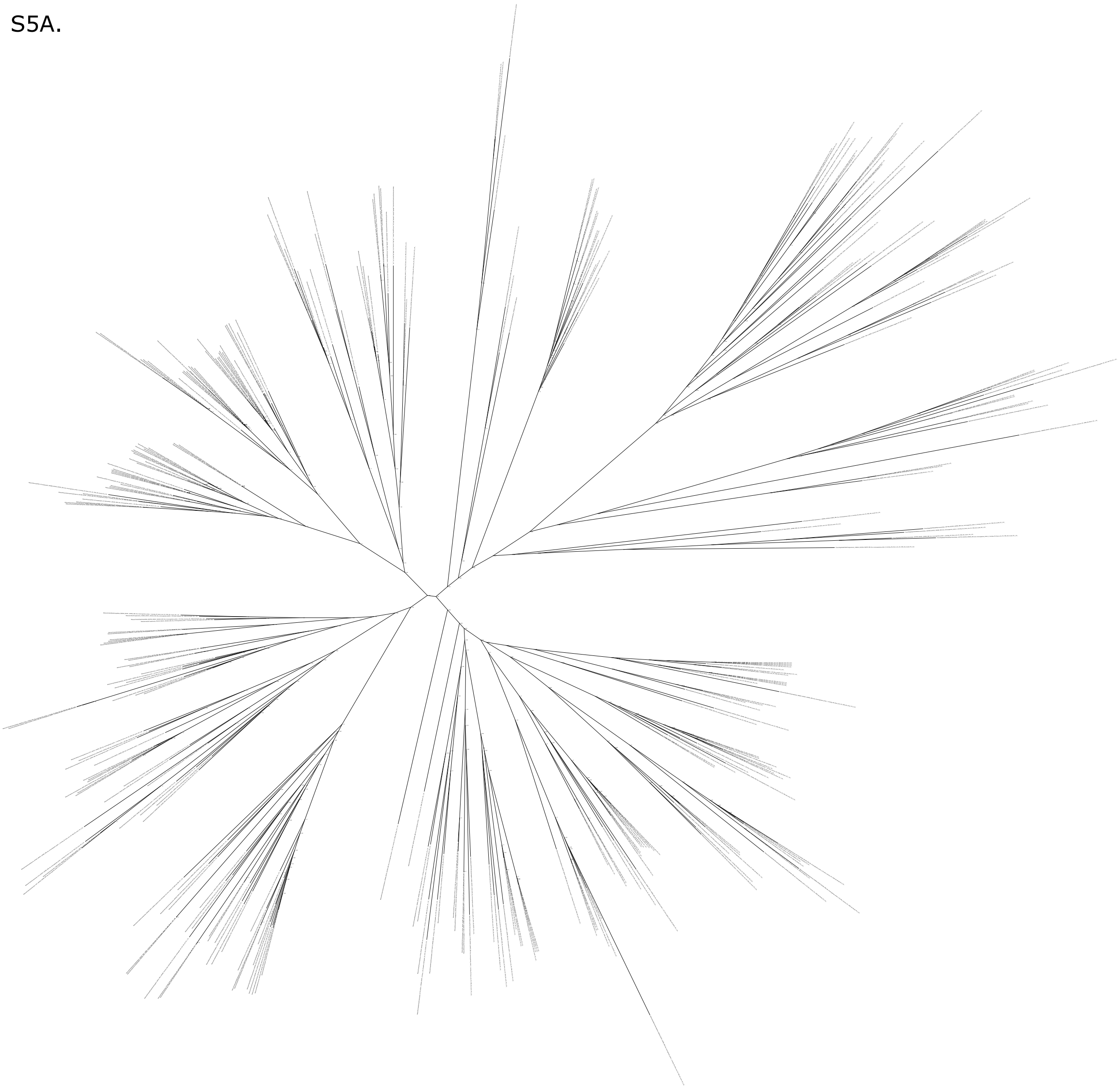

S5B.

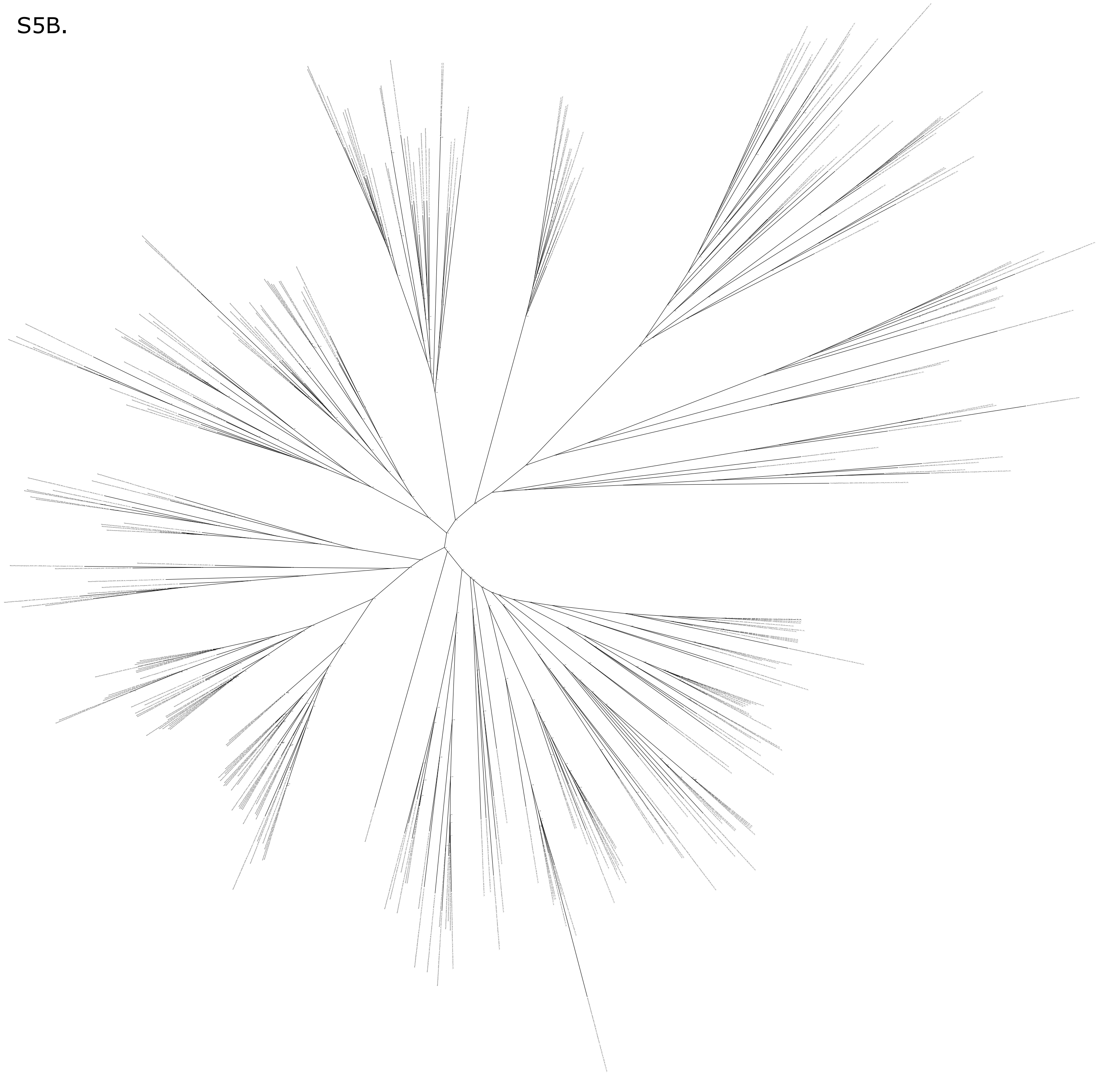

S5C.

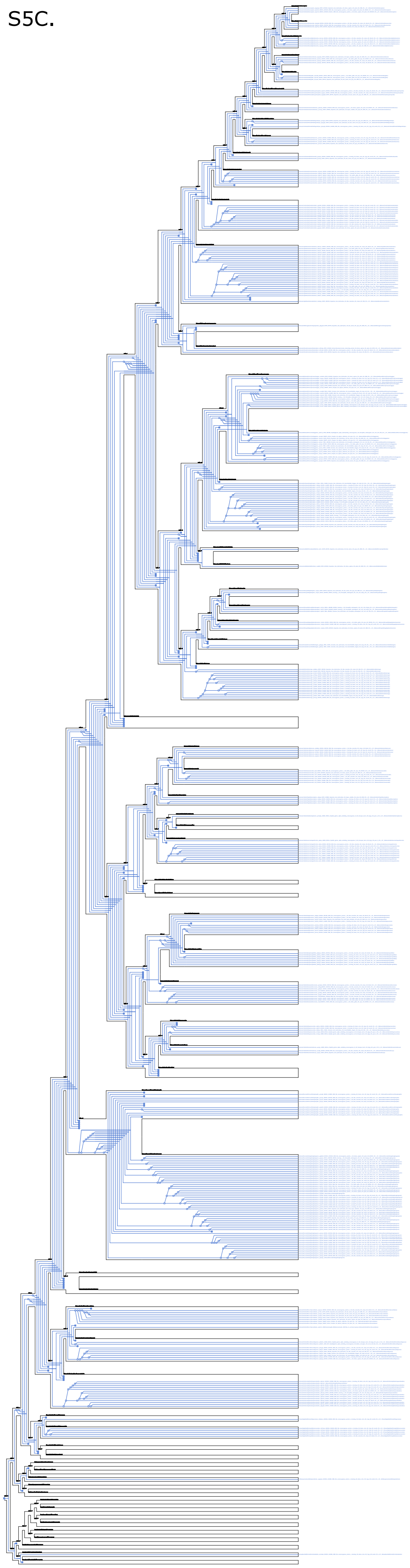

S5D

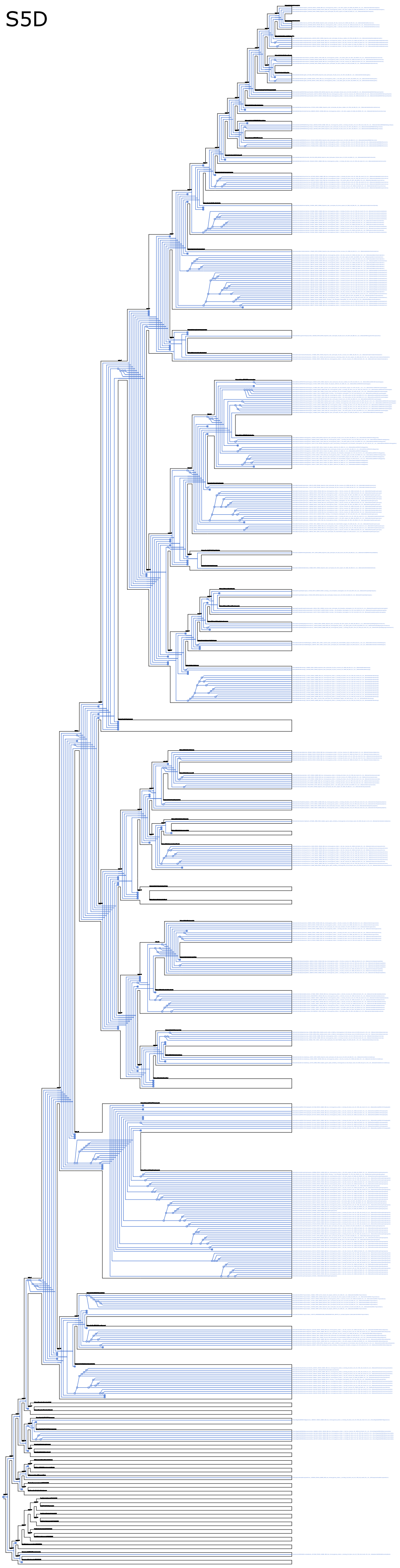

S5E.

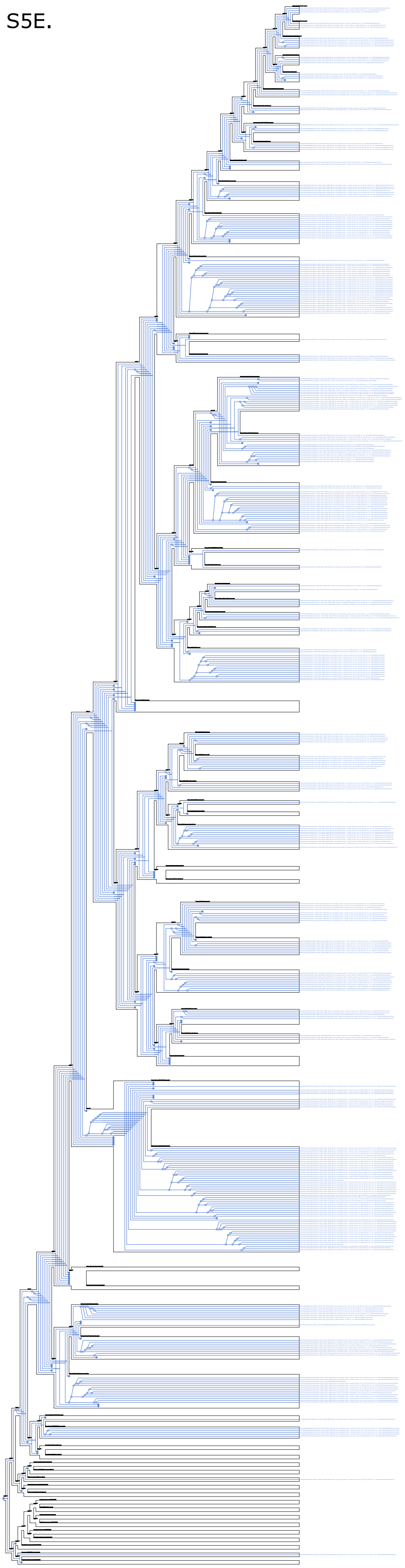

S5F

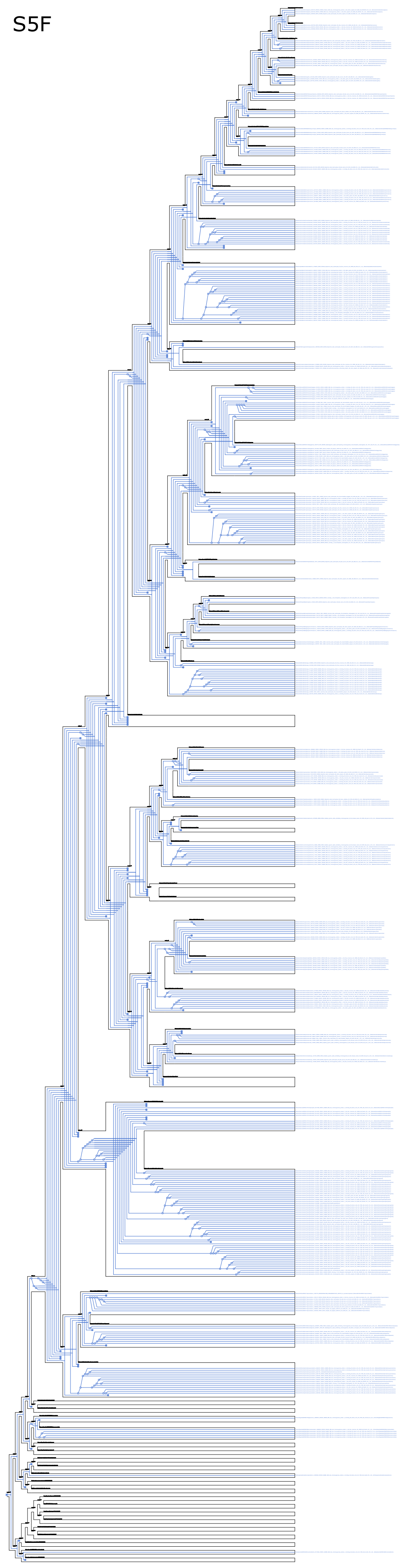

S6A.

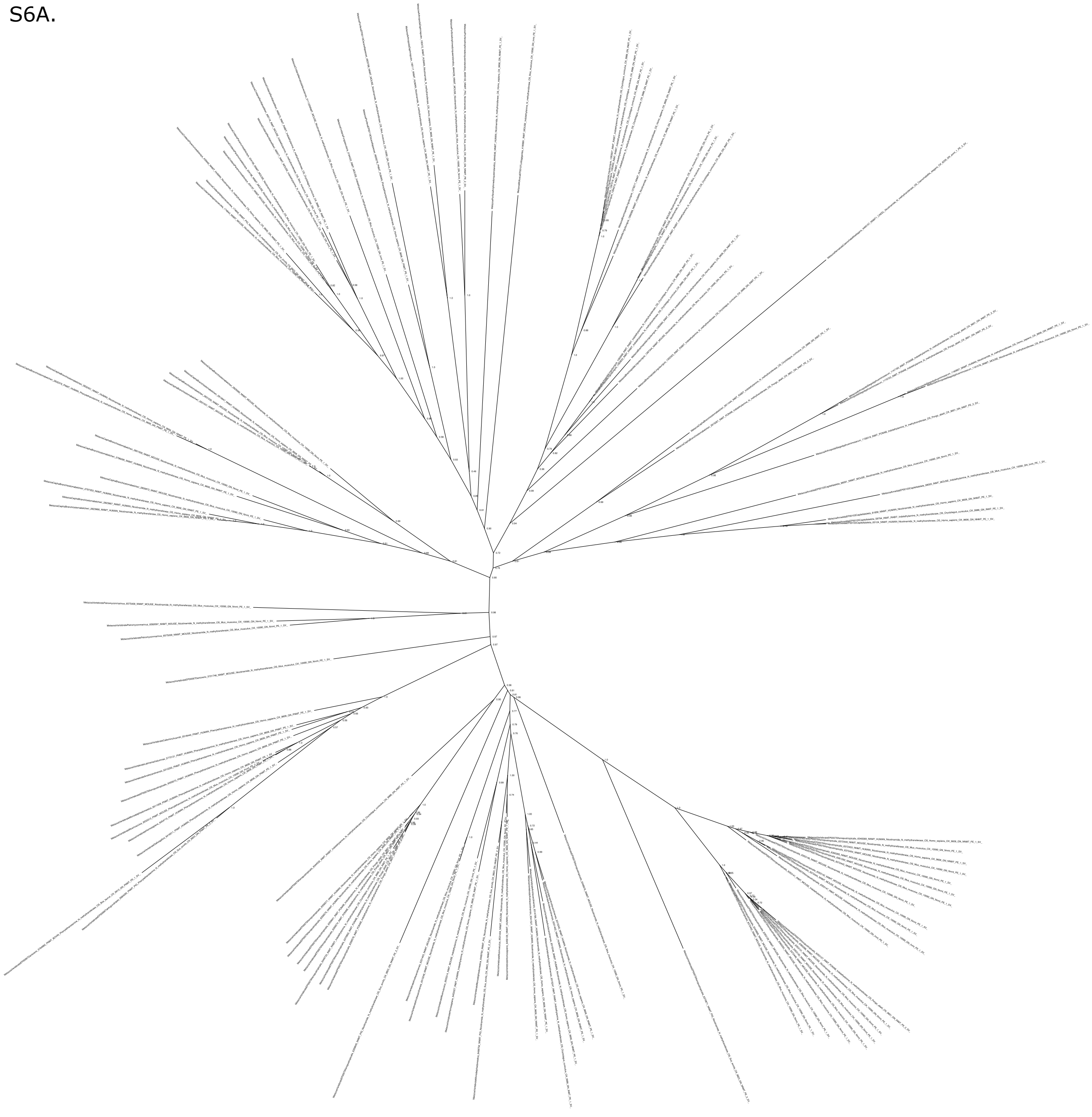

S6B.

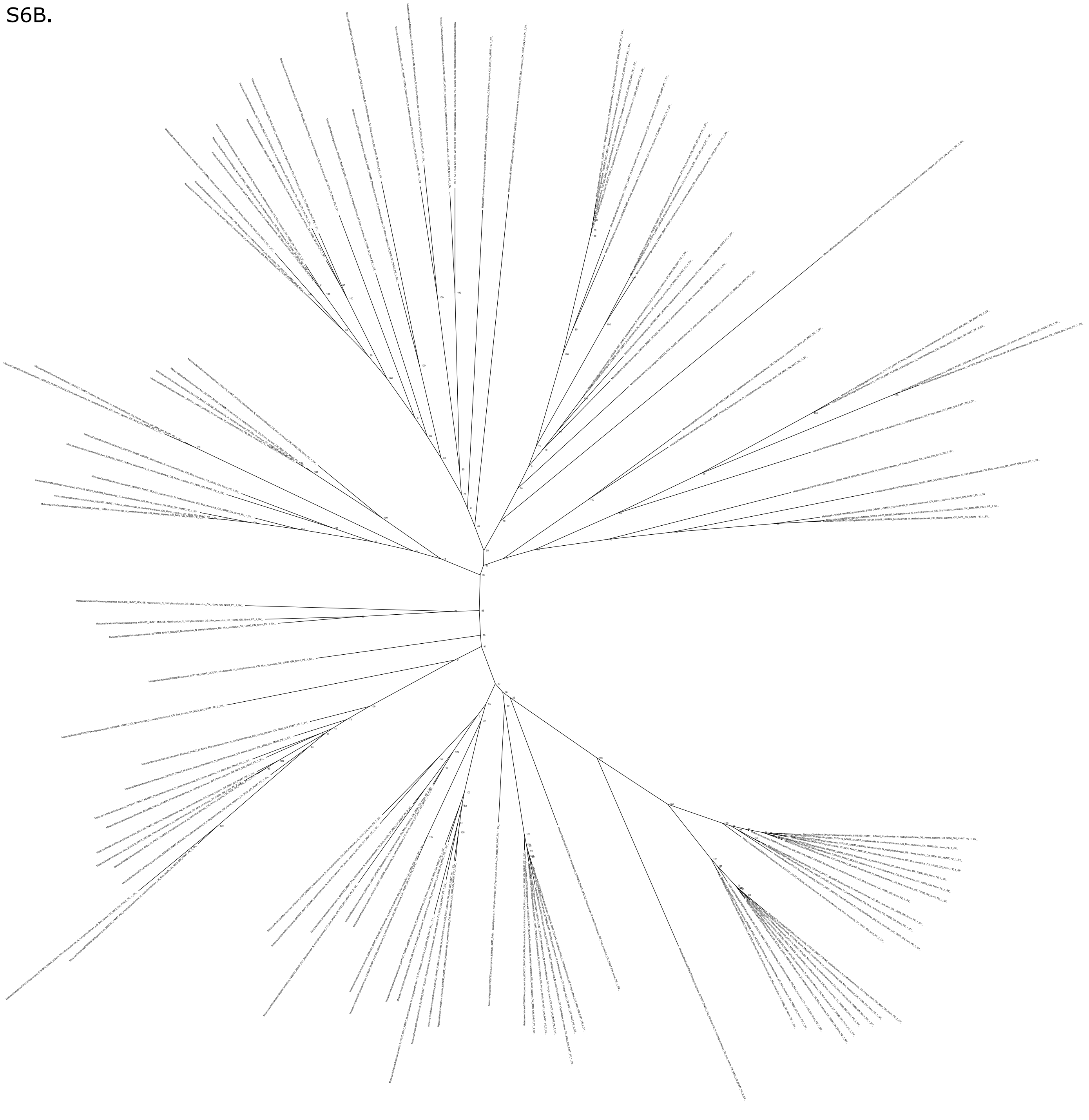

S6C.

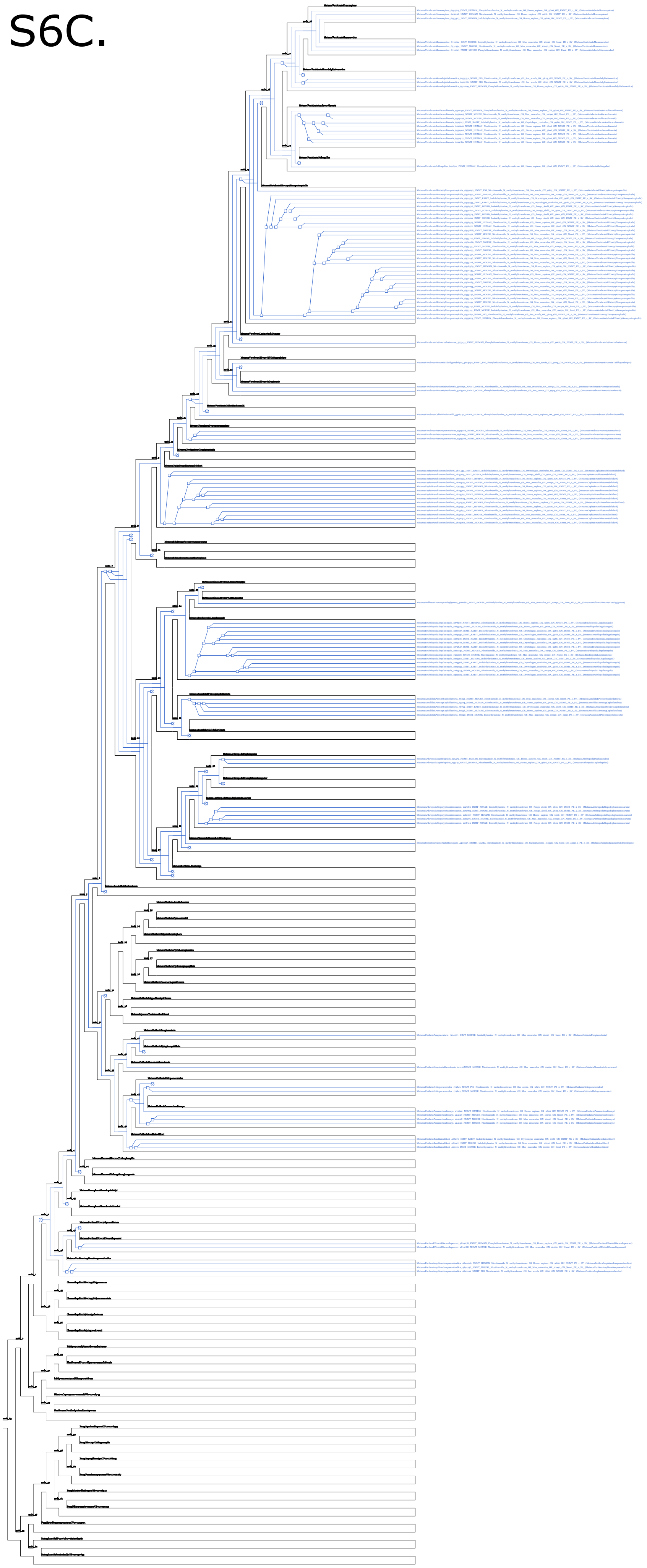

S6D

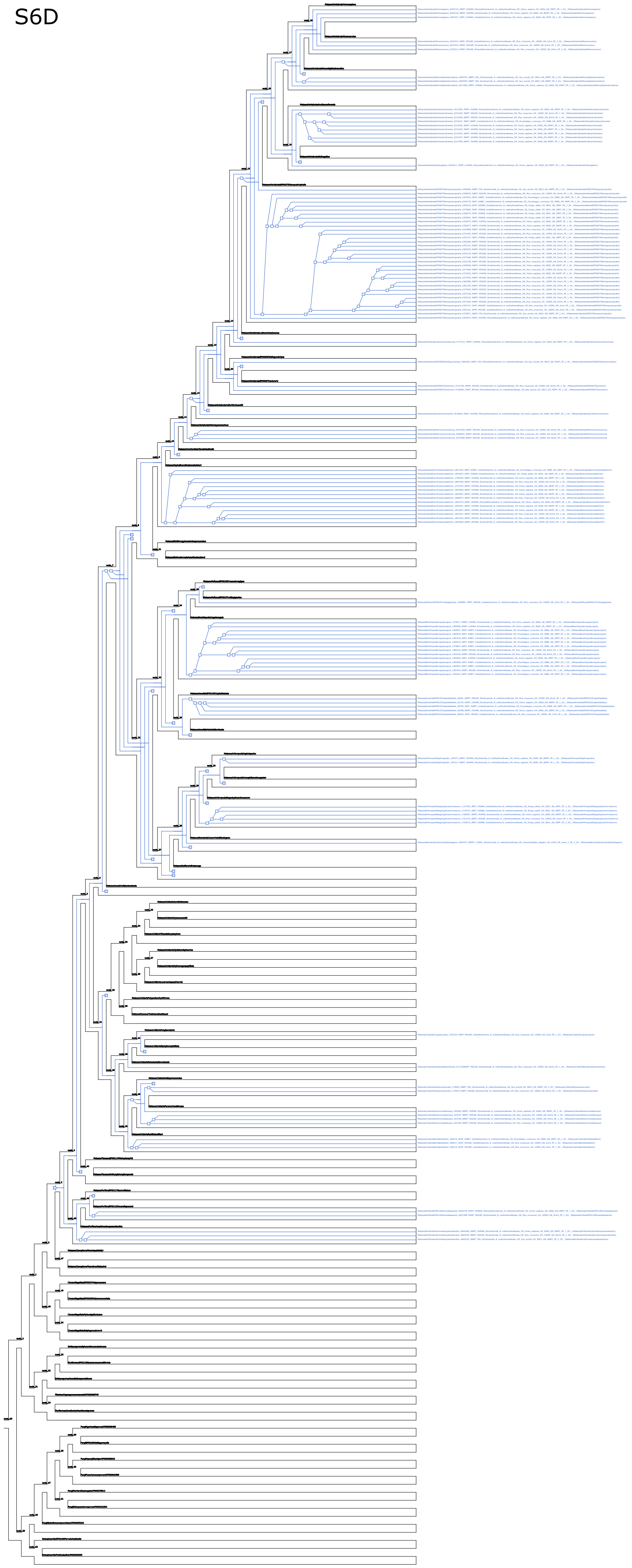

S7A.

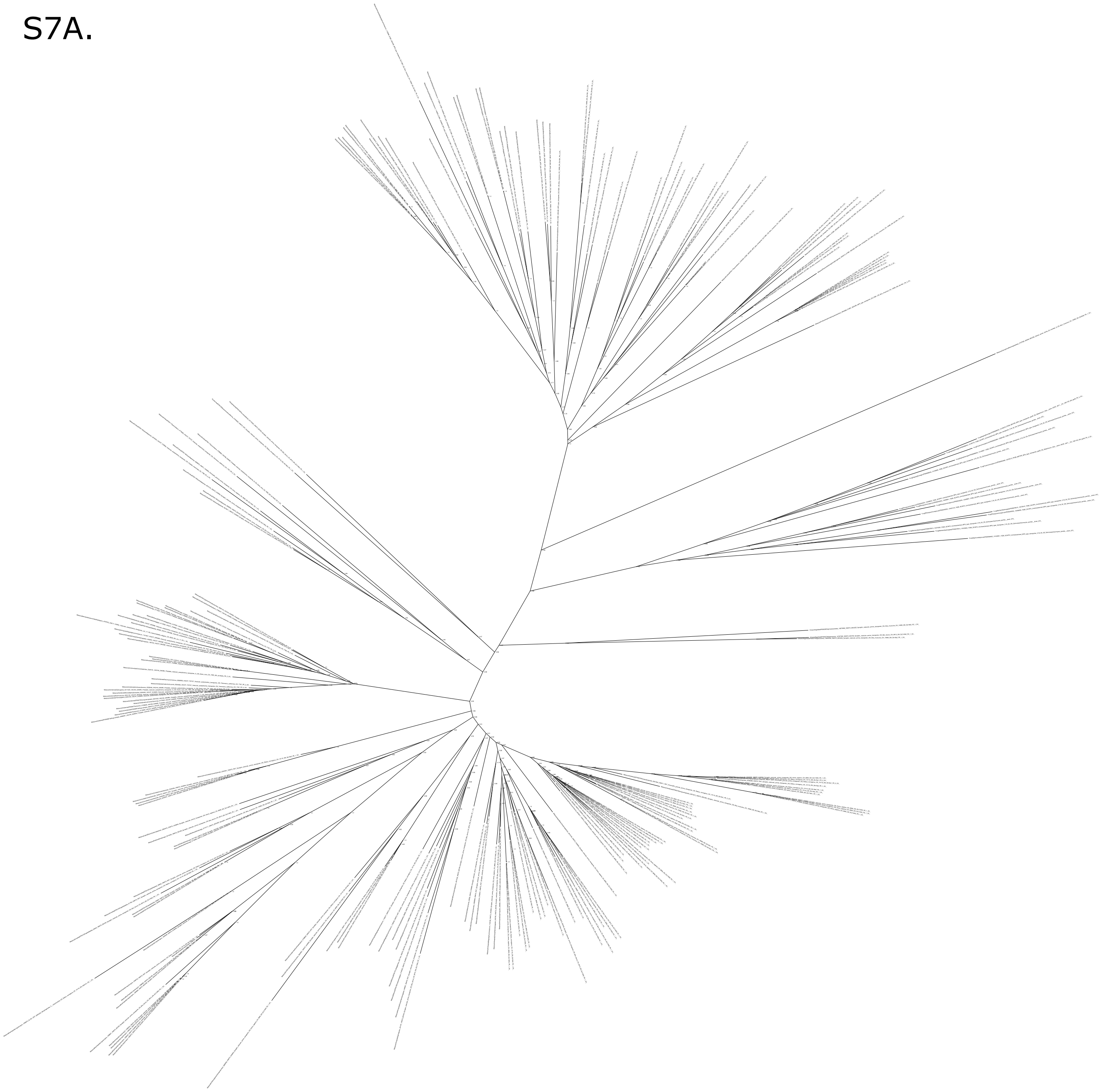

S7B.

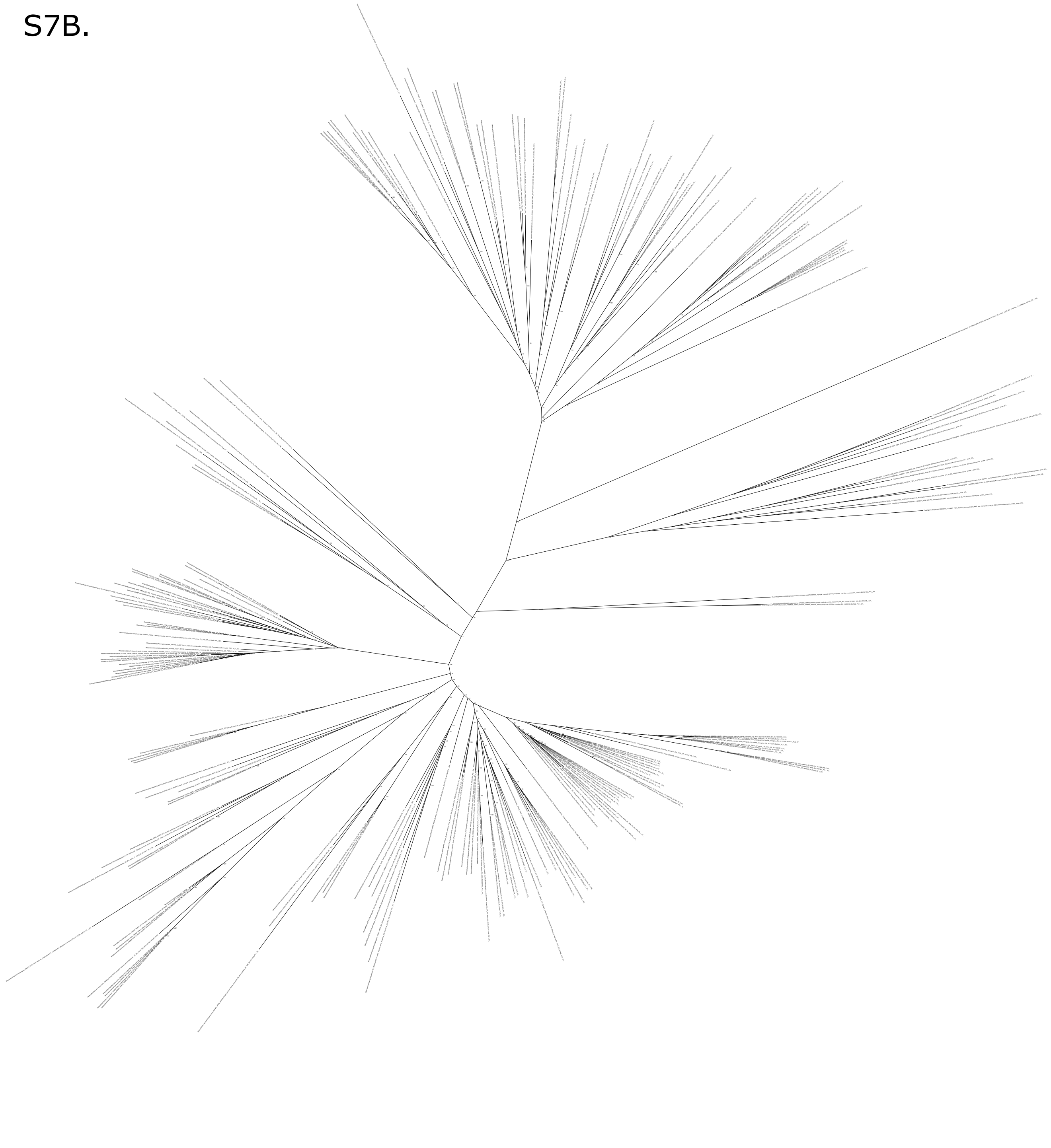

S7C.

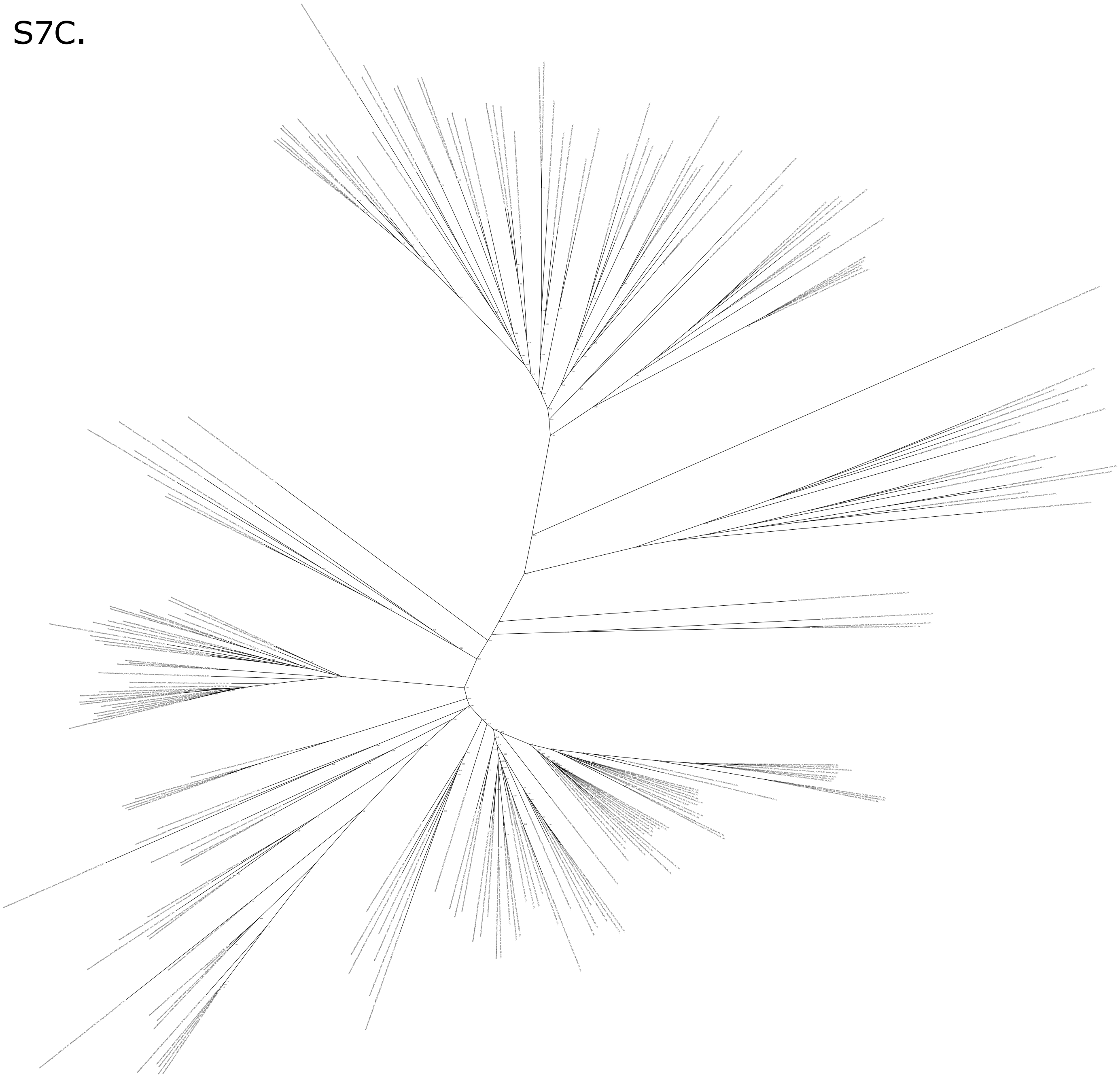

S7D.

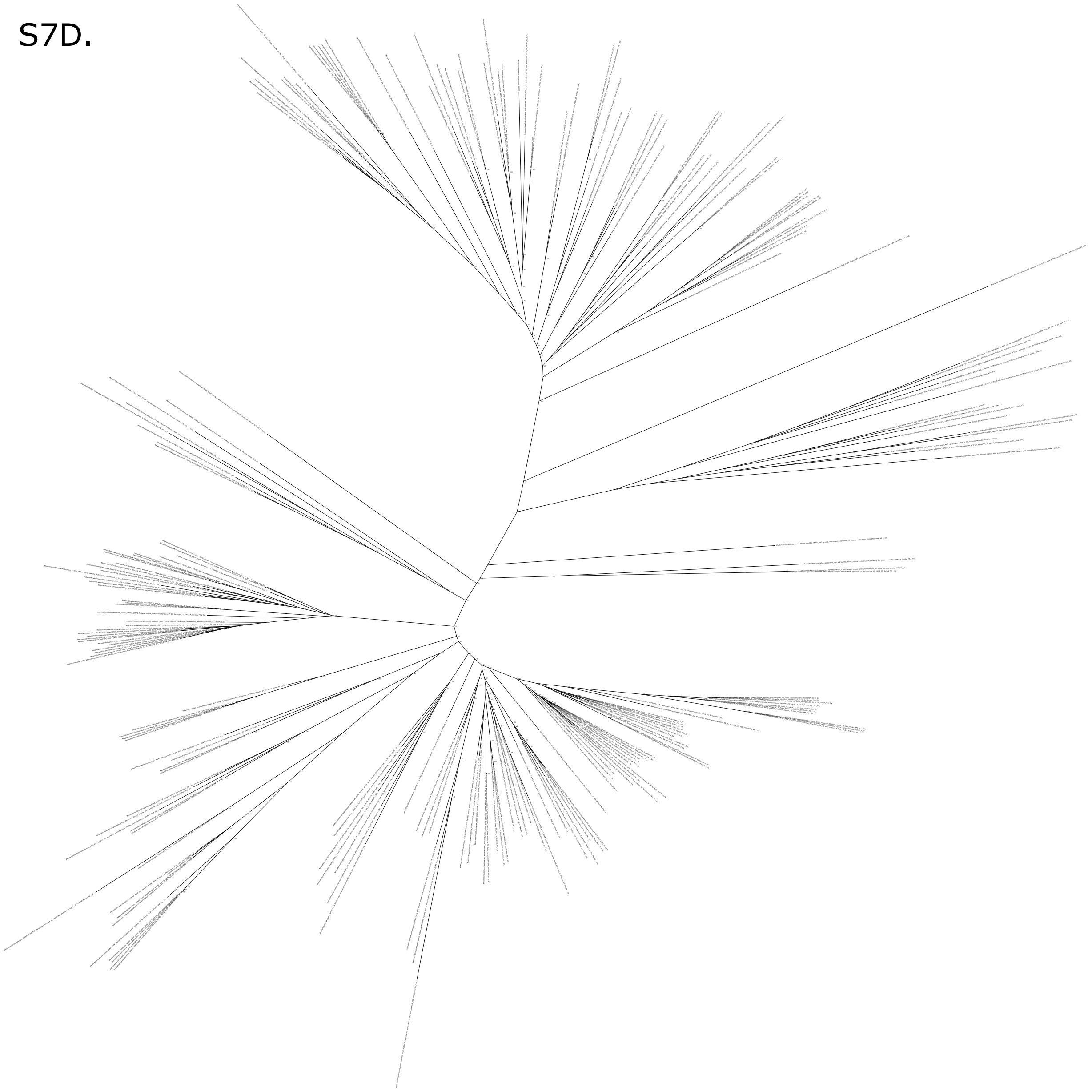

S7E.

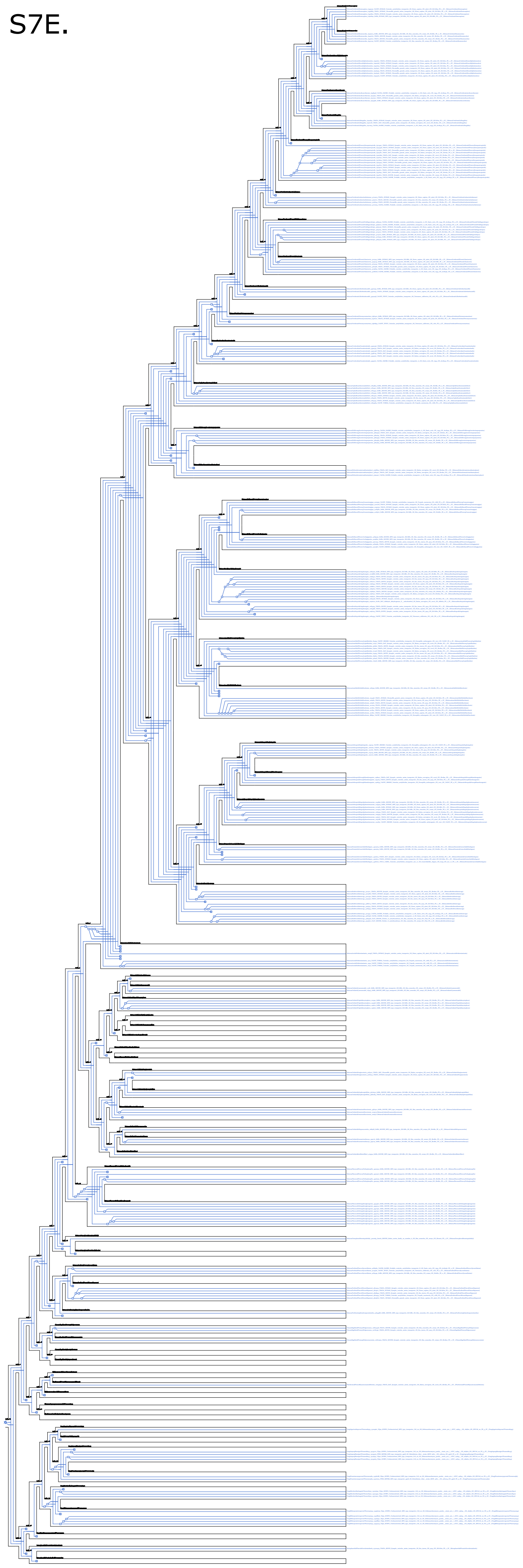

S7F

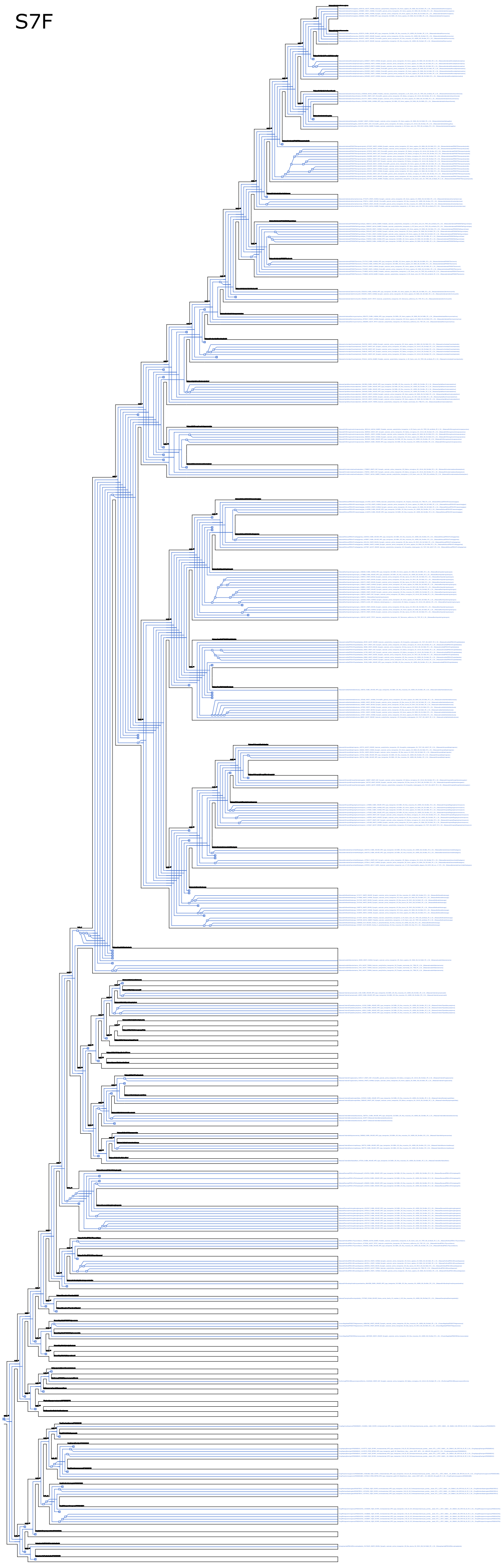

S8A.

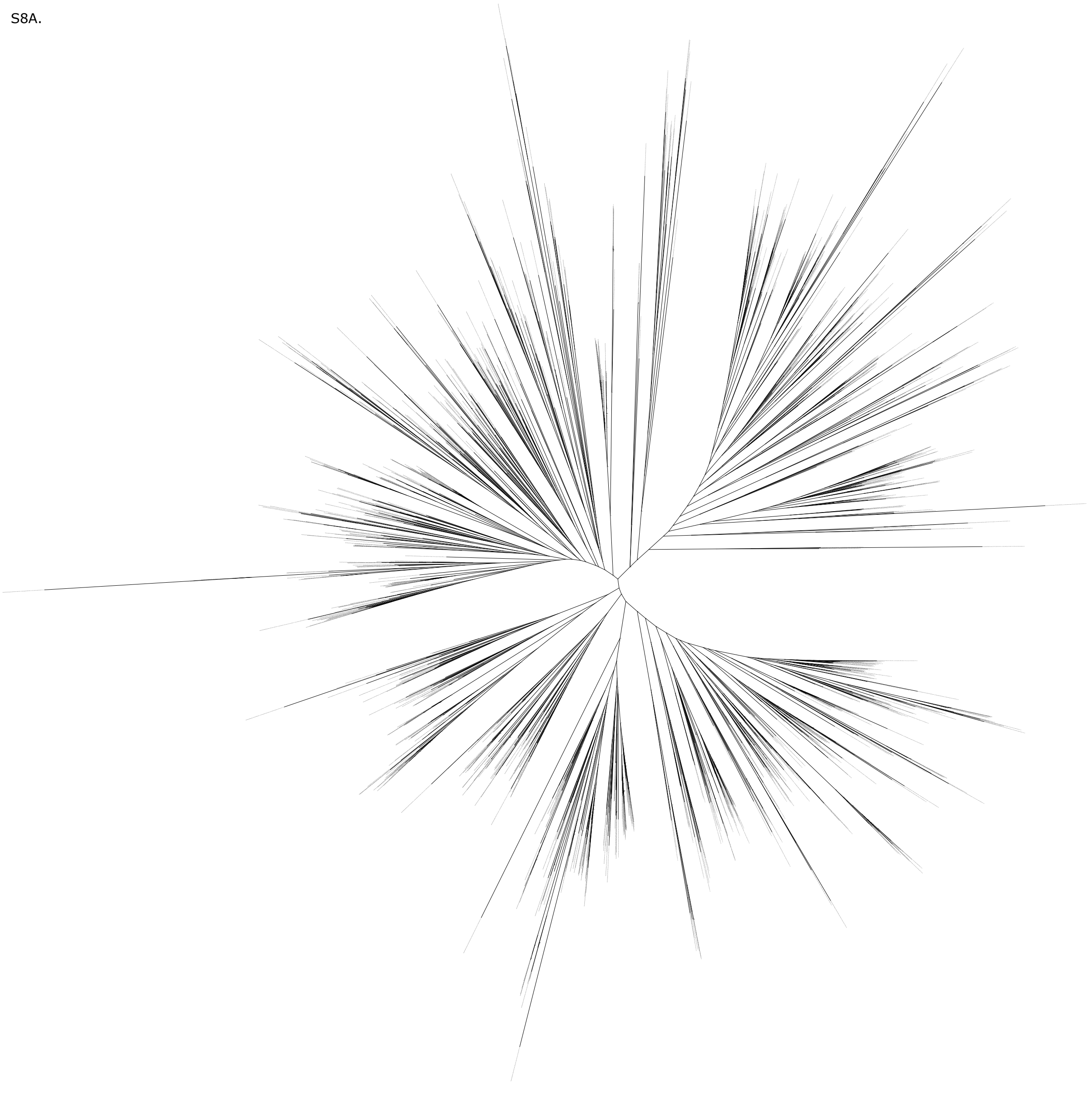

S8B.

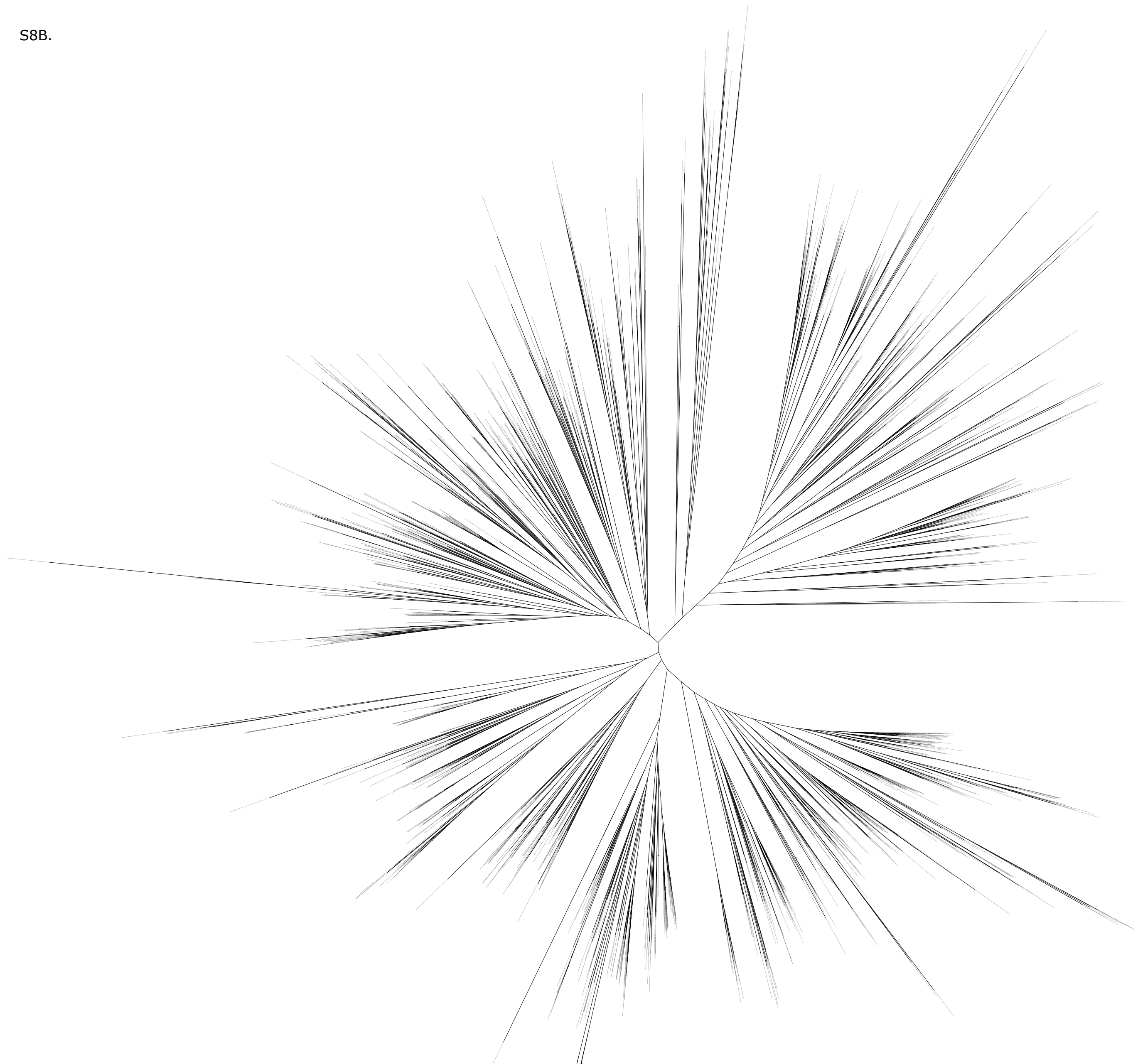

S8C

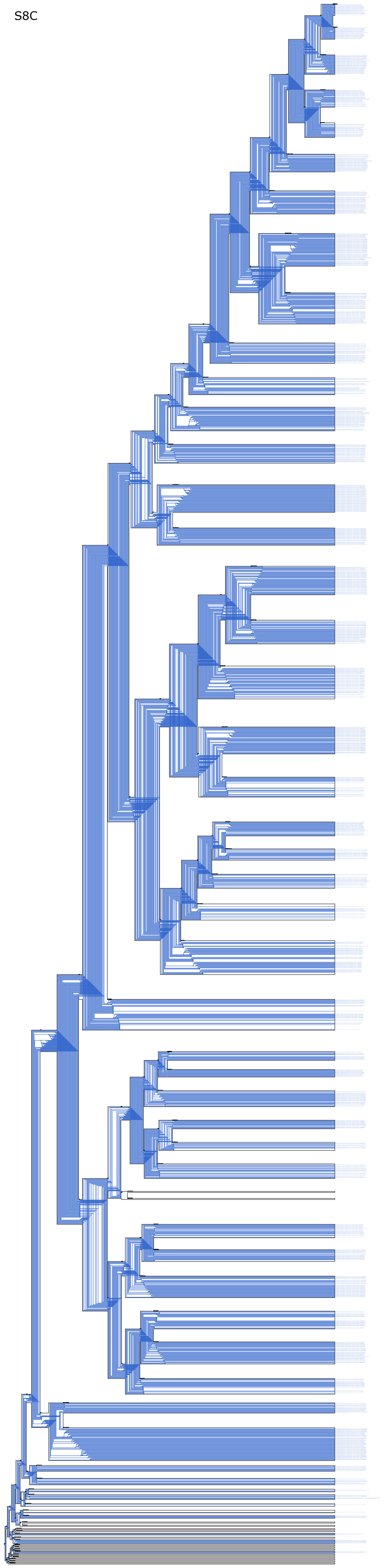

S8D

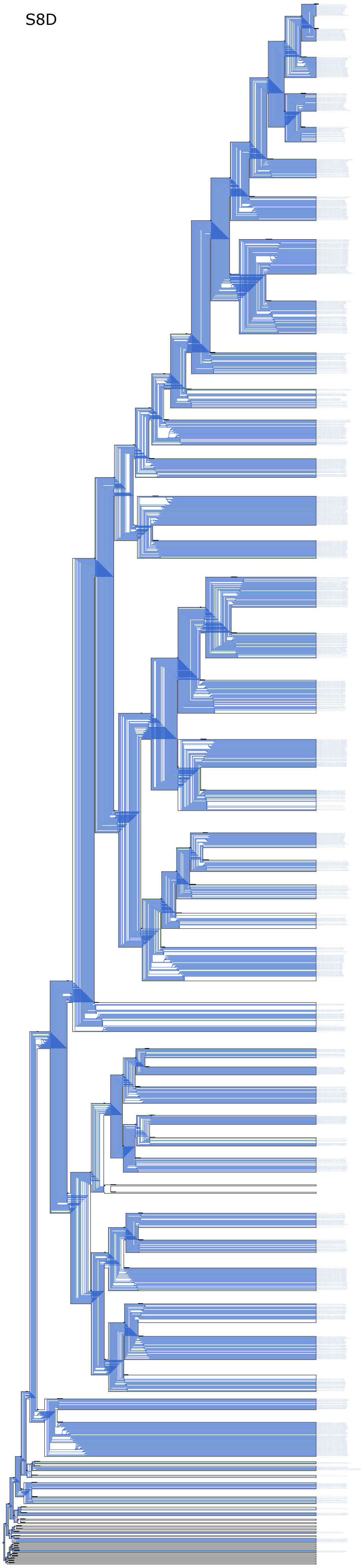

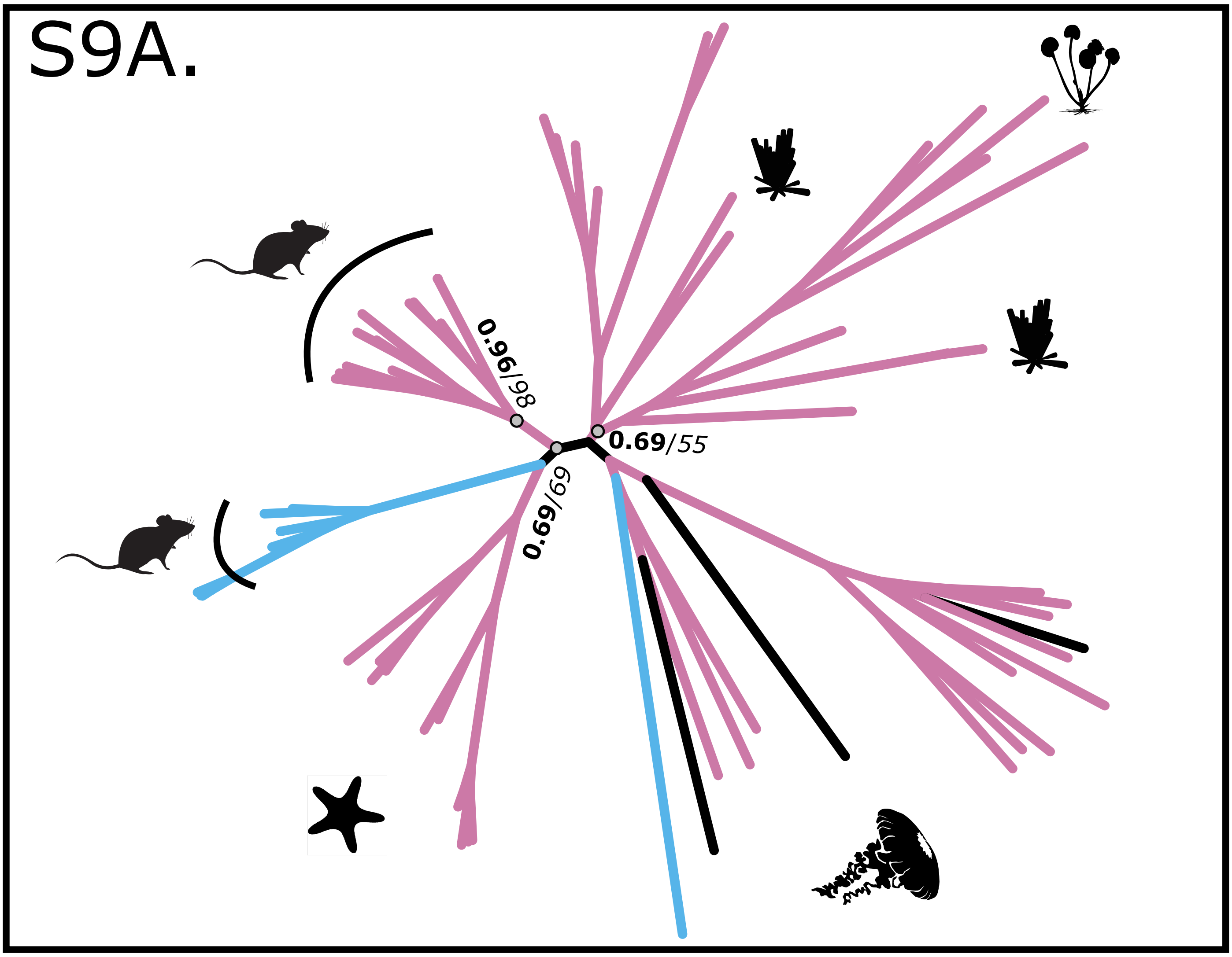
S9A.
0.96/98
0.69/55
0.69/69

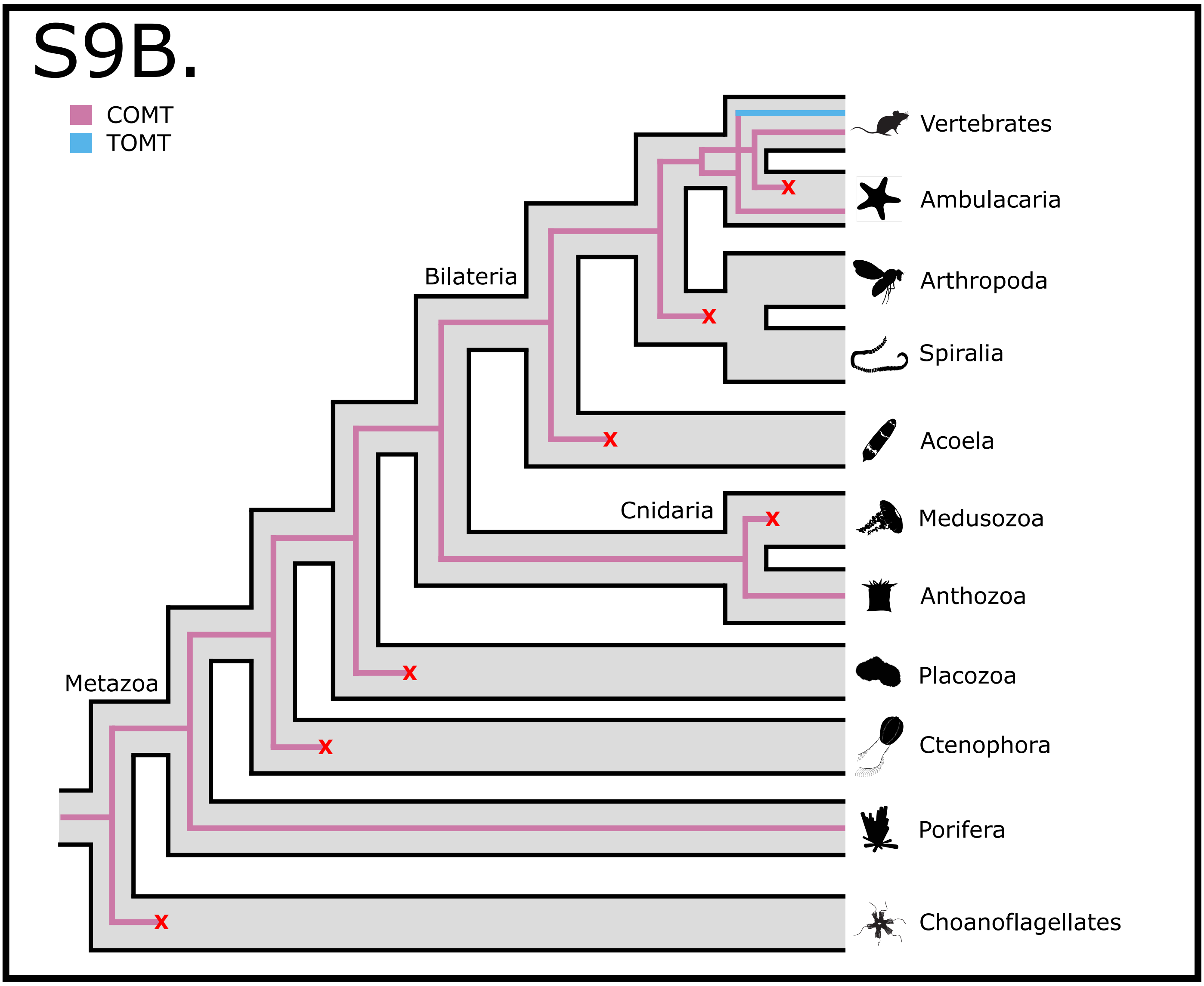
S9B.
COMT
TOMT
Vertebrates
Ambulacaria
Bilateria
Arthropoda
Spiralia
Acoela
Cnidaria
Medusozoa
Anthozoa
Placozoa
Ctenophora
Metazoa
Porifera
Choanoflagellates

S10A.

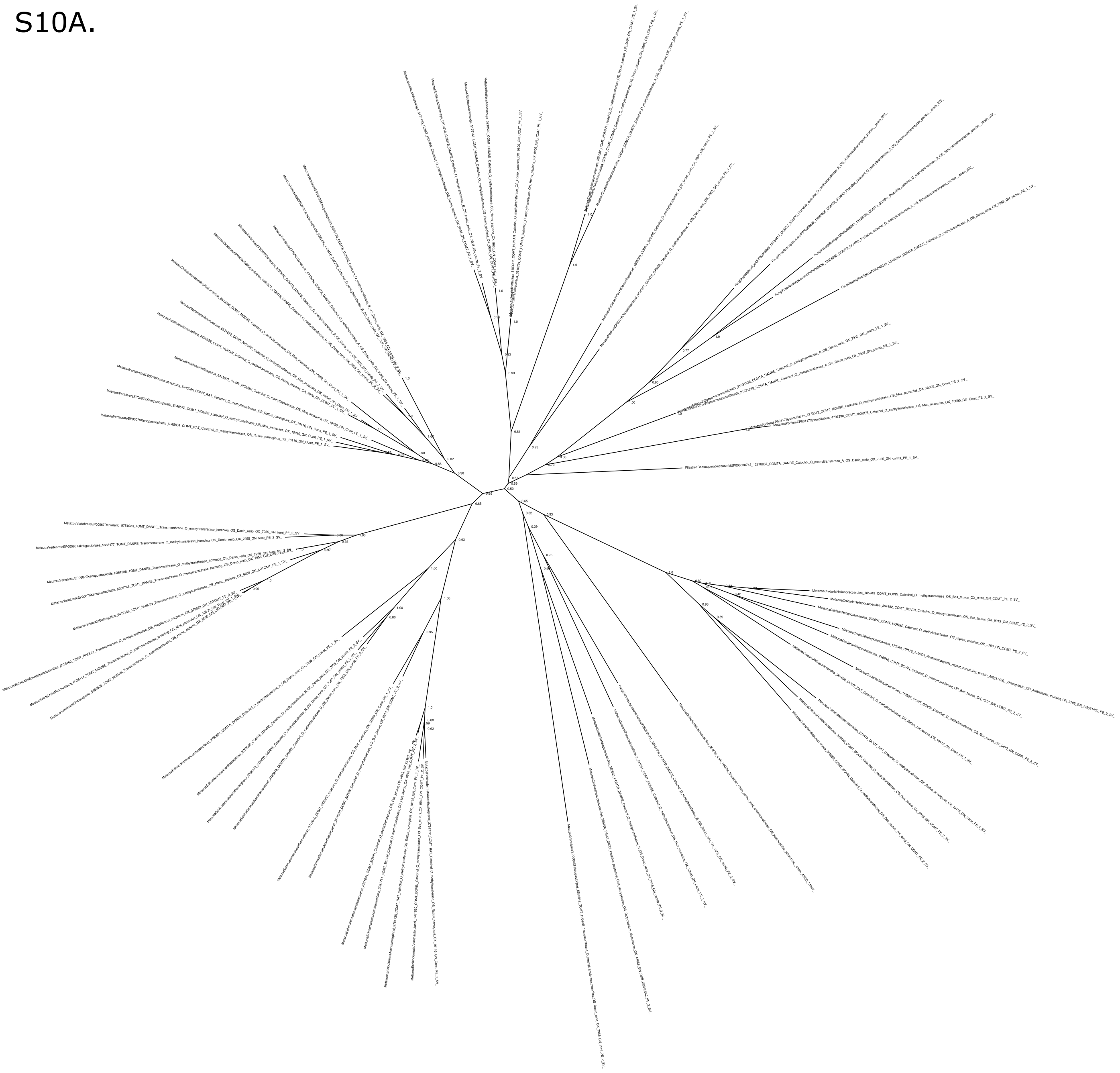

S10B.

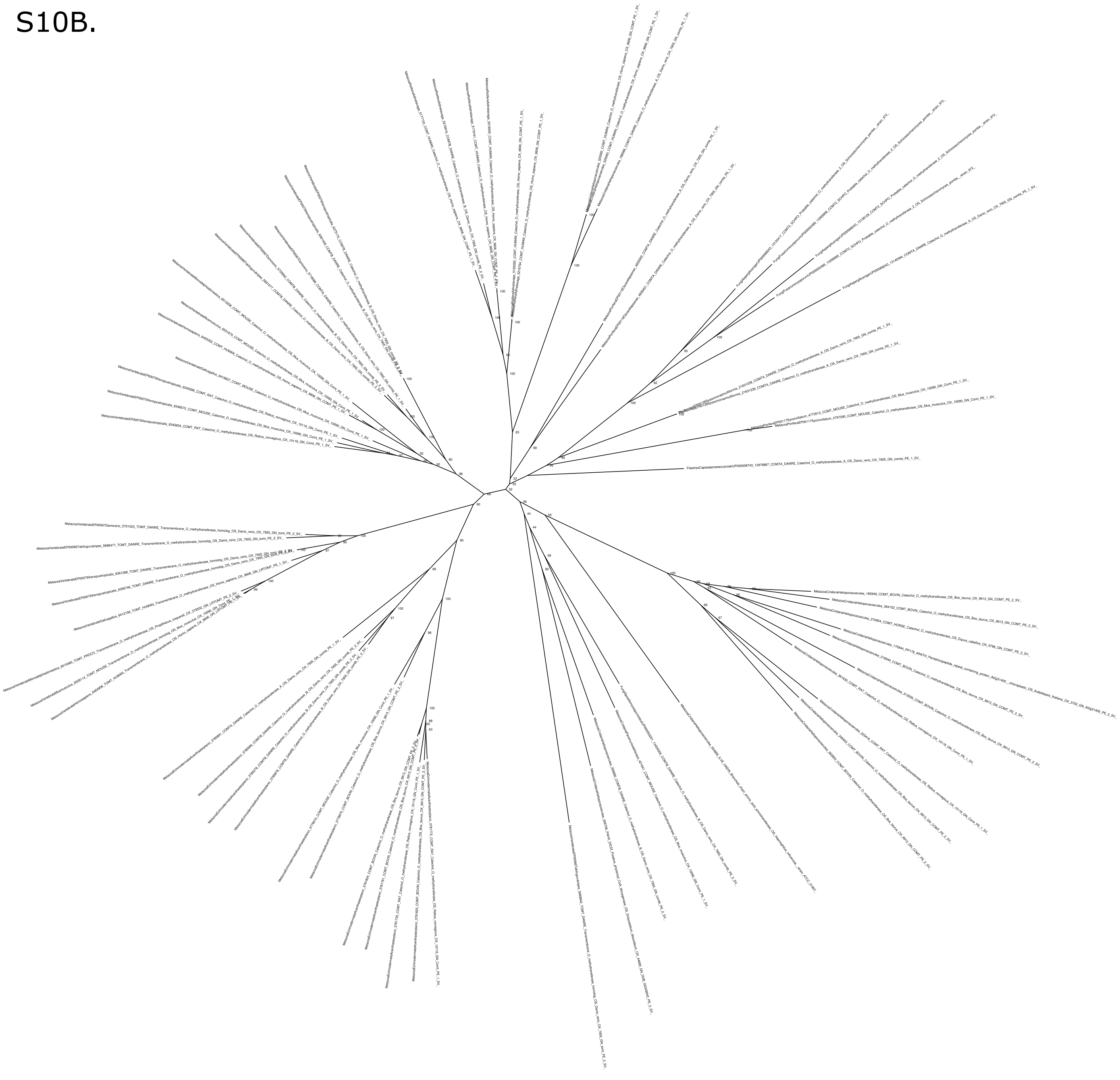

S10C.

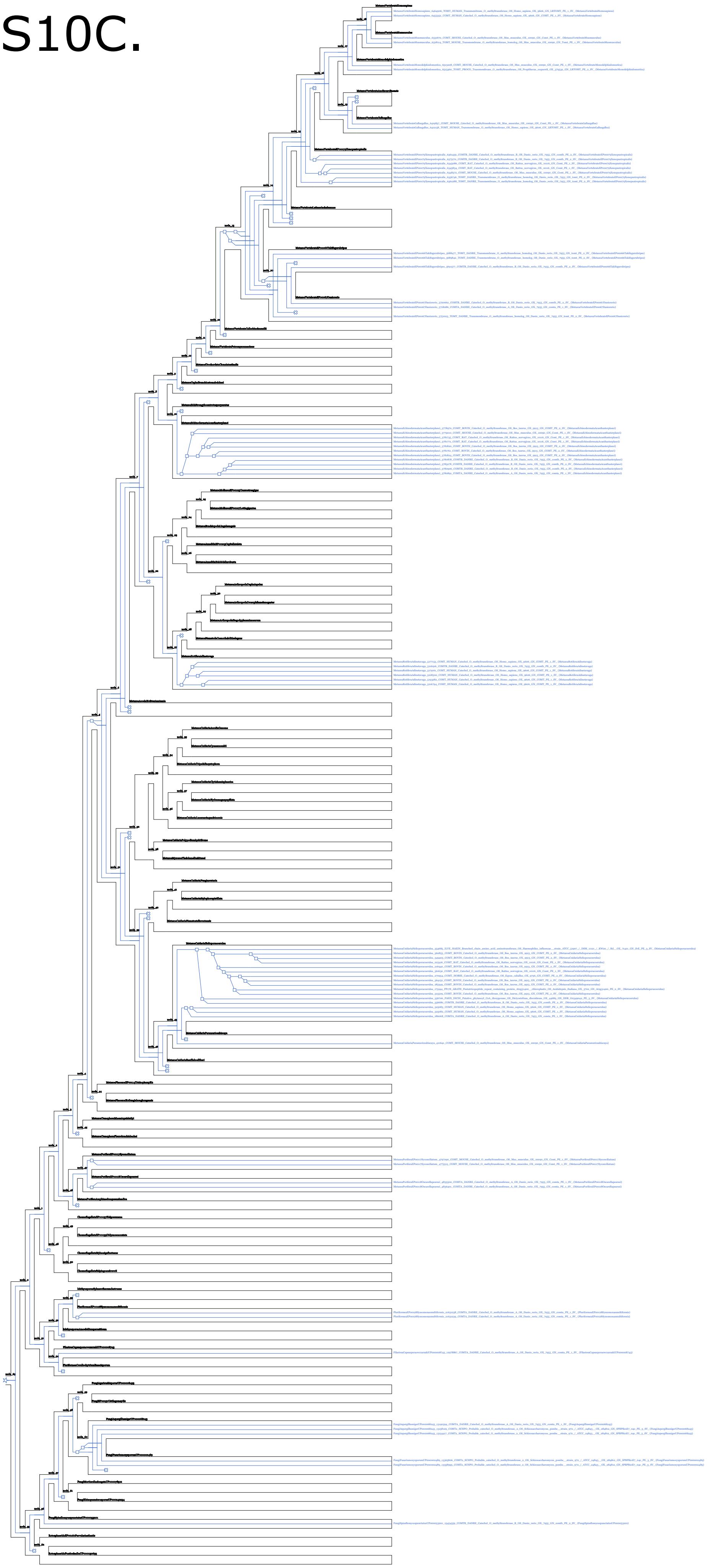

S10D

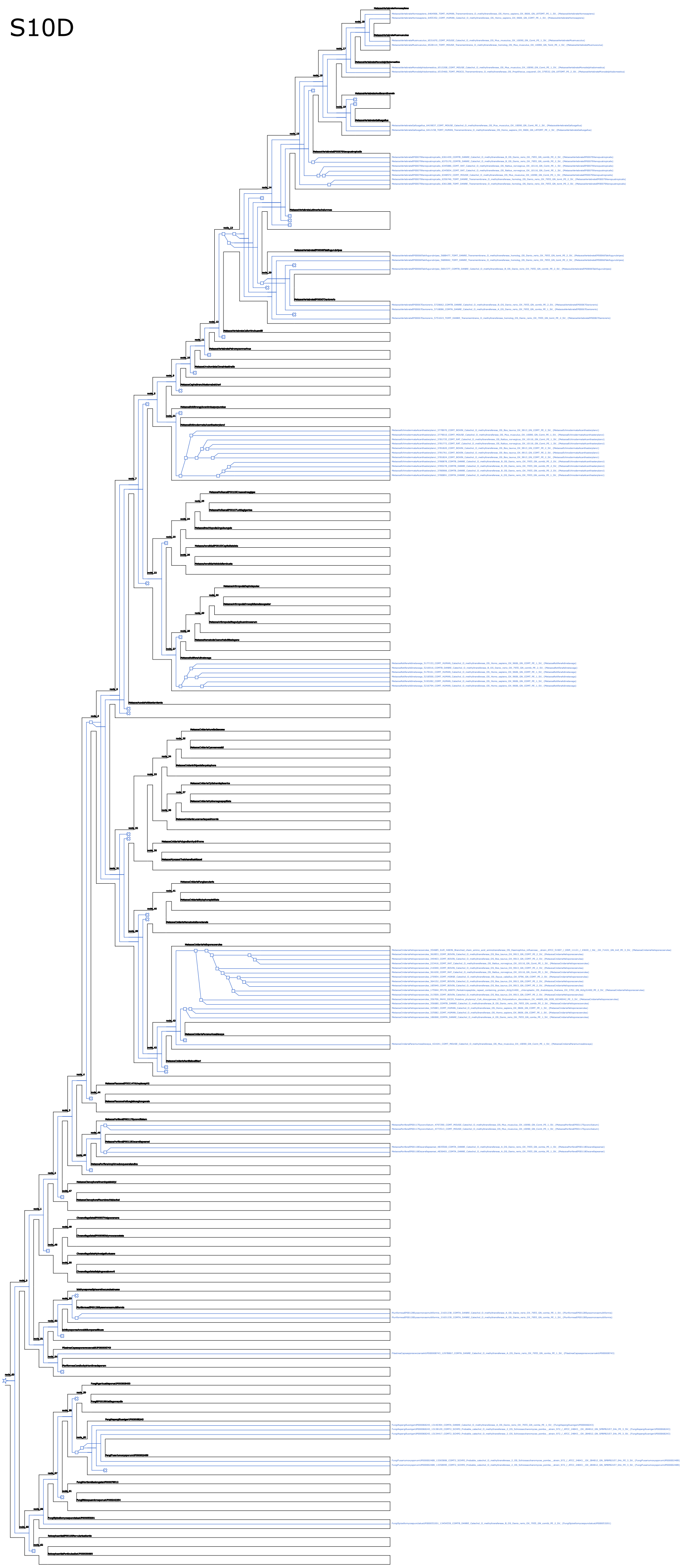

S11A.

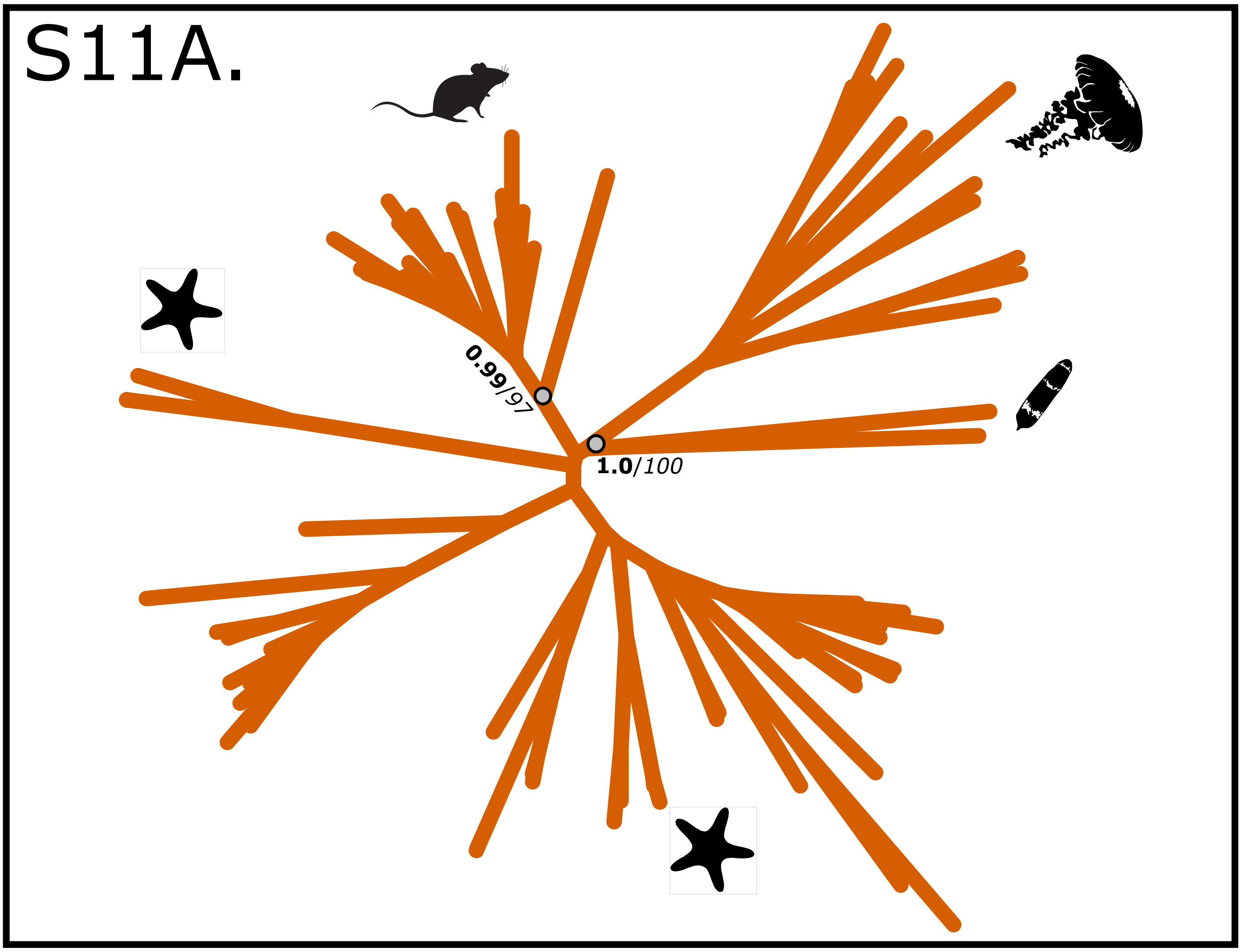

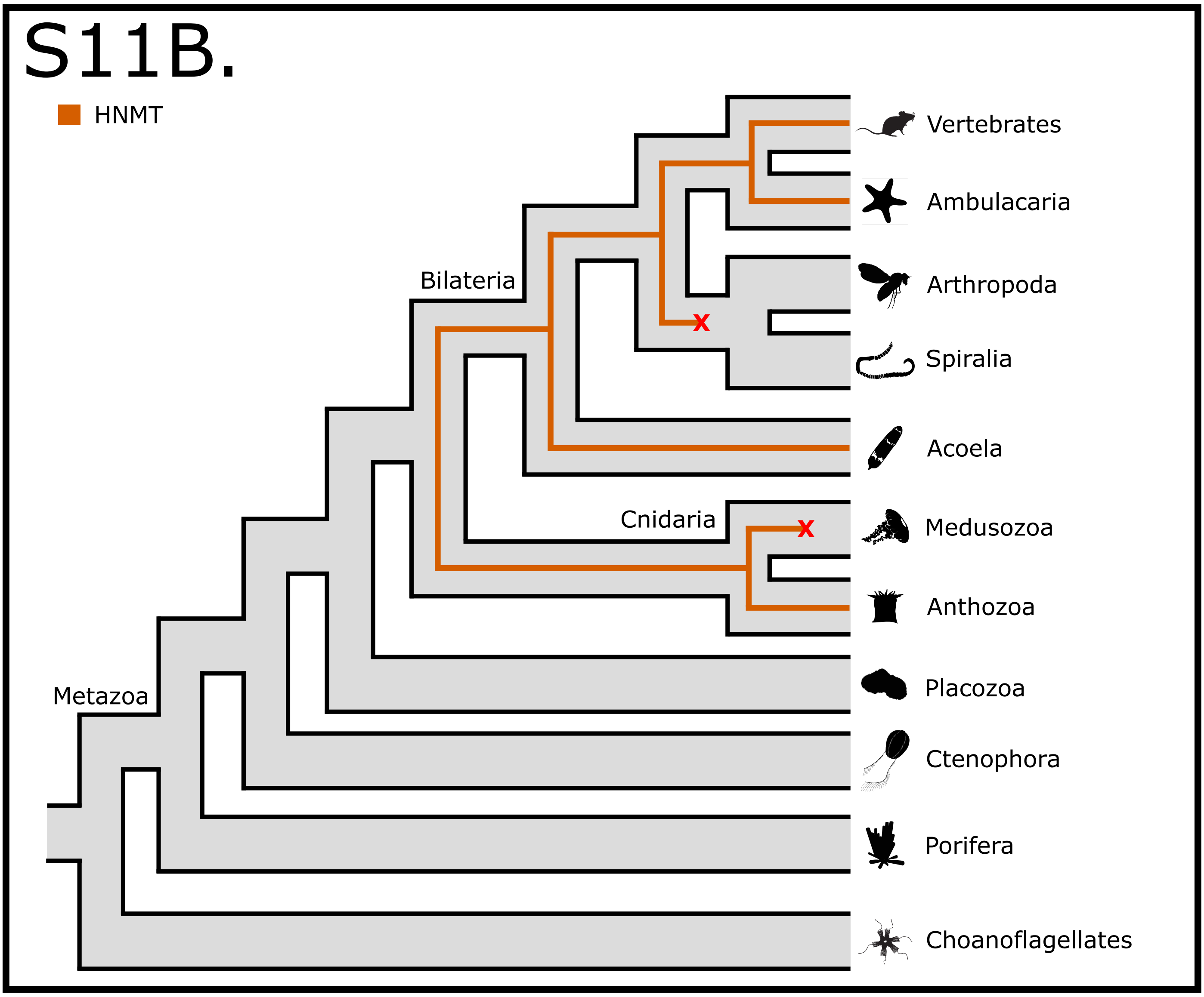
S11B.
HNMT
Bilateria
Cnidaria
Metazoa
Vertebrates
Ambulacaria
Arthropoda
Spiralia
Acoela
Medusozoa
Anthozoa
Placozoa
Ctenophora
Porifera
Choanoflagellates

S12A.

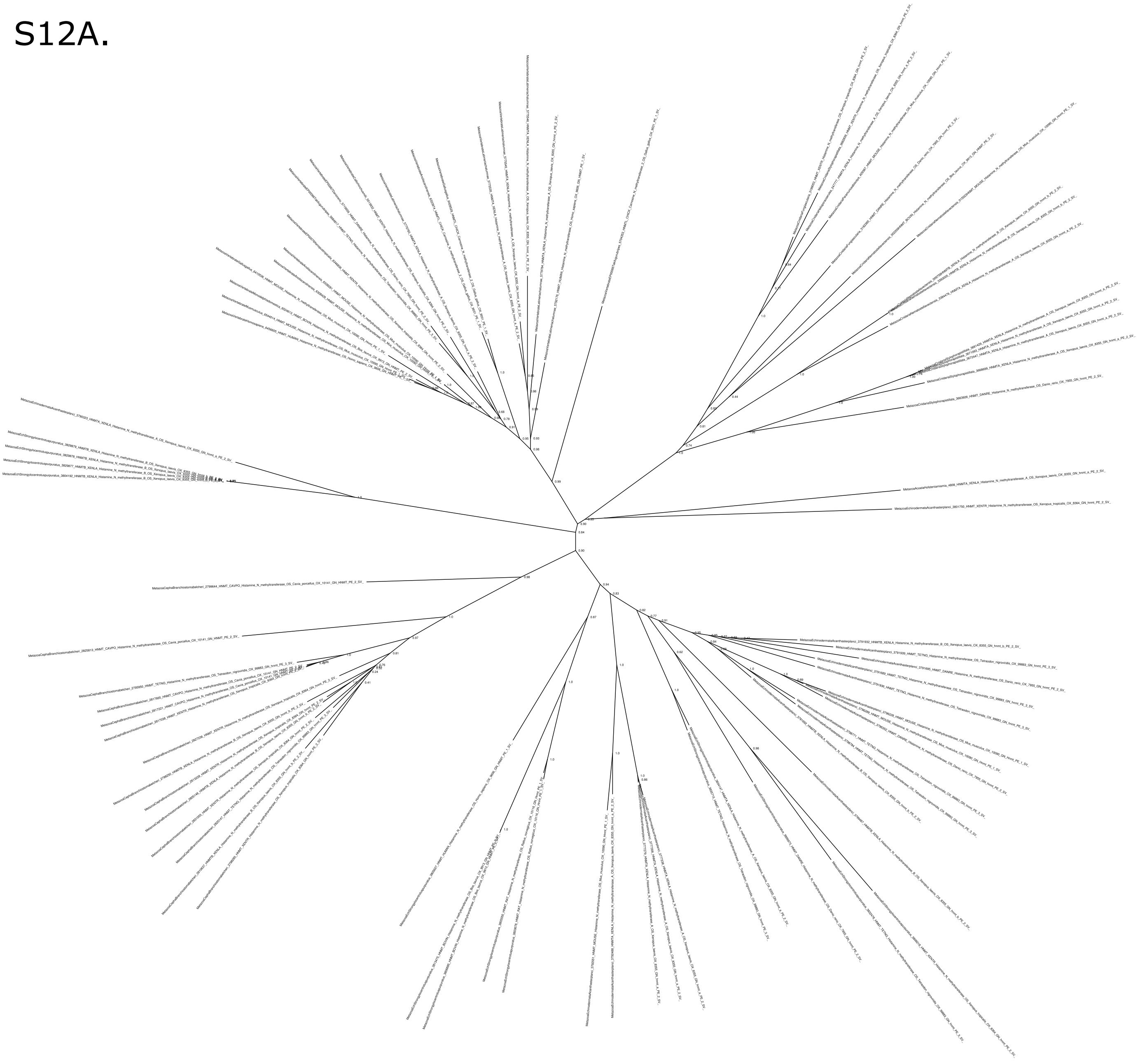

S12B.

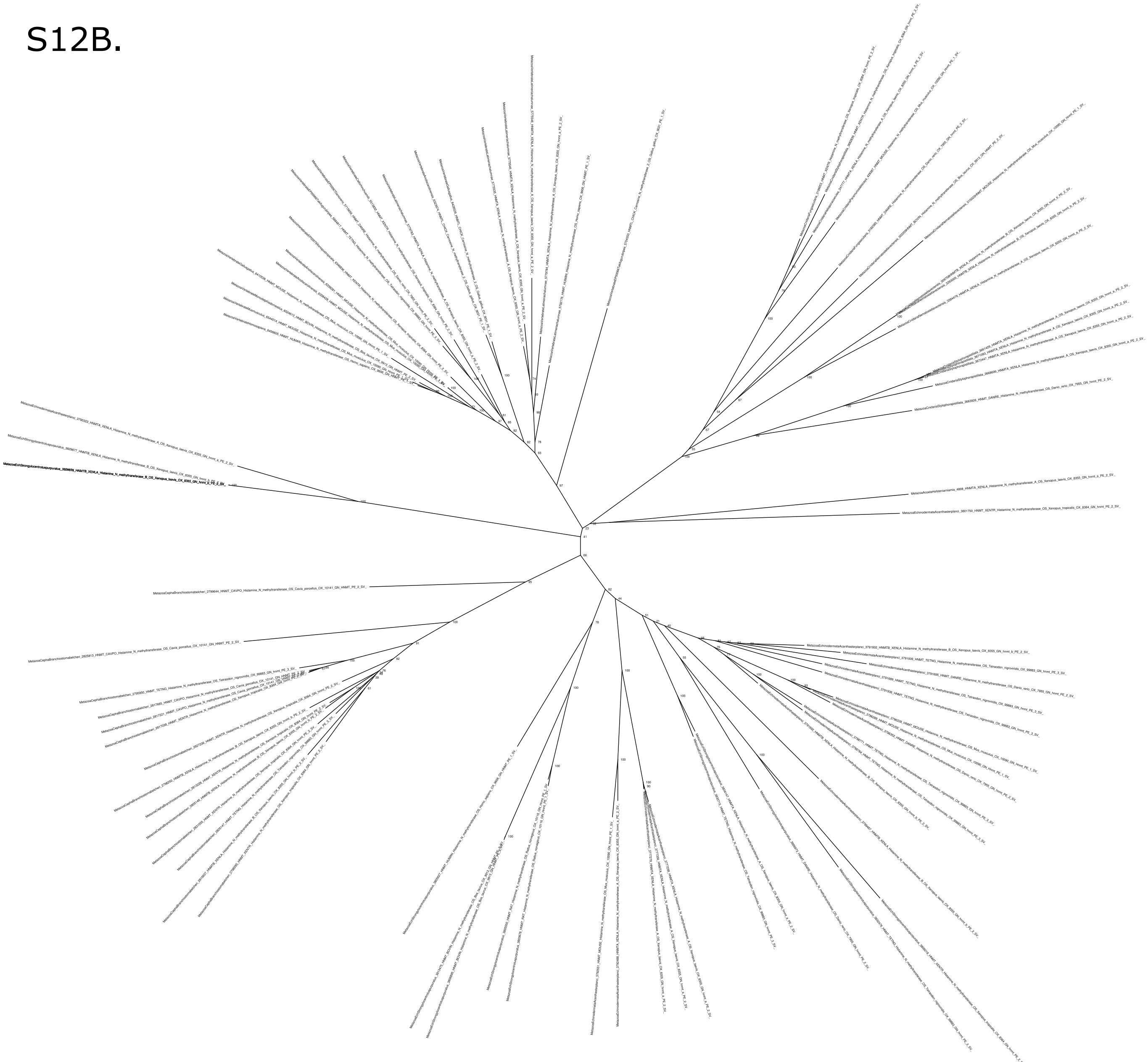

S12C.

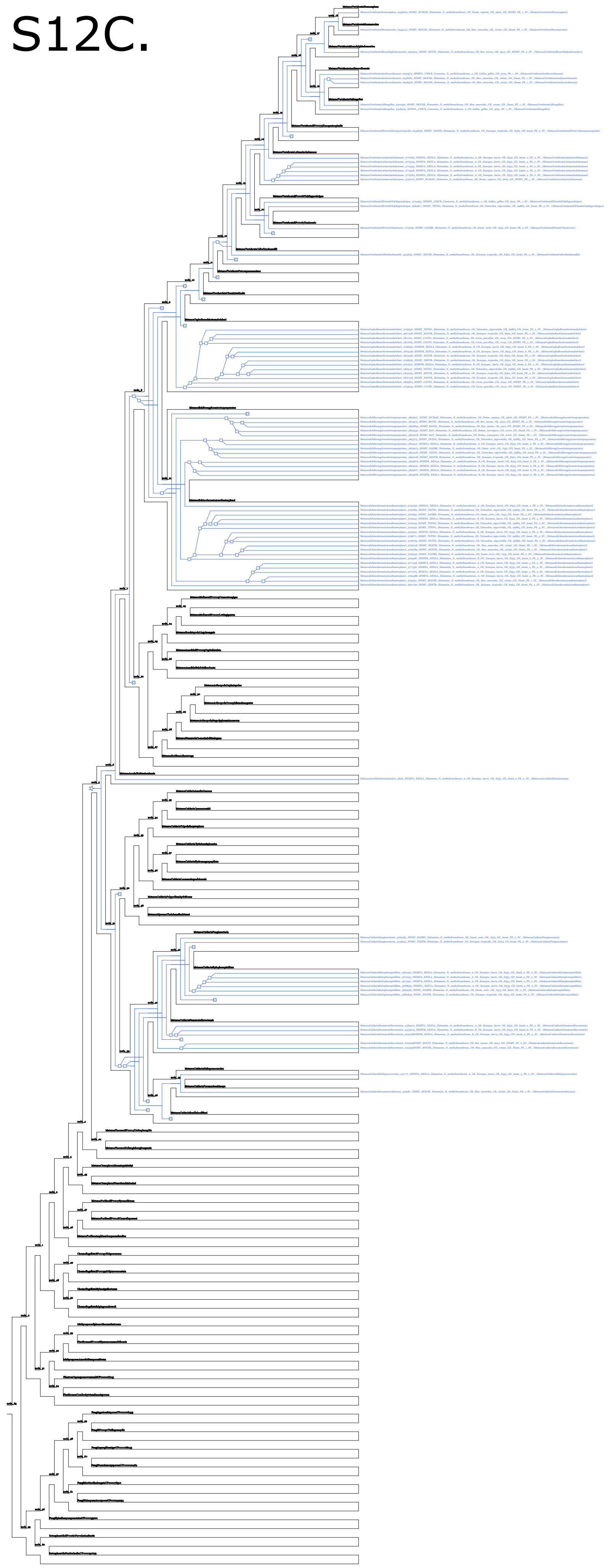

S12D

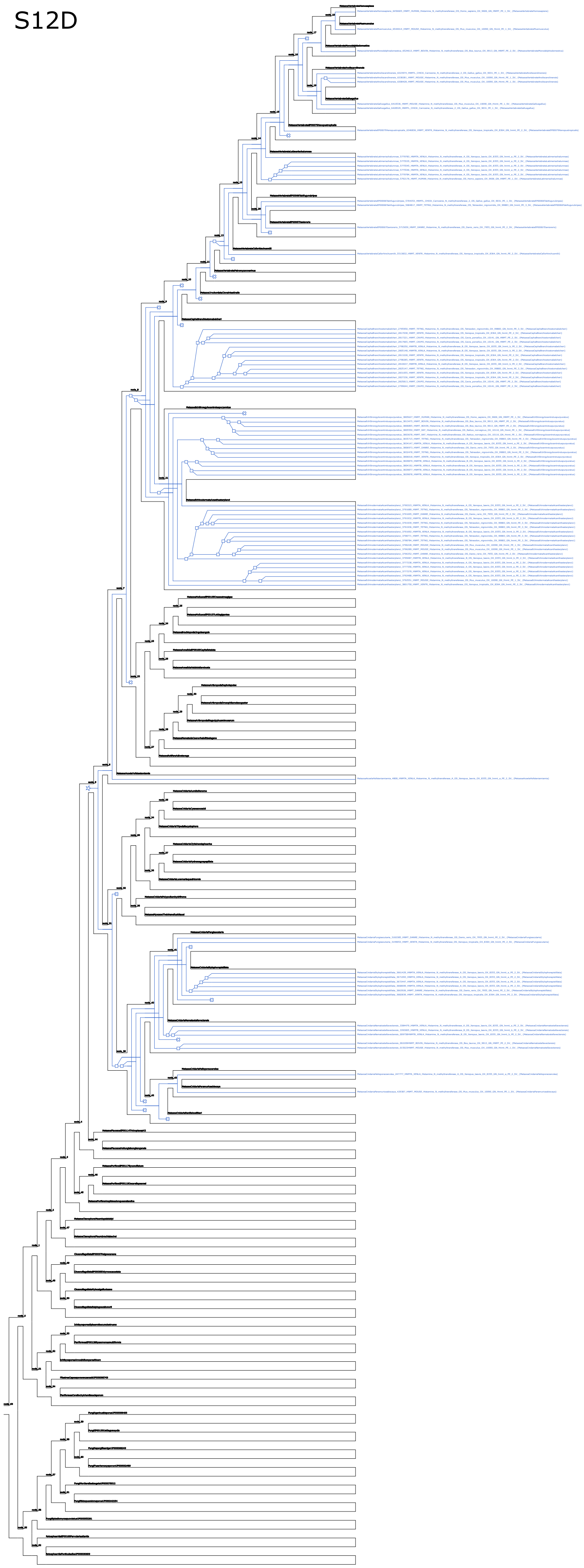

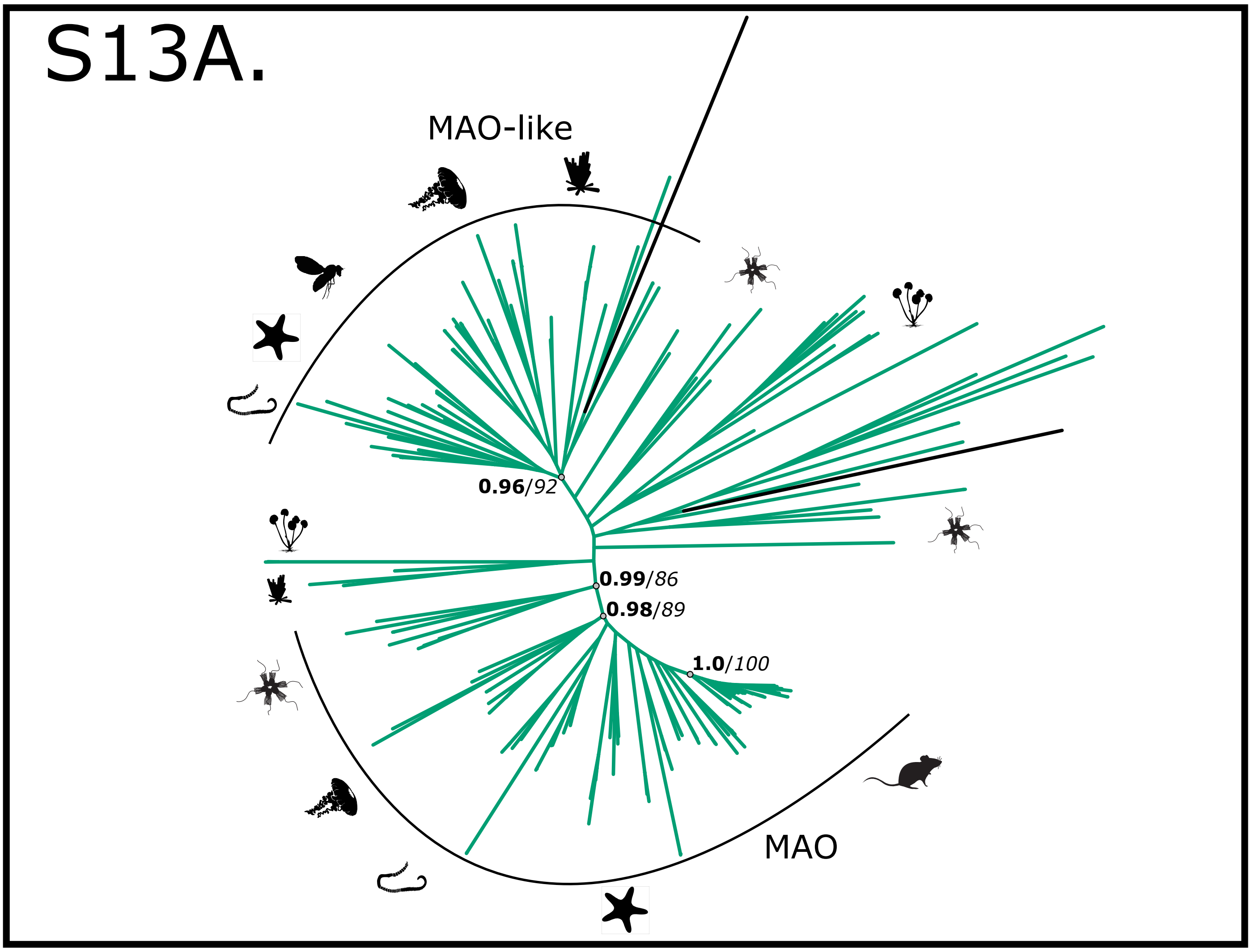

S13A.
MAO-like
0.96/92
0.99/86
0.98/89
1.0/100
MAO

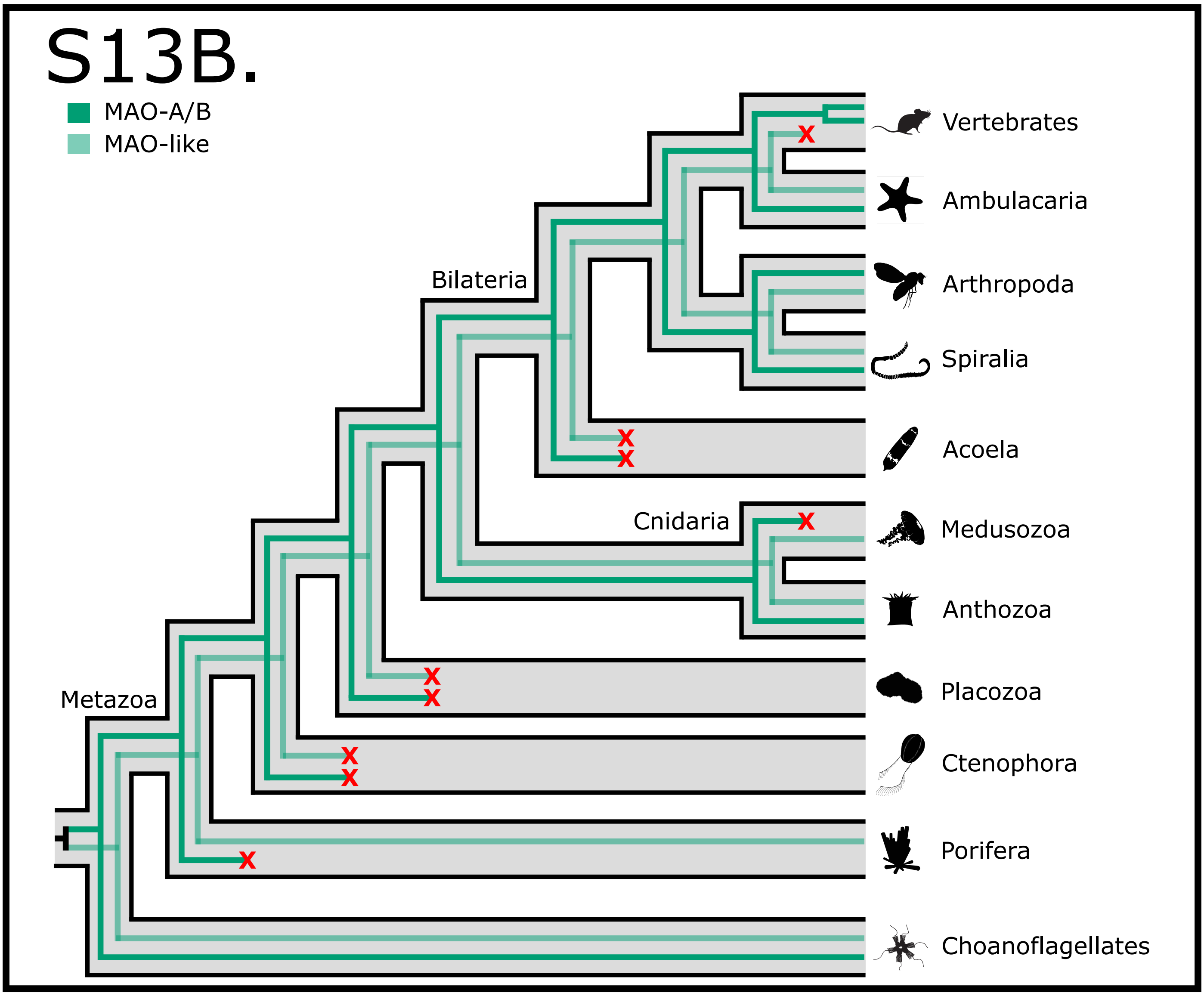
S13B.
MAO-A/B
MAO-like
Bilateria
Cnidaria
Metazoa
Vertebrates
Ambulacaria
Arthropoda
Spiralia
Acoela
Medusozoa
Anthozoa
Placozoa
Ctenophora
Porifera
Choanoflagellates

S14A.

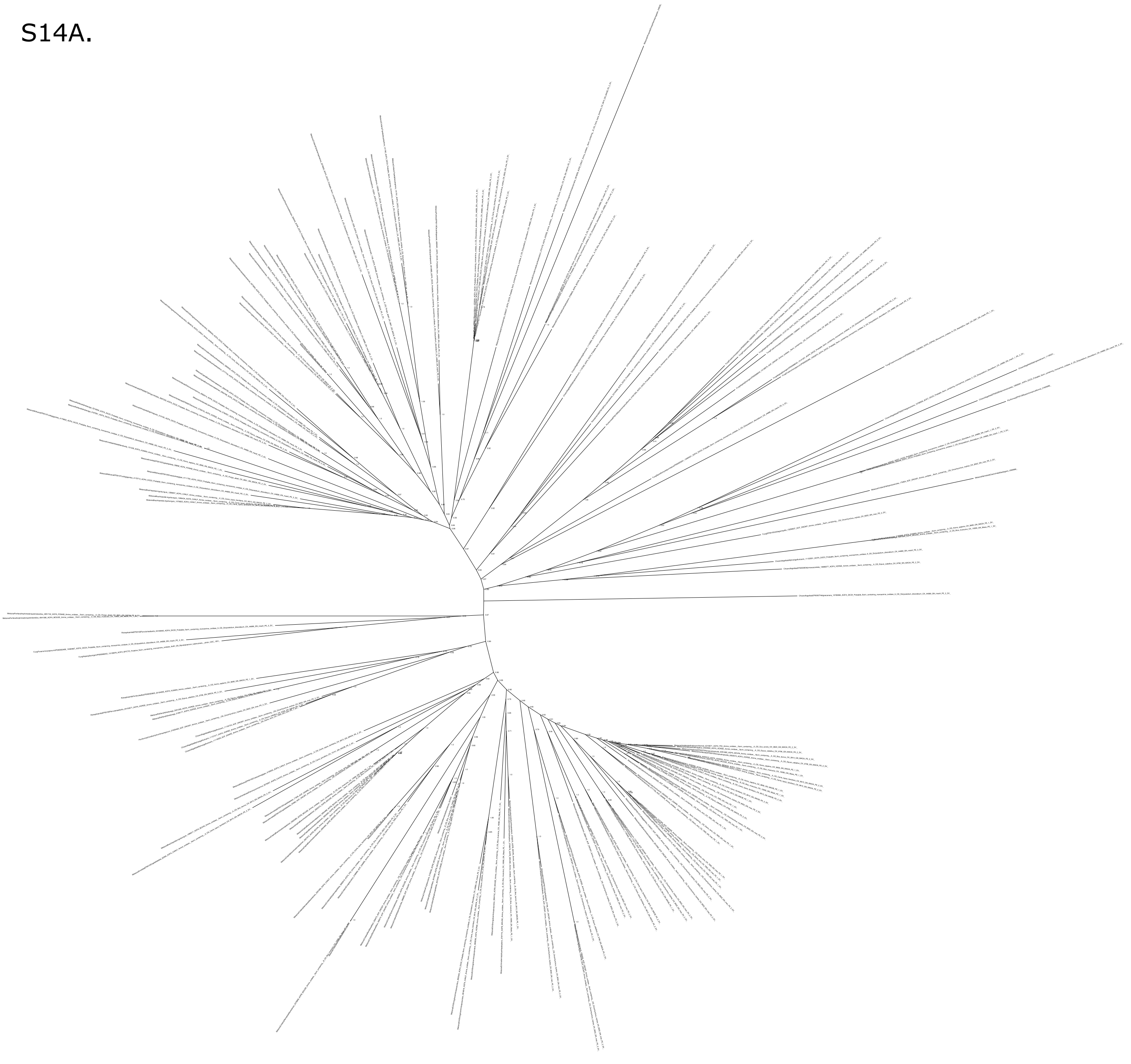

S14B.

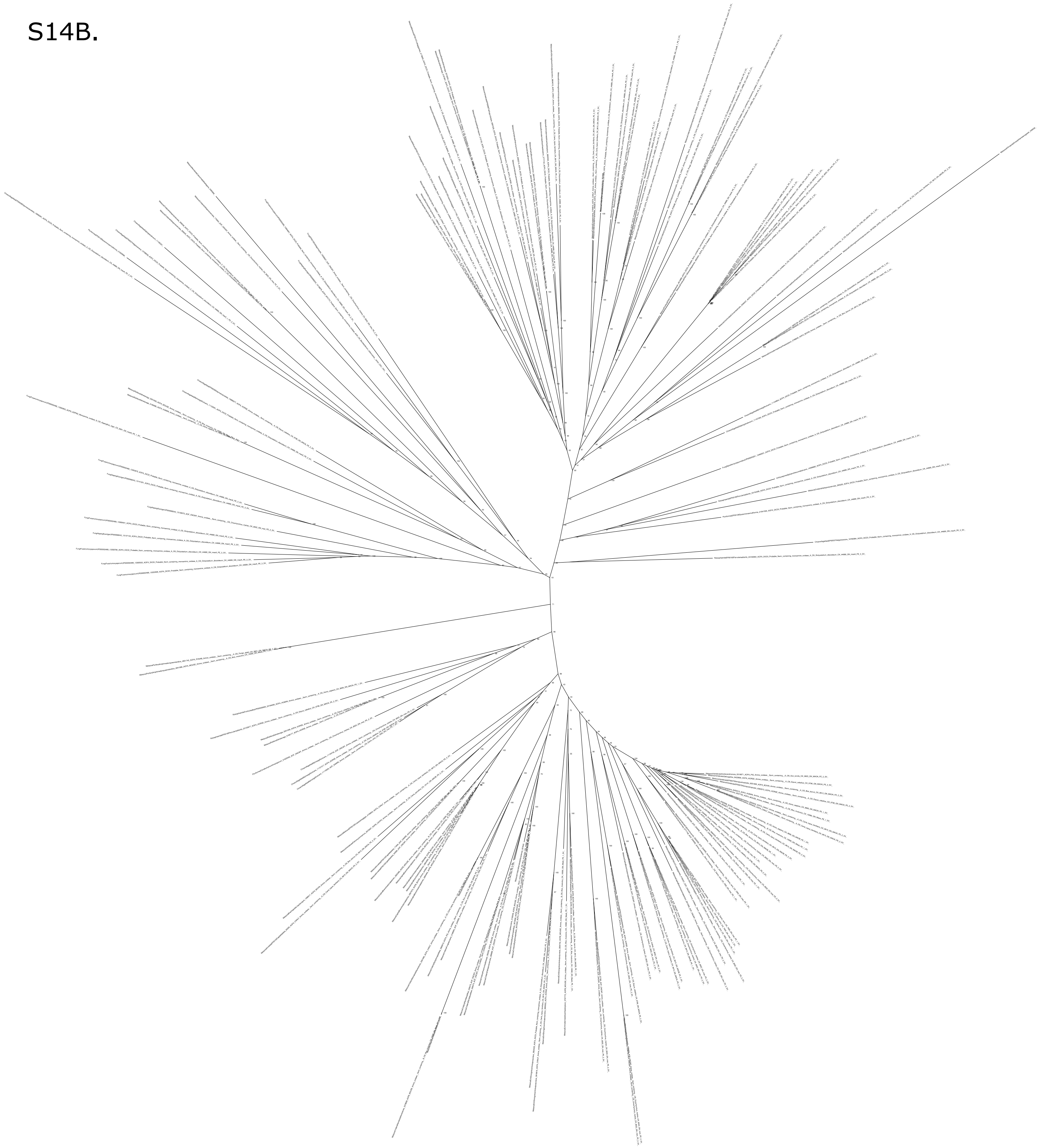

S14C.

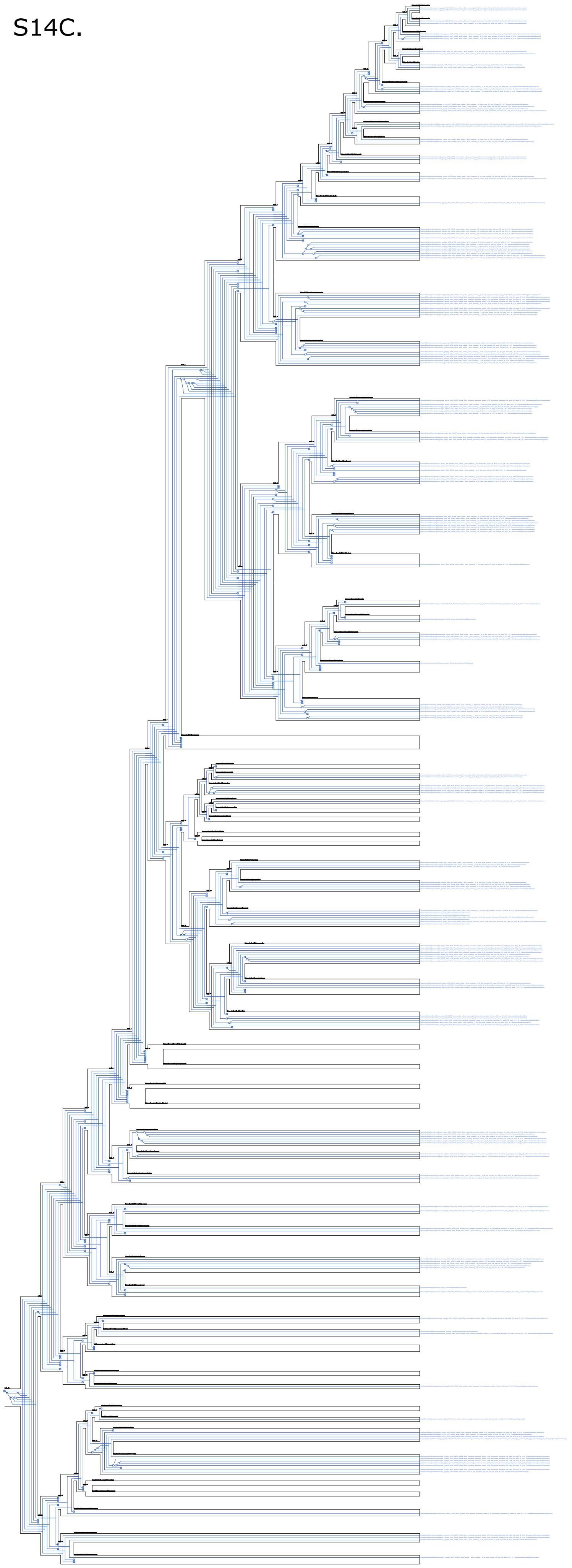

S14D

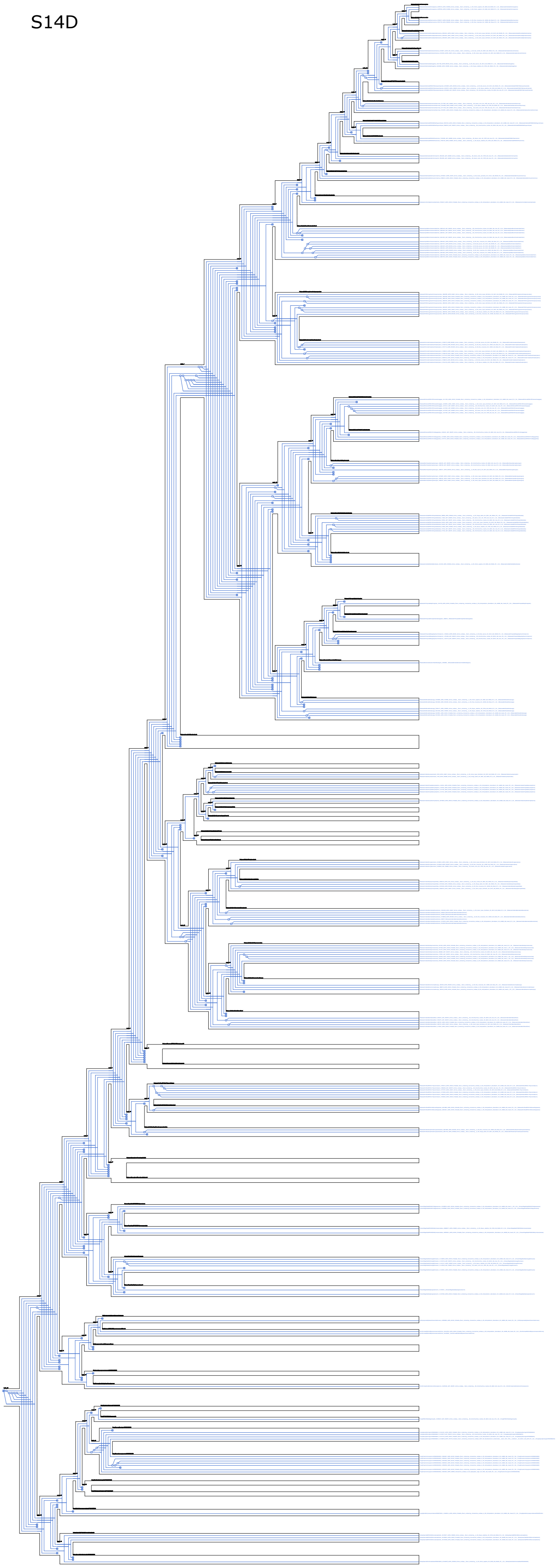

S15A.

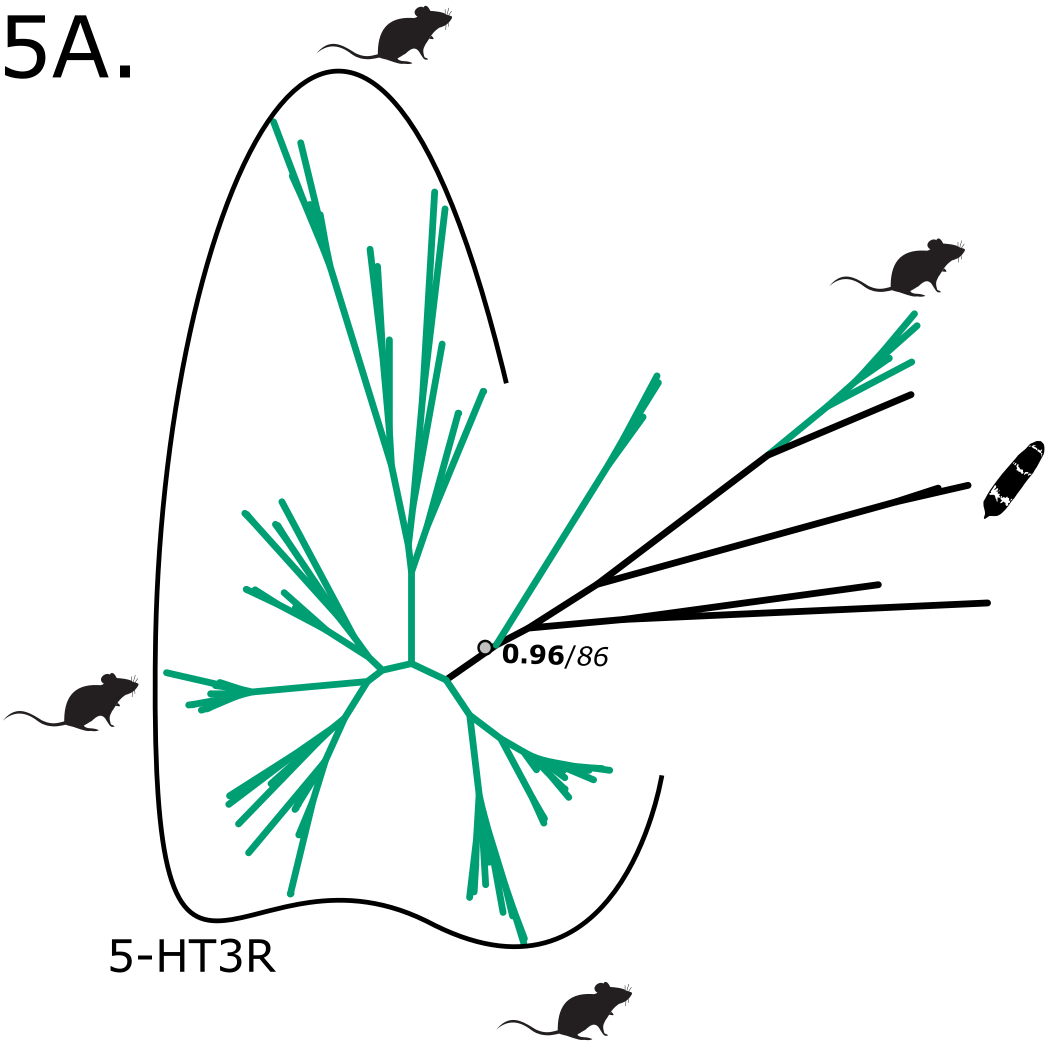

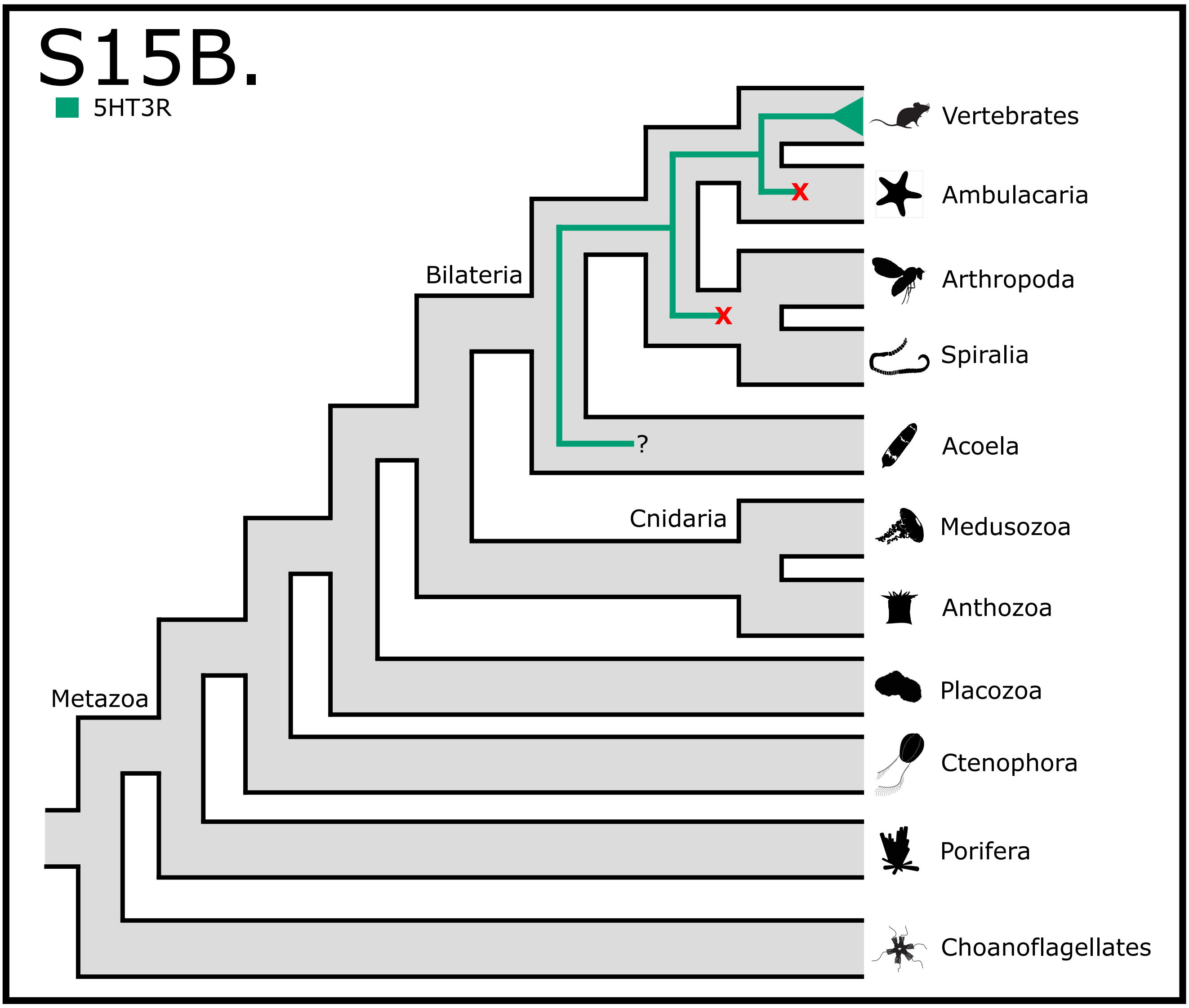
S15B.
5HT3R
Vertebrates
Ambulacaria
Arthropoda
Spiralia
Acoela
Medusozoa
Anthozoa
Placozoa
Ctenophora
Porifera
Choanoflagellates
Bilateria
Cnidaria
Metazoa
?

S16A.

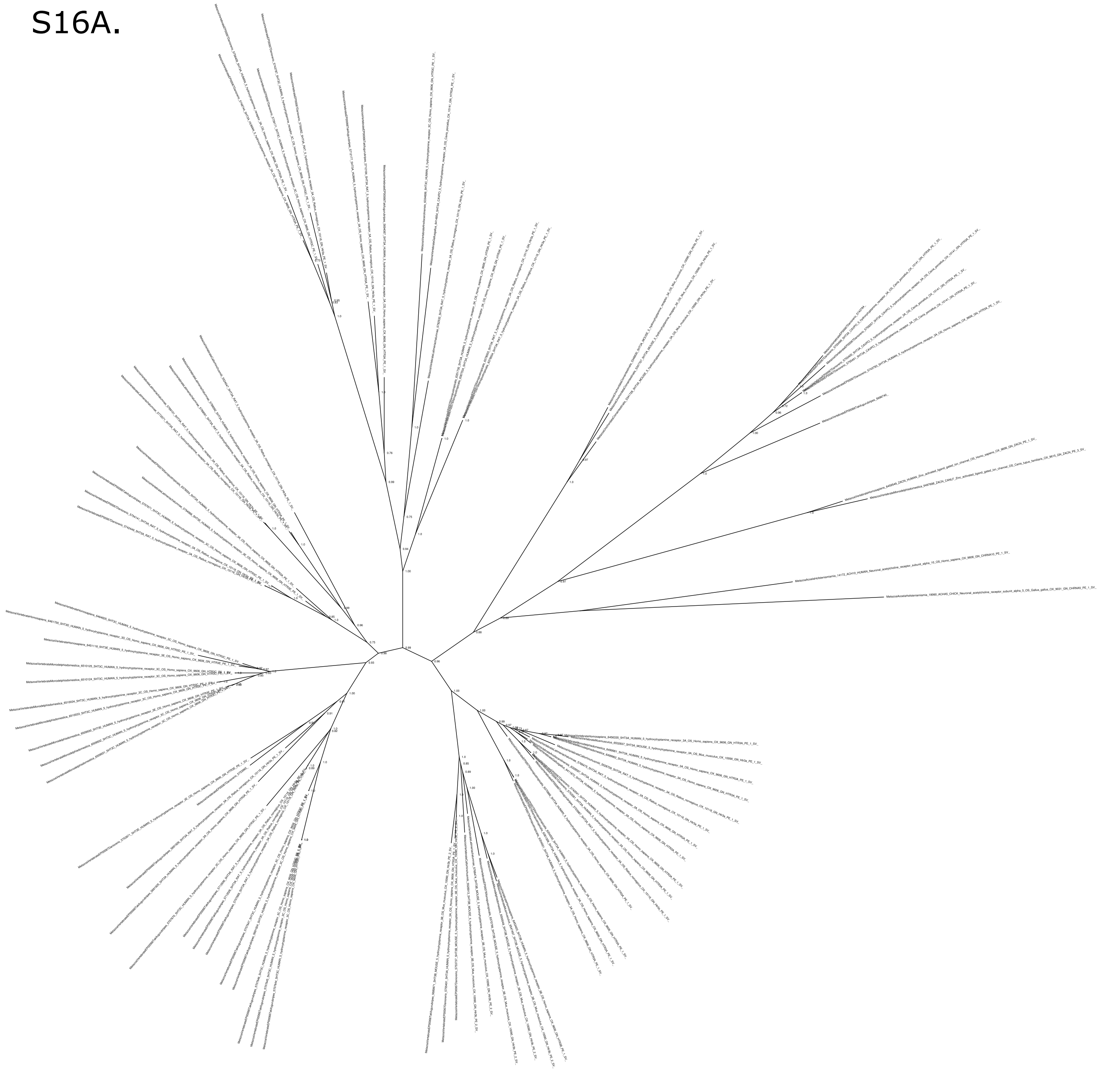

S16B.

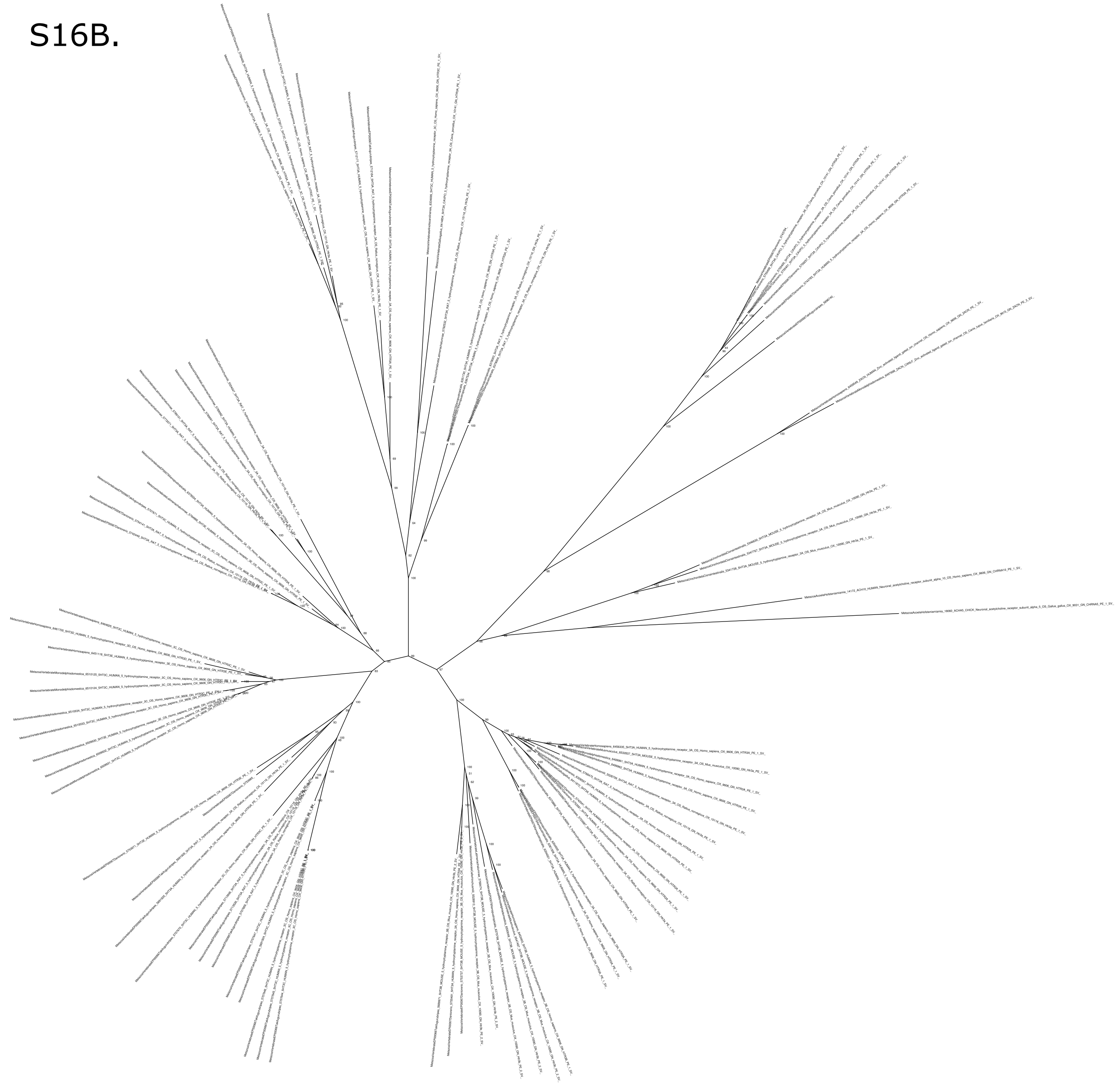

S16C

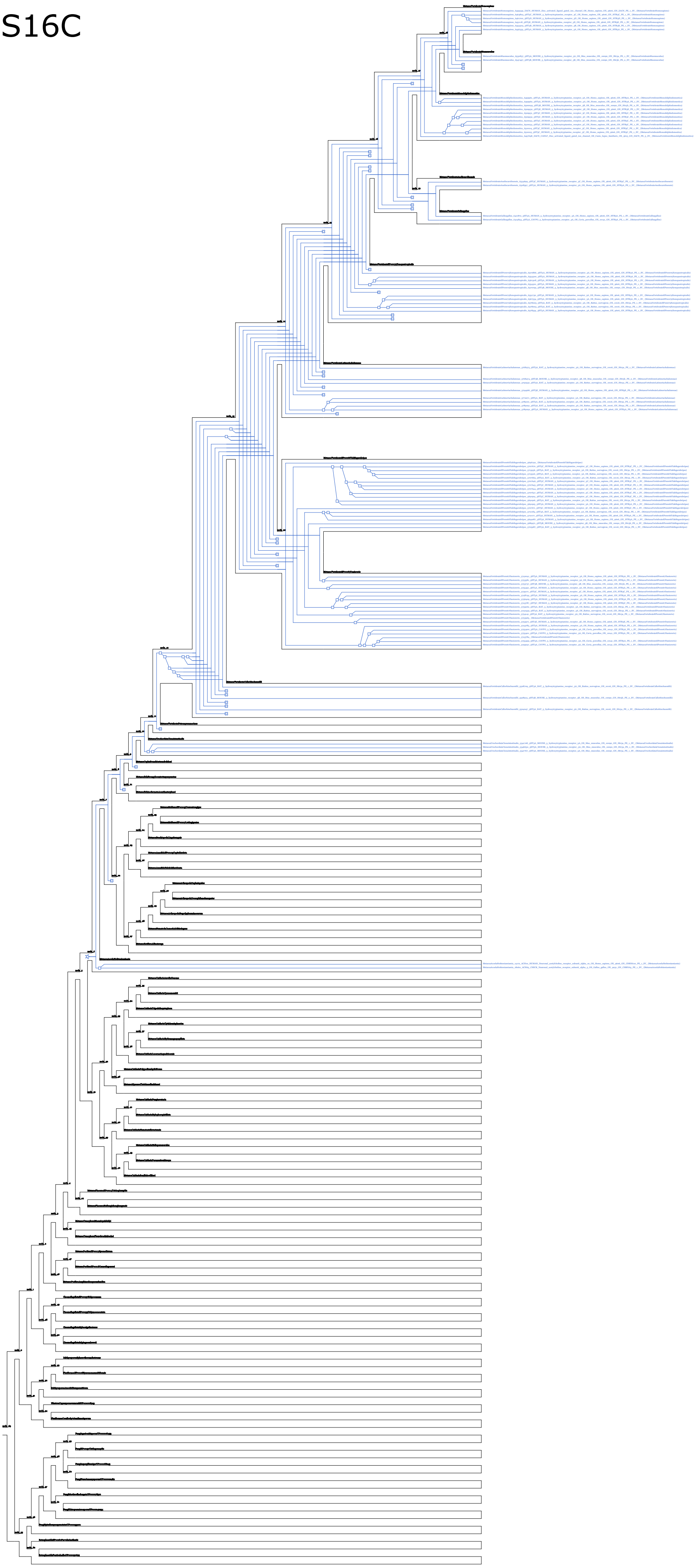

S16D

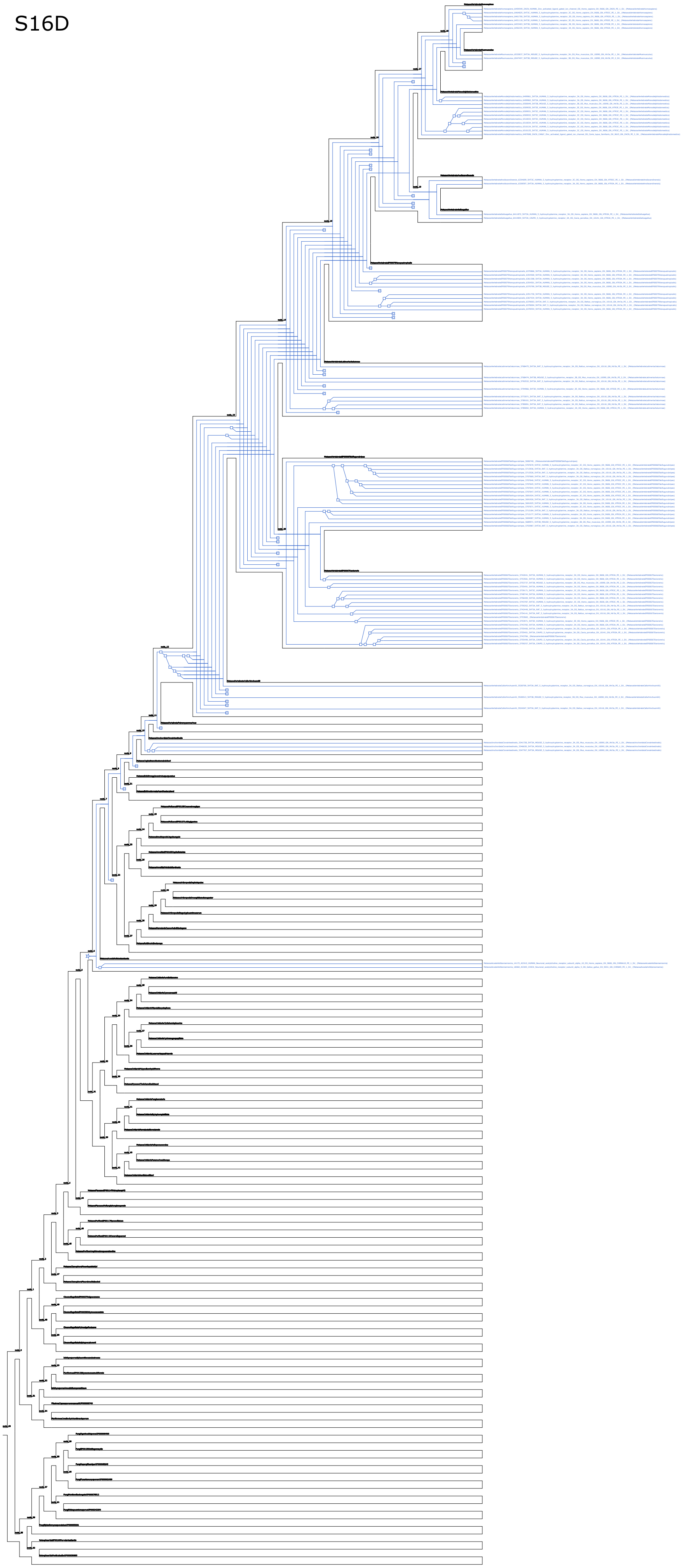

S17A.

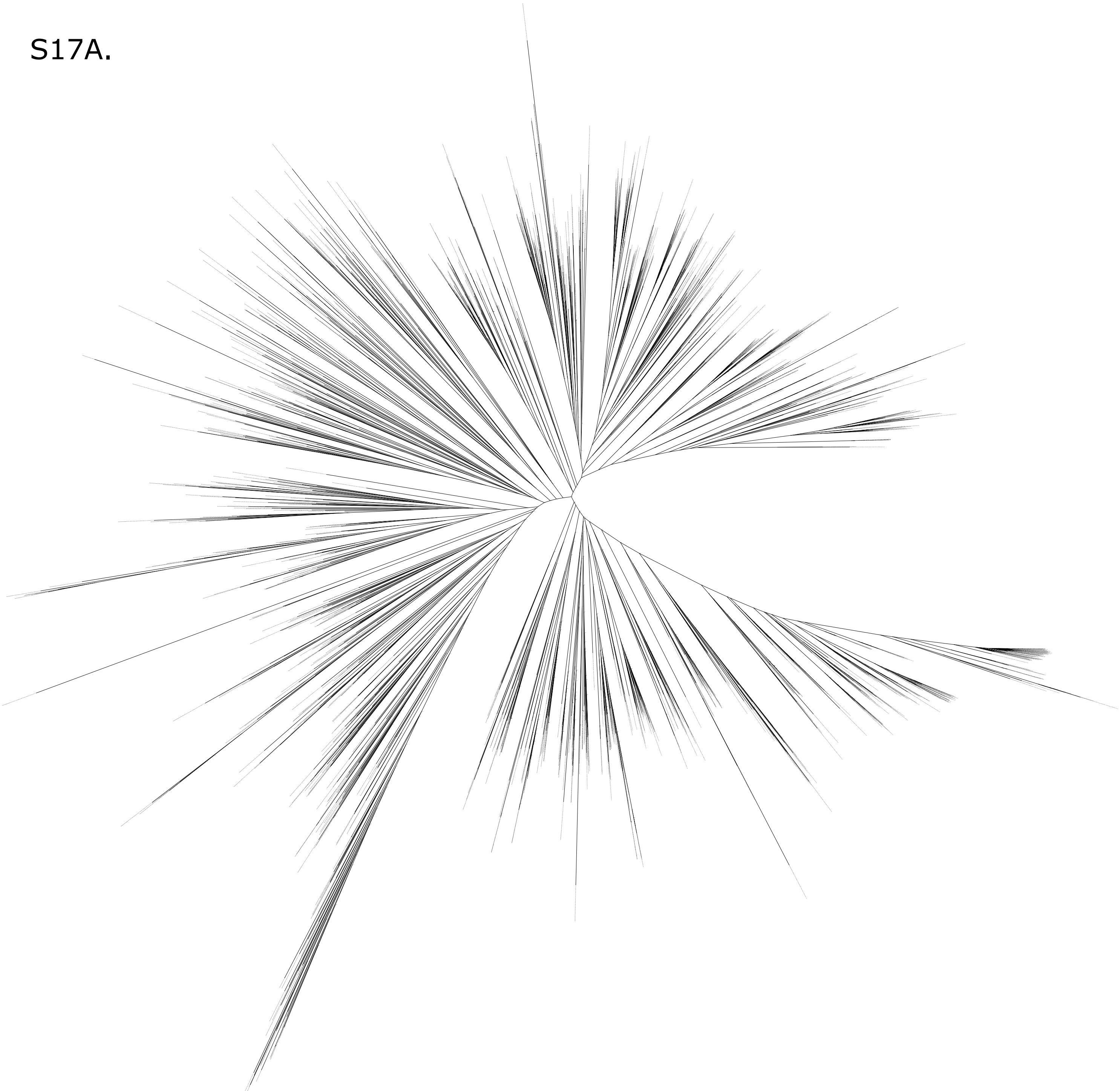

S17B.

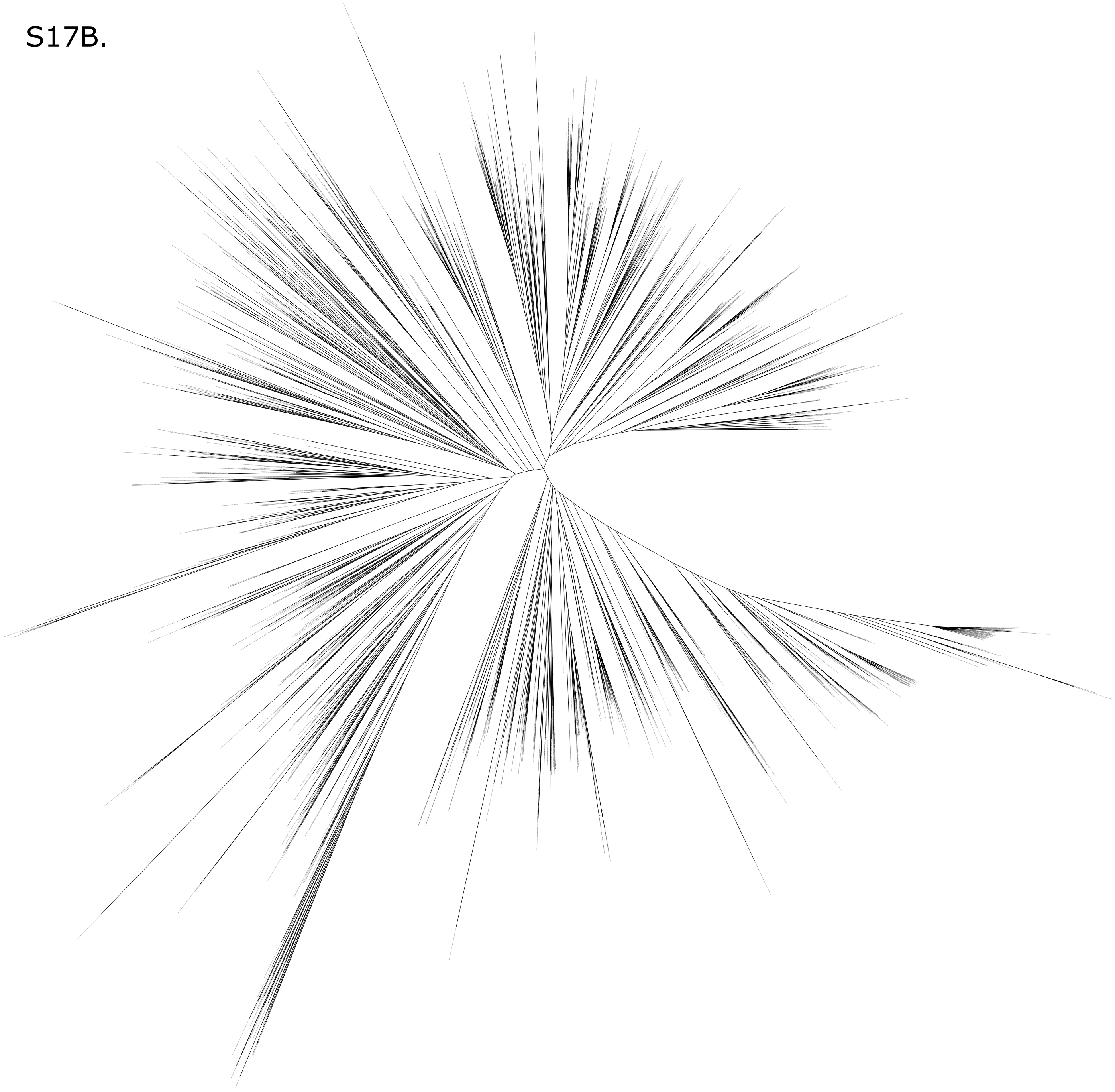

S17C

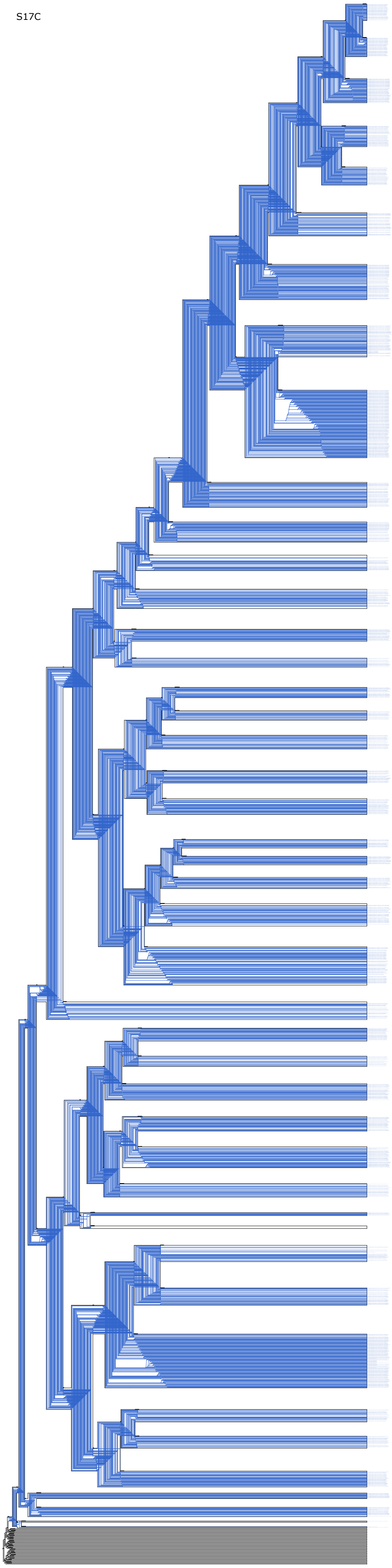

S17D

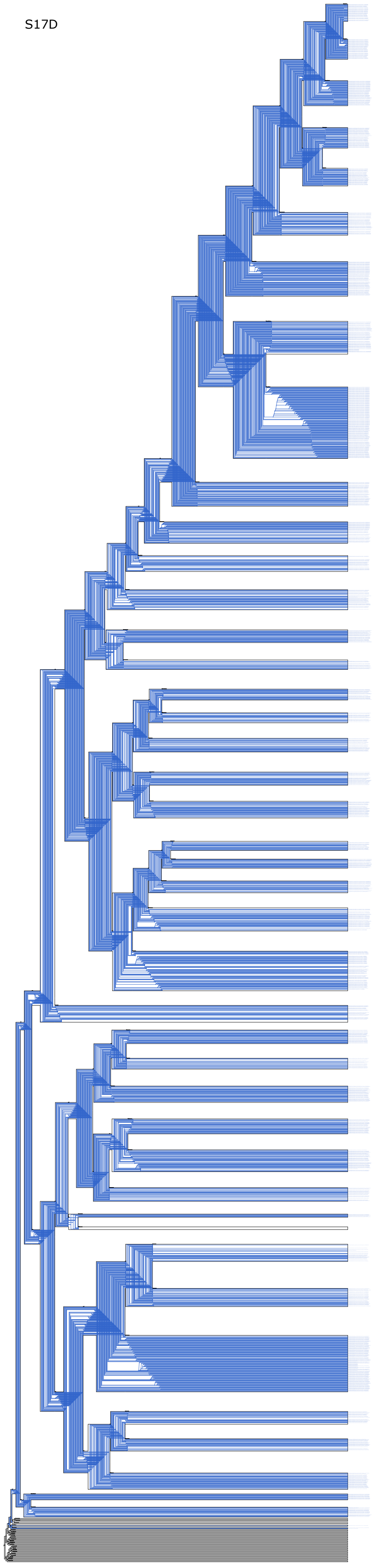

S18A.

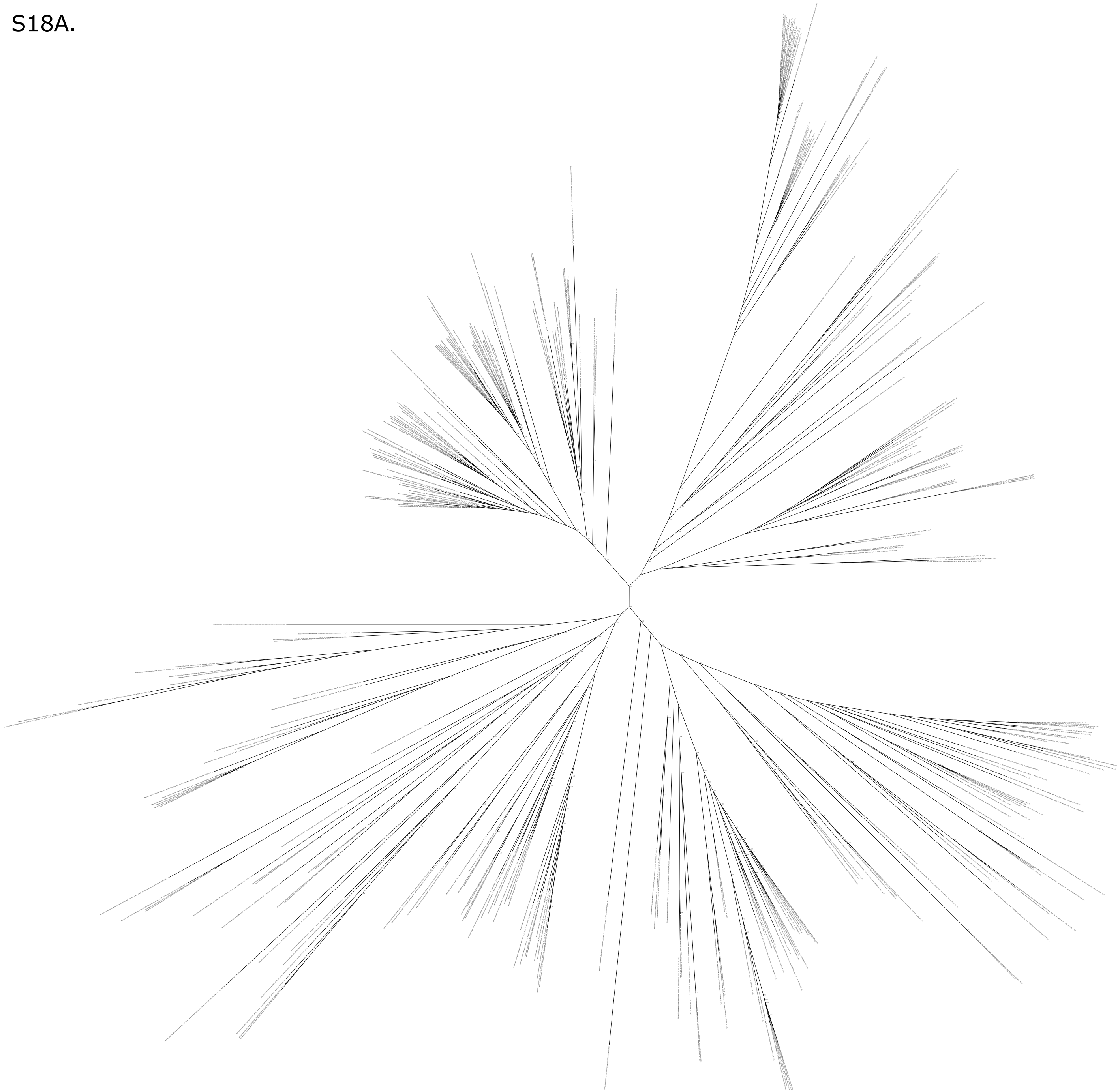

S18B.

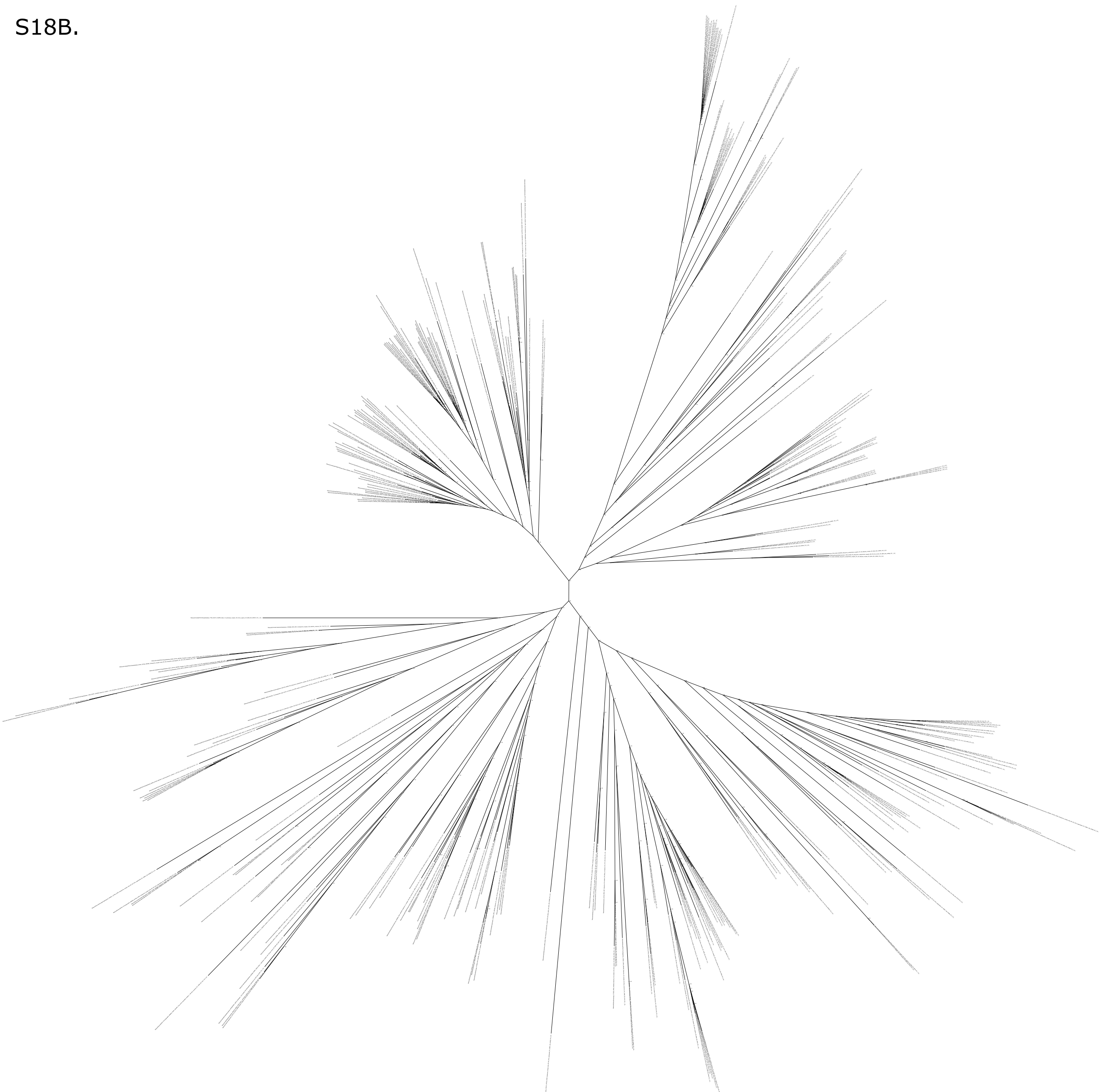

S18C

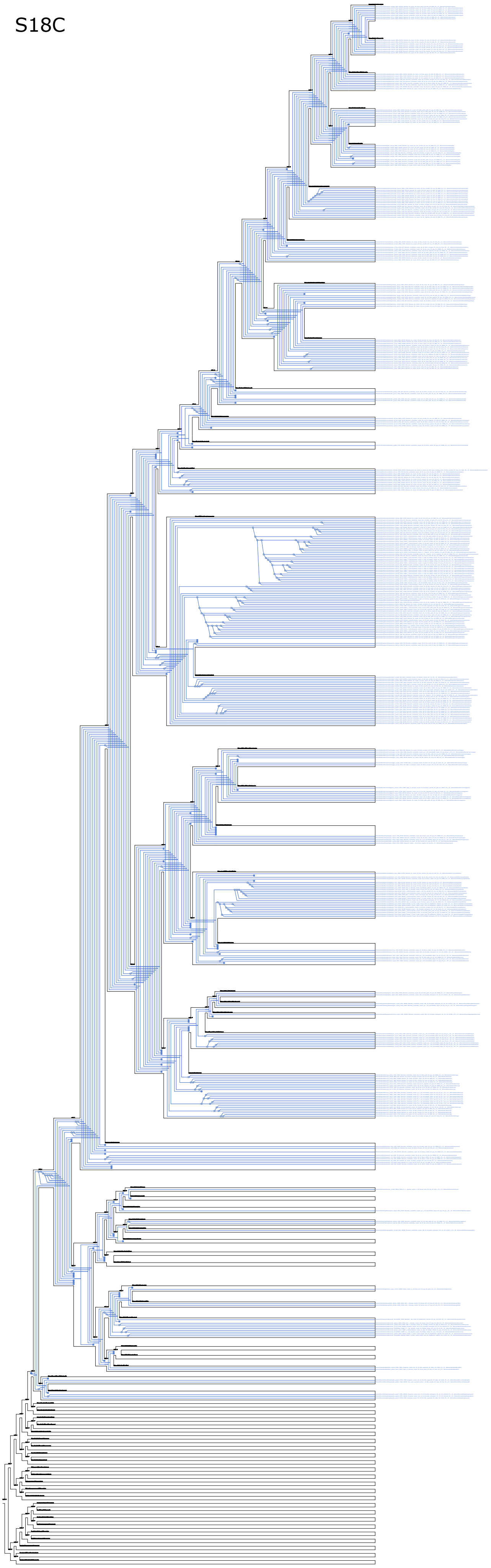

S18D

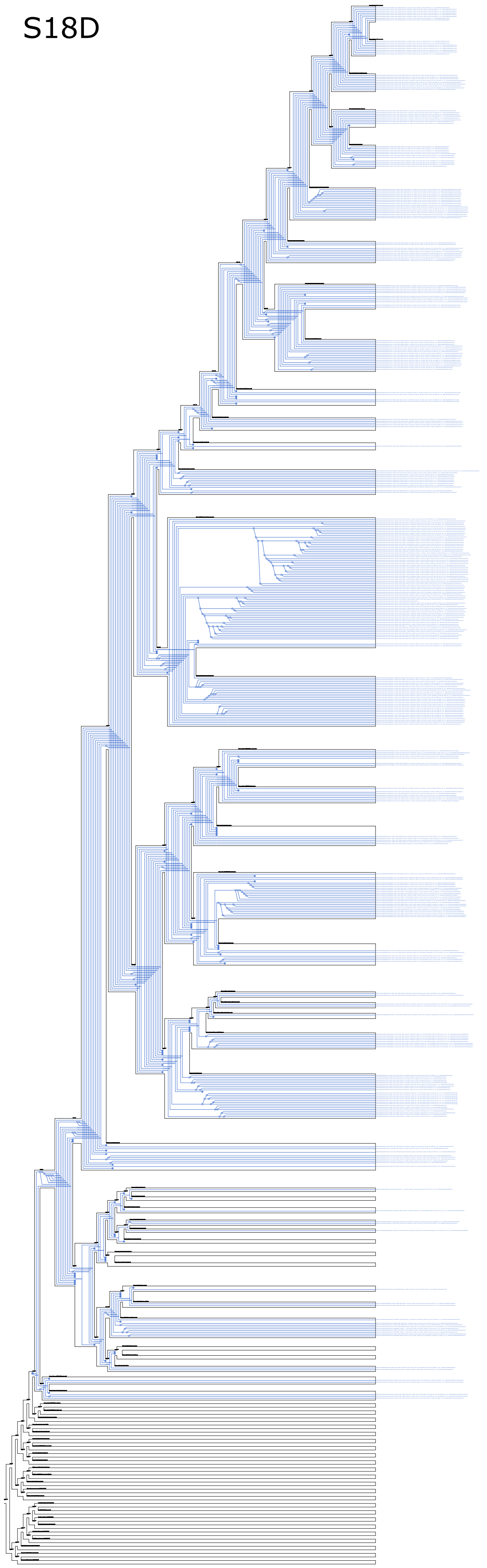

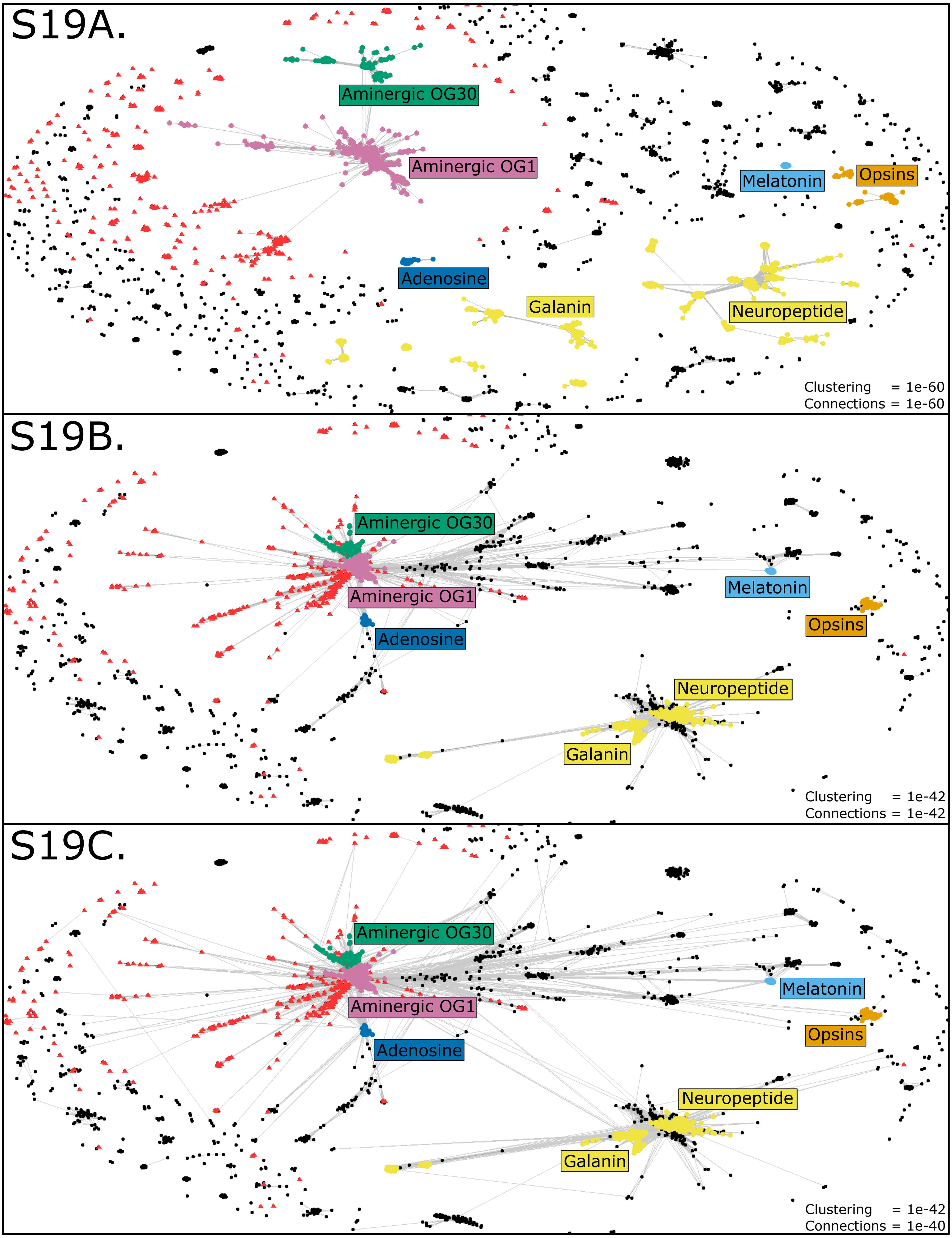
S19A.
Aminergic OG30
Aminergic OG1
Melatonin
Opsins
Adenosine
Galanin
Neuropeptide
Clustering = 1e-60
Connections = 1e-60
S19B.
Aminergic OG30
Aminergic OG1
Melatonin
Opsins
Adenosine
Neuropeptide
Galanin
Clustering = 1e-42
Connections = 1e-42
S19C.
Aminergic OG30
Aminergic OG1
Melatonin
Opsins
Adenosine
Neuropeptide
Galanin
Clustering = 1e-42
Connections = 1e-40

S20A.

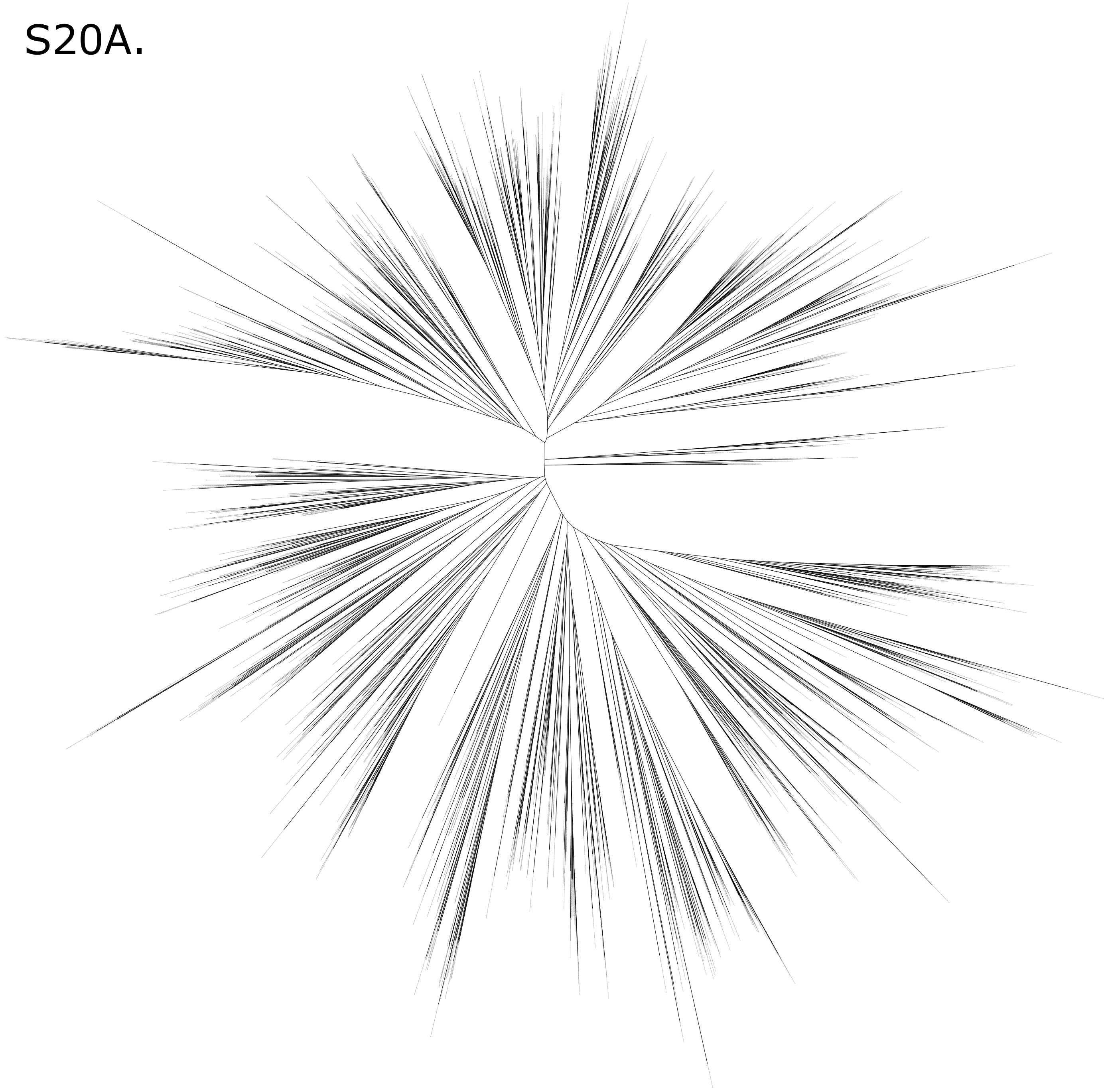

S20B.

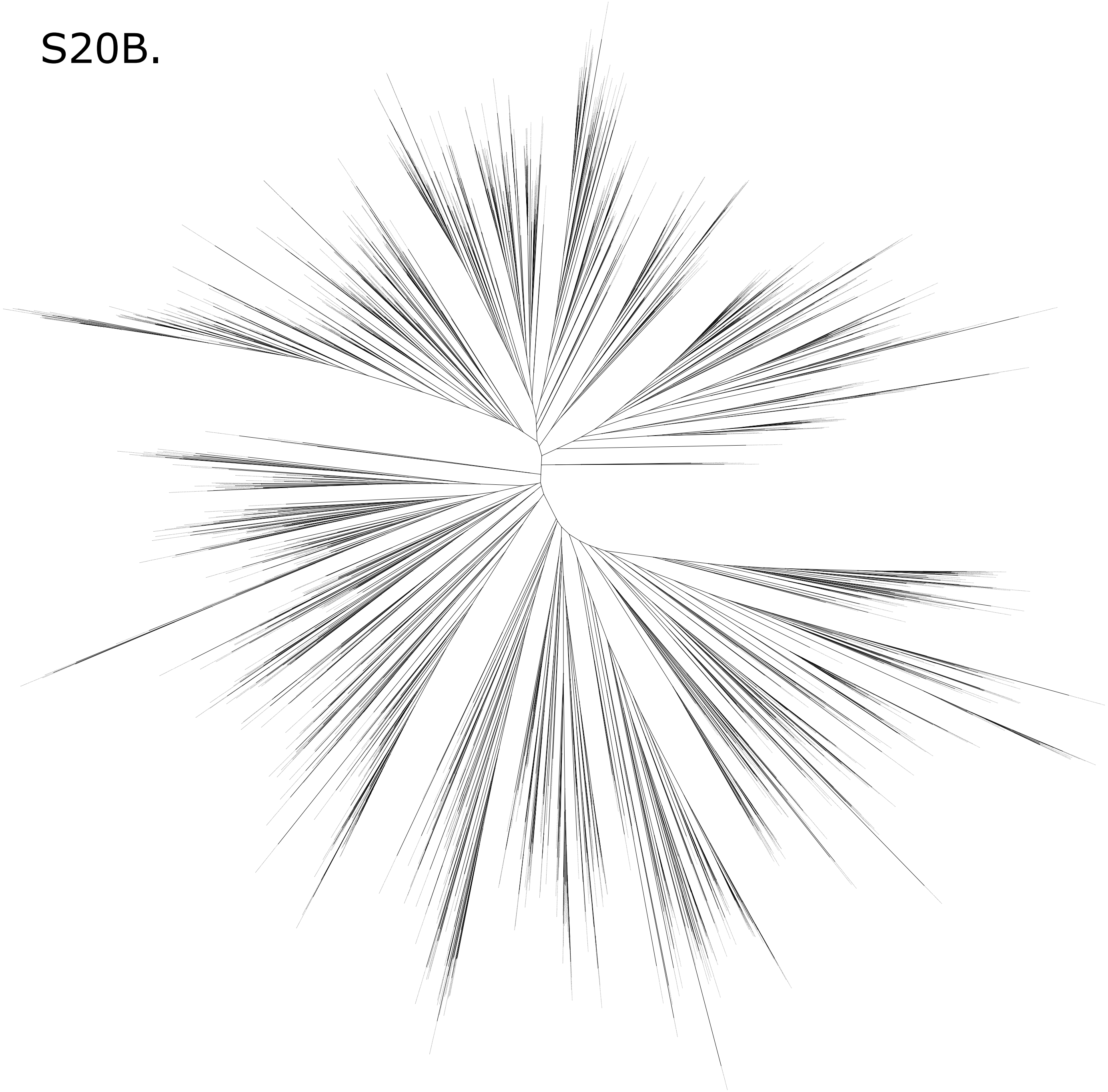

S20C

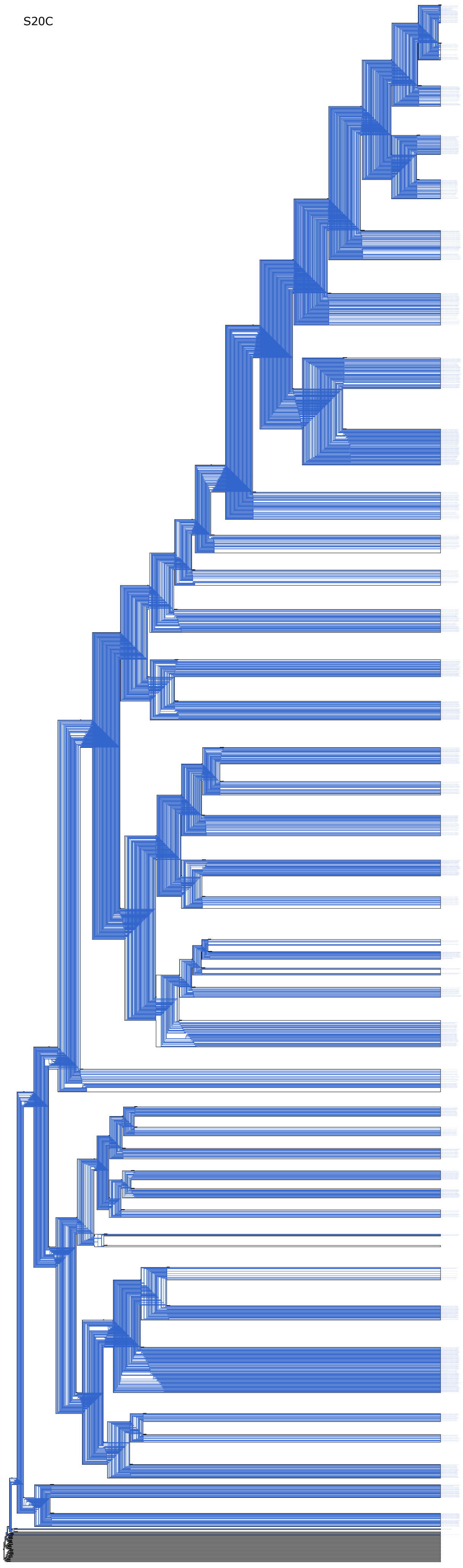

S20D

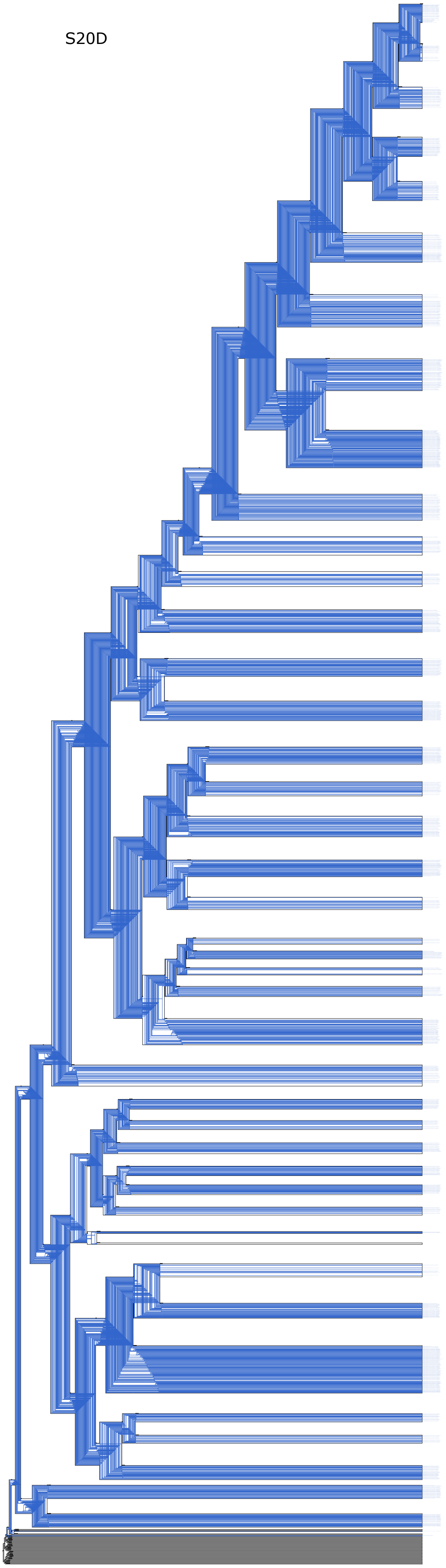

S20E

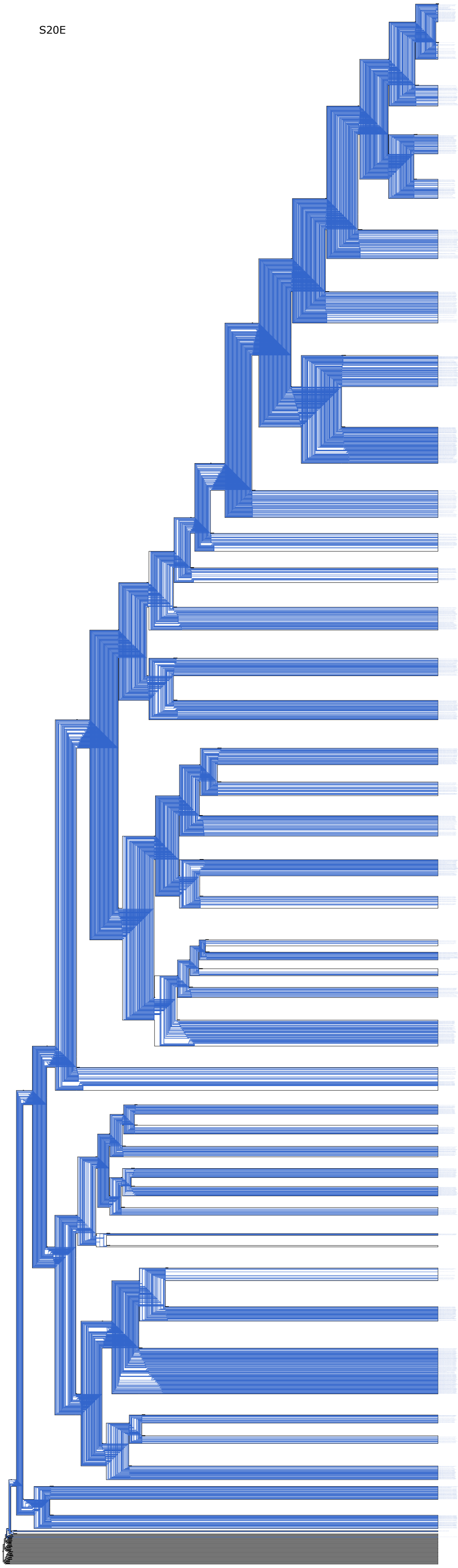

S20F

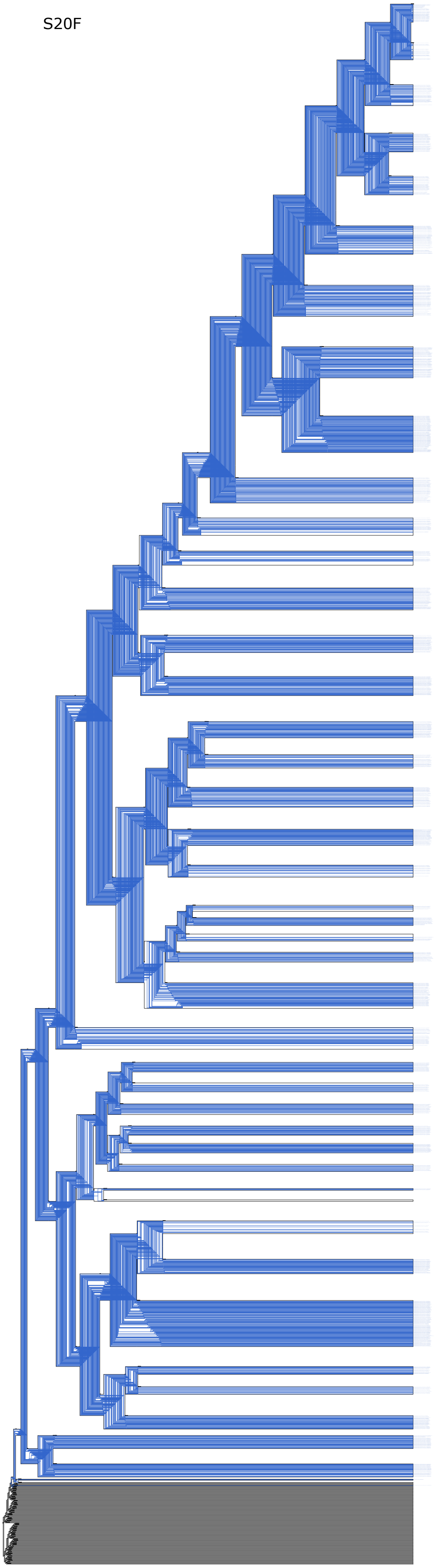
